# Prevention of ribozyme catalysis through cDNA synthesis enables accurate RT-qPCR measurements of context-dependent ribozyme activity

**DOI:** 10.1101/2024.07.19.604288

**Authors:** Nina Y. Alperovich, Olga B. Vasilyeva, Samuel W. Schaffter

## Abstract

Self-cleaving ribozymes are important tools in synthetic biology, biomanufacturing, and nucleic acid therapeutics. These broad applications deploy ribozymes in many genetic and environmental contexts, which can influence activity. Thus, accurate measurements of ribozyme activity across diverse contexts are crucial for validating new ribozyme sequences and ribozyme-based biotechnologies. Ribozyme activity measurements that rely on RNA extraction, such as RNA sequencing or reverse transcription-quantitative polymerase chain reaction (RT-qPCR), are generalizable to most applications and have high sensitivity. However, the activity measurement is indirect, taking place after RNA is isolated from the environment of interest and copied to DNA. So these measurements may not accurately reflect the activity in the original context. Here we develop and validate an RT-qPCR method for measuring context-dependent ribozyme activity using a set of self-cleaving RNAs for which context-dependent ribozyme cleavage is known *in vitro*. We find that RNA extraction and reverse transcription conditions can induce substantial ribozyme cleavage resulting in incorrect activity measurements with RT-qPCR. To restore the accuracy of the RT-qPCR measurements, we introduce an oligonucleotide into the sample preparation workflow that inhibits ribozyme activity. We then apply our method to measure ribozyme cleavage of RNAs produced in *Escherichia coli* (*E. coli*). These results have broad implications for many ribozyme measurements and technologies.

## INTRODUCTION

Self-cleaving ribozymes are ubiquitous throughout the tree of life (1, 2) and play crucial roles in viral replication and regulation of gene expression (3, 4). Ribozymes are also commonly repurposed for applications in synthetic biology (5–7), biomanufacturing (8–10), and nucleic acid therapeutics (11, 12). For example, self-cleaving ribozymes have been used to produce RNA transcripts of uniform length (8, 10), build biosensors that control gene expression (6), insulate gene expression levels from upstream sequences in genetic circuits (5), and produce multi-stranded RNA components that fold cotranscriptionally (13–15). These applications span *in vitro* transcription (IVT), cell-free lysates, and prokaryotic and eukaryotic cells, with each environment presenting unique conditions that could influence ribozyme activity (16–19). Further, ribozymes are often used in many different genetic contexts, *i.e.,* with different sequences flanking the ribozyme, and each new context has the potential to introduce alternative folds that change activity (20–25). Together, these differences in ribozyme activity across genetic and environmental contexts represent context-dependent effects (Figure 1A) that need to be measured when developing new ribozyme-based technologies or discovering new ribozyme sequences.

**Figure 1:**
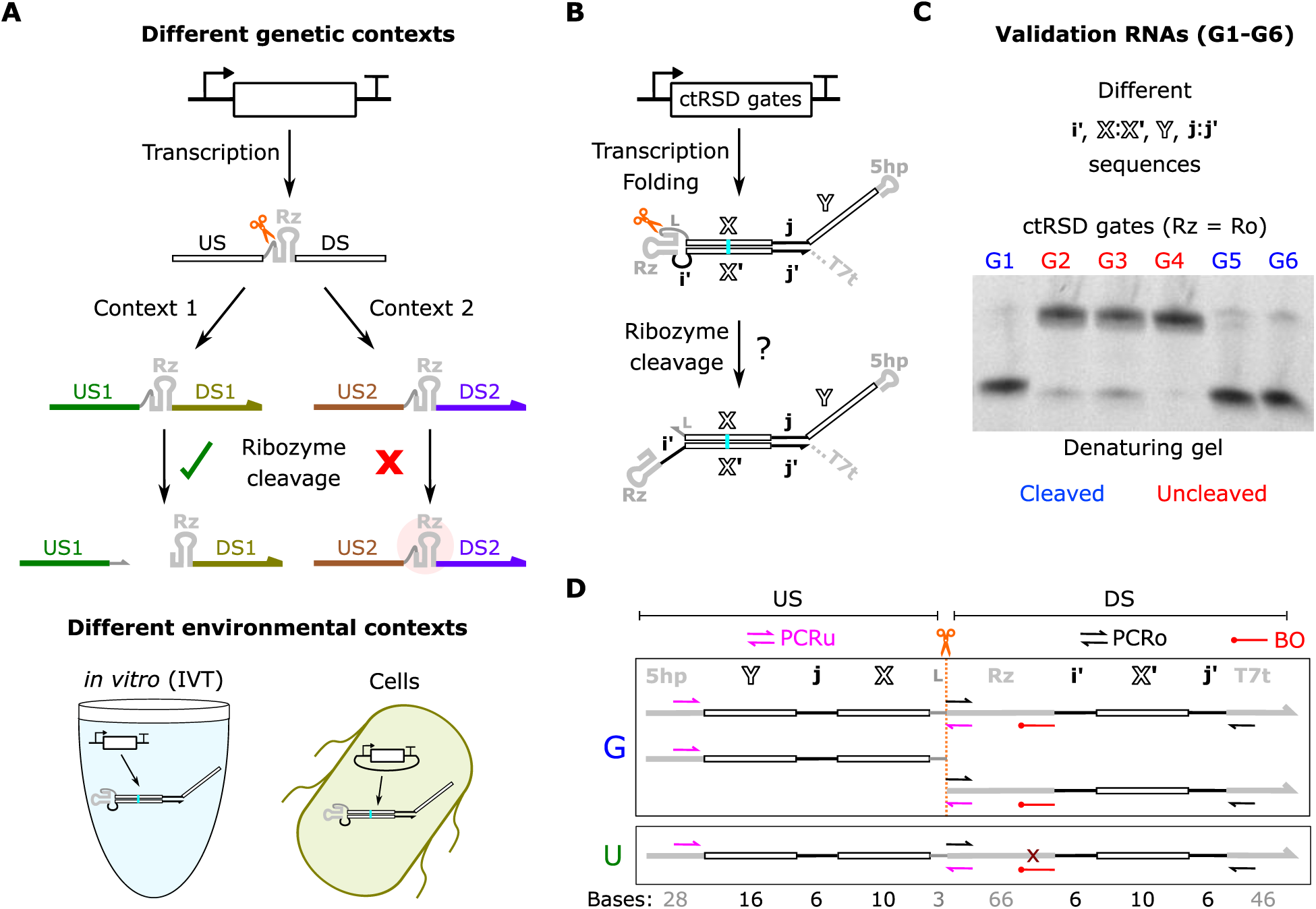
Overview of context-dependent ribozyme cleavage and RT-qPCR measurements. (**A**) Context-dependent ribozyme (Rz) cleavage. Left: Different genetic contexts *i.e.*, upstream (US) or downstream (DS) sequences, can inhibit ribozyme cleavage. Orange scissors indicate the intended ribozyme cleavage site. Right: Ribozyme activity can vary in different environments such as *in vitro* (IVT) and in cells. (**B**) Schematic of cotranscriptionally encoded RNA strand displacement (ctRSD) gates. Domains Y, j, X and domains i′, X′, j′ represent different upstream and downstream sequences, respectively. (**C**) Denaturing gel electrophoresis results of ctRSD gate sequences selected to validate the RT-qPCR method. G1 to G6 have different sequences upstream and/or downstream of the ribozyme. See Supplementary Section 1 for gate schematics and sequences. (**D**) Schematic of primer layout to measure ribozyme cleavage with RT-qPCR. The PCRu primers (pink) span the cleavage site (orange dashed line) and should only amplify uncleaved products. The PCRo primers amplify both cleaved and uncleaved products to amplify total RNA. U is a reference RNA which contains a single base mutation (maroon X) compared to G, which abolishes ribozyme activity. The difference in PCRu Ct between G and U transcripts, corrected for total RNA differences by PCRo, yields a measure of ribozyme cleavage (Methods). BO represents a blocking oligonucleotide that is added to prevent ribozyme cleavage during sample preparation (Methods). The red circle at the left end of BO indicates a 3’ amino modification to prevent extension during RT-qPCR. The numbers below the box indicate the length of each domain in bases.

Many methods for measuring self-cleaving ribozyme activity exist, such as electrophoretic separation (26–28), Förster resonance energy transfer (FRET) (29–31), connecting ribozyme activity to the expression of a reporter gene (32–37), RNA sequencing (RNA-seq) (32, 38–43), and reverse-transcription-polymerase chain reaction (RT-PCR)-based techniques (5, 44–46). Gel electrophoresis and FRET are well suited for IVT studies, but these techniques can be difficult for RNA produced in cells. Particularly for RNAs with low expression, for which signal may be below the detection limit (47). Connecting ribozyme activity to the expression of a reporter gene is ideal for *in situ* measurements in cells but can be difficult to generalize across cell lines, typically requiring different implementations for prokaryotic (33, 36, 37) and eukaryotic systems (32, 34, 35). These methods are also not applicable outside of cells as they rely on cellular machinery not usually present in many cell-free systems. RNA-seq and RT-qPCR methods have high sensitivities and are applicable to any environment because RNA is extracted prior to measurement, making these techniques ideal for evaluation of broad, context-dependent effects on ribozyme activity.

When considering RNA-seq and RT-qPCR, the choice of method often comes down to the necessary measurement scale. RNA-seq is often used for high throughput measurements, such as screening large libraries of candidate ribozyme sequences or genetic contexts, as sequencing allows for a single multiplexed measurement of the entire library (32, 38–42). RT-qPCR often makes sense for measuring a small set of sequences, *e.g.,* < 20, as it is less costly than sequencing, requires less sample preparation, and can be performed and analyzed quickly (44, 45). For both RNA-seq and RT-qPCR, RNA must be extracted from the environment in which it was produced and then reverse transcribed before a measurement takes place. These unavoidable manipulations could perturb RNA structure, potentially influencing the final ribozyme activity measurement. This is especially important when measuring context-dependent ribozyme cleavage, as RNAs that do not cleave in the context of interest may be metastable and cleave during sample preparation. For example, reverse transcription is usually performed at (40 to 60) °C to reduce RNA secondary structure, but these elevated temperatures could induce unwanted ribozyme activity in RNAs that were not active when transcribed at biological temperatures of (25 to 37) °C. Further, RNA extraction and reverse transcription buffers have different ion compositions compared to the environments the RNAs were produced in, which could alter secondary structure and catalytic activity (17). So, indirect measurements of ribozyme cleavage that rely on RNA extraction and reverse transcription may not reflect ribozyme activity in the original context.

Here, we develop an RT-qPCR method for measuring self-cleaving ribozyme activity in different contexts (Figure 1B) and evaluate the validity of the method with RNAs produced by IVT for which we have direct orthogonal measurements of cleavage (Figure 1C,D). Importantly, we find that RNA extraction and reverse transcription conditions can induce RNA sequences with ribozymes that did not cleave during IVT to self-cleave, resulting in incorrect ribozyme cleavage measurements with RT-qPCR. To circumvent this issue, we develop a workflow that prevents ribozyme cleavage during extraction and reverse transcription (Figure 2A), which enables accurate ribozyme activity measurements with RT-qPCR. We validate our method on two HDV-like ribozymes (48) in eight genetic contexts, four of which do not cleave during IVT. We then apply the method to RNAs produced in *Escherichia coli* (*E. coli*) and identify context-dependent differences in activity compared to IVT. Our findings that sample preparation influences ribozyme activity measurements have broad implications for other ribozyme measurement techniques, particularly those that involve reverse transcription (39, 45). These results are also important for characterization of DNAzymes (49), riboswitches (6, 50), and ribozymes that perform reactions other than self-cleavage (51–55). Together these results provide a straightforward and accurate method for measuring ribozyme activity that should generalize to different ribozymes and riboswitches in diverse genetic contexts and transcription environments.

**Figure 2:**
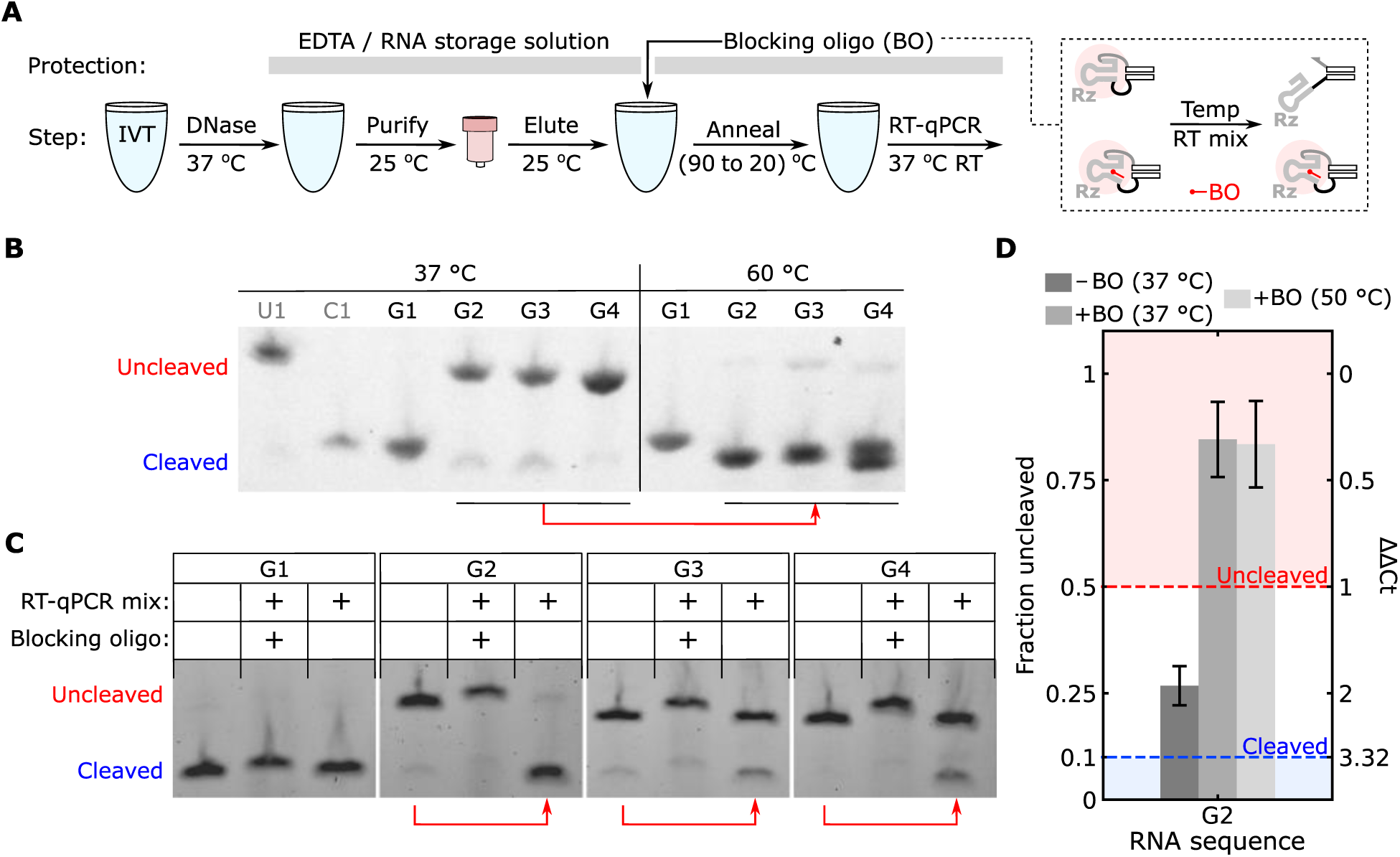
Identifying and preventing ribozyme cleavage during RNA purification and reverse transcription. (**A**) The workflow for preparing IVT RNA for RT-qPCR cleavage measurements. After DNA digestion, RNA is purified with buffers supplemented with EDTA to prevent cleavage (Methods). After RNA purification, a blocking oligonucleotide (BO) that is partially complementary to the ribozyme is added and annealed prior to RT-qPCR. The blocking oligonucleotide is designed to hybridize to the ribozyme and prevent it from adopting an active fold (right inset). (**B**) Denaturing gel electrophoresis results for RNAs after a 15 min incubation at 37 °C or 60 °C. Prior to this incubation, RNA was prepared by 30 min IVT at 37 °C followed by a 30 min DNase I digestion at 37 °C. U1 and C1 are uncleaved and cleaved control RNAs for G1, respectively. (**C**) Denaturing gel electrophoresis results for RNAs after a 10 min incubation at 37 °C in RT-qPCR reaction mixture with or without BO. Prior to incubation, the RNAs were purified from IVT reactions and annealed with or without blocking oligonucleotide (Methods). In (B) and (C), red arrows below the gel indicate undesired cleavage. G1 and G2 samples were on one gel and G3 and G4 samples were on another gel. (**D**) RT-qPCR measurements of ribozyme cleavage for G2 RNA with and without blocking oligonucleotide with reverse transcription conducted at 37 °C or 50 °C. The ΔΔCt (Eq. 2, Methods) and corresponding fraction uncleaved are labeled on the right and left axes, respectively. Error bars indicate standard deviation from three technical replicates.

## MATERIALS AND METHODS

### DNA and materials

DNA transcription templates were ordered as eBlock gene fragments from Integrated DNA Technologies (IDT). eBlocks arrived in 96-well plates eluted to 10 ng/µL in Buffer IDTE, pH 8.0. To meet the length requirements for ordering eBlock DNA, flanking sequences were appended adjacent to the sequence of the ctRSD gate amplified with PCR (See Supplementary Methods). DNA primers for PCR of transcription templates and RT-qPCR experiments were ordered from IDT without purification (standard desalting). The blocking oligonucleotides were ordered from IDT without purification (standard desalting) and a 3’ amino modification to prevent extension. The pET-Duet-1 plasmid, used as the backbone for RNA expression in *E. coli*, was ordered from Millipore Sigma (71146-3).

All DNA sequences from this study are available in Supplementary File S1. Annotated DNA and RNA sequences are in Supplementary Section 1. Reagents used for specific techniques are specified in the relevant methods below. A full list of materials with vendors and catalogue numbers is available in the Supplementary Methods.

### PCR amplification of DNA

PCRs were conducted with 2x Phusion High-Fidelity Master Mix (ThermoFisher, F531L) and 0.5 µmol/L of each DNA primer. Amplification of linear DNA templates for *in vitro* transcription reactions were conducted with 0.02 ng/L of eBlock DNA and T7fwd and T7rev primers (Supplementary Table S2) with 30 cycles consisting of a 30 s denaturing step at 98 °C, a 30 s primer annealing step at 60 °C, and a 30 s extension step at 72 °C. A 3 min extension step at 72 °C was executed at the end of the program. The same protocol was followed for preparation of linear DNA inserts for cloning into plasmids but with ga4_fwd and pET_ds_rev primers (Supplementary File S1). Amplification of the pETDuet-1 backbone was conducted with 0.2 ng/uL of plasmid DNA and the pET_BB_ds_fwd and pET_BB_rev_ga4 primers (Supplementary File S1) with 30 cycles consisting of 30 s denaturing step at 98 °C, a 30 s primer annealing step at 62 °C, and a 2 min extension step at 72 °C. A 5 min extension step at 72 °C was executed at the end of the program. Following PCR, samples were purified with a QIAquick PCR purification kit (Qiagen, 28104), eluted in Qiagen Buffer EB (10 mmol/L tris-HCl, pH 8.5), and measured with absorbance at 260 nm on a DeNovix D-11 Series Spectrophotometer.

### Gel electrophoresis

All RNA gel electrophoresis experiments were conducted with 4 % agarose EX E-gels. These gels come prestained with SYBR Gold for fluorescence imaging. Electrophoresis was conducted on an E-gel powerbase (ThermoFisher, G8200), and E-gels were imaged using a FAS-Digi PRO system equipped with a blue-green light source (470 nm to 520 nm) and Canon 250D camera (Nippon Genetics). For denaturing gels, a solution of 100 % formamide and 36 mmol/L ethylenediaminetetraacetic acid (EDTA) was mixed 1:1 by volume with the samples and the samples were heated to 85 °C for 5 min before electrophoresis. The samples were immediately loaded on gels and run for 30 min before imaging. Gel images were not postprocessed; any brightness and contrast adjustments were applied uniformly across each image during image acquisition. Unless otherwise stated, white spaces between gel images represent images taken from different gels. Each gel had size markers, the uncleaved (U) and cleaved (C) controls, which were used to align gel images.

### RNA production and purification from IVT

To prepare IVT RNA for RT-qPCR, 25 nmol/L of DNA template was transcribed at 37 °C for 1 h in transcription buffer prepared in house (40 mmol/L tris-HCl (pH 7.9), 6 mmol/L MgCl2, 10 mmol/L dithiothreitol, 10 mmol/L NaCl, and 2 mmol/L spermidine) supplemented with 1 U/µL of T7 RNAP (ThermoFisher, EP0113) and with 2 mmol/L of each NTP (ThermoFisher, R0481). These conditions were used for all DNA templates other than G2h and U2h, for which 100 nmol/L of DNA template and 10 U/µL of T7 RNAP were used. Following transcription, 3.33 U/µL of DNase I and DNase reaction buffer (ThermoFisher, EN0523) were added, and samples were incubated at 37 °C for 30 min. RNA was then purified with an RNA clean and concentrate kit (ZymoResearch, R1016) with the RNA binding buffer supplemented with 36 mmol/L of EDTA and without the prep buffer step. RNA was eluted in TE buffer (ThermoFisher, AM9849) and immediately mixed with an equal volume of RNA storage solution (1 mM sodium citrate, pH 6.5) (ThermoFisher, AM7001). RNA concentrations were determined with a Qubit using a high-sensitivity RNA quantification kit (ThermoFisher, Q32852). If not used immediately, RNA samples were stored at – 80 °C. A step-by-step RNA extraction protocol is in the Supplementary Methods.

### Plasmid assembly and cloning

Plasmids were assembled with Gibson Assembly (56) using 30 base homology domains. After PCR and clean up, the plasmid backbone was subsequently digested with a volume fraction of 4.3 % FastDigest DPNI (ThermoFisher, FD1703) for 1 hour at 37 °C and then purified with a QIAquick PCR purification kit. Backbone and insert DNA were then mixed to the same final mass concentrations (≈15 ng/µL each) with 2x Gibson Assembly mix (New England Biolabs, E2611L) and incubated at 50 °C for 1 hour. These samples were then transformed into electrocompetent DH5α cells prepared in house (derived from ThermoFisher, 18265017) and grown overnight on Luria broth (LB, ThermoFisher, BP1426) agar plates supplemented with 100 µg/mL ampicillin (Millipore Sigma, A9518-25G). The next day colony PCRs of single colonies were conducted with sequencing primers (pET_seq_fwd and pET_seq_rev, Supplementary File S1) and PCR products with the correct insert size from gel electrophoresis were sequence verified with Sanger sequencing. Colonies with sequence verified plasmids were then grown overnight in 3 mL of LB supplemented with 100 µg/mL ampicillin. Plasmids were extracted from overnight cultures using a Qiagen Spin Miniprep kit (27104) and these plasmids were then transformed into electrocompetent BL21 Star (DE3) cells prepared in house (derived from Invitrogen, C601003). Electrotransformations were conducted with MicroPulser Electroporation Cuvettes, 0.2 cm gap (BioRad, 1652086) using an Eporator (Eppendorf, 4309000027).

### RNA production and extraction from *E. coli*

BL21 Star (DE3) cells transformed with a pET plasmid encoding for a gate sequence with a constitutive T7 promoter were grown at 37 °C overnight in 3 mL of LB supplemented with 100 µg/mL ampicillin. Overnight cultures were diluted 100-fold into 10 mL of LB supplemented with 100 µg/mL ampicillin and 100 µmol/L of isopropyl ß-D-1-thiogalactopyranoside (IPTG) and subsequently grown at 37 °C until the culture reached an optical density at 600 nm of 0.5 to 0.8 (≈6 h). Optical density measurements were conducted on a Genesys 30 visible spectrophotometer (ThermoFisher, 840-277000). One and a half mL of culture was then mixed with twice the volume of RNAprotect bacteria reagent (Qiagen, 1018380). Cells were then pelleted, resuspended in 0.1 mL of 50 mmol/L of EDTA (pH 8.0), and lysed with 1 mL of TRI reagent (ZymoResearch, R2050). Total RNA was extracted using chloroform separation and ethanol precipitation and subsequently purified using Zymo-Spin IIICG columns (ZymoResearch, C1006-50-G) and wash buffer (ZymoResearch, C1001-50). Purified RNA was eluted in nuclease free water (Ambion, AM9938) and incubated for 30 min at 37 °C with 4.17 U/µL of DNase I in DNase reaction buffer (ThermoFisher, EN0523). Samples were then purified as in “RNA production and extraction from IVT”. If not used immediately, RNA samples were stored at – 80 °C. A step-by-step protocol is in Supplementary Methods.

### Preventing ribozyme cleavage with blocking oligonucleotides

After purification, 14 µL mixtures of RNA and blocking oligonucleotides were prepared in RNA storage solution to a final concentration of 8.6 ng/µL for IVT RNA or 17.1 ng/µL for *E. coli* total RNA and 157 µmol/L of blocking oligonucleotide. These samples were then heated to 90 °C for 5 min and subsequently cooled to 20 °C at a rate of -1 °C per minute. Samples were diluted to 1 ng/µL in RNA storage solution with concentrations verified with a Qubit using a high-sensitivity RNA quantification kit (ThermoFisher, Q32852). Gate (G), uncleaved (U) control, and cleaved (C) control samples were prepared with blocking oligonucleotide prior to RT-qPCR. A step-by-step protocol is in Supplementary Methods.

### RT-qPCR experiments

RT-qPCR experiments were conducted using a SuperScript III Platinum SYBR Green One-Step Kit (ThermoFisher, 11736059) in an Applied Biosystems ViiA 7 Real-Time PCR System. RT-qPCR reactions contained a final concentration of 3 mmol/L of MgSO4, 0.2 mmol/L of each dNTP, 50 nmol/L of ROX reference dye, and 0.2 μmol/L of forward and reverse primers. A final concentration of 0.08 pg/µL or 4 pg/µL of RNA with blocking oligonucleotide was used for IVT RNA or *E. coli* total RNA, respectively. RT-qPCR experiments were initiated with a 10 min reverse transcription step. Unless otherwise stated, the reverse transcription step was conducted at 37 °C for sequences with Ro and 50 °C for sequences with Rh. Samples were then heated to 95 °C for 5 min followed by 40 cycles of a 15 s denaturing step at 95 °C and a 60 s annealing/extension step at 60 °C. Fluorescence readings were taken during the annealing/extension step of each cycle. This was followed by a 60 °C to 95 °C temperature increase for melt curve analysis.

For a given gate sequence, samples with the gate (G) RNA and samples with the corresponding uncleaved (U) control RNA were prepared with either the PCRu or PCRo primer sets (Supplementary File S1). Experiments were conducted with three technical replicates, making a minimum of 12 samples for a given sequence. Most experiments also included samples for a dilution series of the relevant uncleaved (U) control RNA to assess primer amplification efficiency, a sample with the relevant cleaved (C) control RNA, a no template control, and a no reverse transcriptase control. A plate layout with descriptions is in Supplementary Section 2.1.

From melt curve analysis and gel electrophoresis, we confirmed PCRu and PCRo primers yielded single DNA products of expected size. We also confirmed that no template and no reverse transcription controls did not have amplification within 10 cycles of our measurements (Supplementary Section 2.3).

### RT-qPCR data analysis

In each experiment the cycle threshold (Ct) value was manually adjusted in Applied Biosystems ViiA software and applied to all samples on the plate. The thresholds used for each experiment are in the uploaded data files and shown in amplification curves in Supplementary Section 2.5. Data was exported as Excel files and analyzed with custom Python code (Python 3.8.18, Spyder 5.4.3).

Using a relative quantification technique typically termed the ΔΔCt method (44, 45, 57) (Supplementary Section 2.2), the fraction of uncleaved gate for a given sequence was calculated with Eq. 1. The ΔΔCt value is the difference between the ΔCt of U and G for PCRu and the ΔCt of U and G for PCRo (Eq. 2). Because the PCRu primers only amplify uncleaved products, ΔCtPCRu indicates the concentration difference in uncleaved RNA between the U and G samples. Because the PCRo primers amplify both uncleaved and cleaved, ΔCtPCRo accounts for any differences in the total concentration of U and G RNA added in an experiment (Figure 1B).

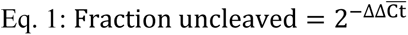

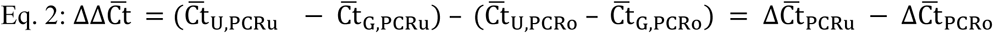

Where C̅t indicates the mean of three technical replicates.

For the above analysis to be valid, PCR amplification efficiencies should be between 90% to 110% (58). Using dilution series of U control RNAs, we verified the PCR amplification efficiencies for PCRo and PCRu primers ranged between 94 % to 106 % across sequences and replicates from independent RNA extractions (Supplementary Sections 2.4 and 2.5). Further, reverse transcription efficiency for PCRo cannot vary substantially between the G and U samples of a given sequence, otherwise this PCR cannot be used to control for differences in concentration between the samples, as the two samples would have different Ct values at the same concentration. We found the amplification curves from PCRo for U, C, and G RNAs were nearly identical for most sequences (Supplementary Sections 2.4 and 2.5). For Rh sequences, we found reverse transcription efficiencies appeared to differ for U and G RNAs at 37 °C but not at 50 °C (Supplementary Figures S8 and S10), so we used 50 °C reverse transcription for these experiments. Reverse transcription temperatures should be tested for each new ribozyme sequence or primer set.

*Error bars.* Uncertainty in Ct measurements was obtained by calculating the standard deviation of the mean Ct values from three technical replicates for each sample. These errors were then propagated to ΔCt and ΔΔCt using Eq. 3 and the final error in measured fraction uncleaved was estimated using Eq. 4 (57, 59). Step-by-step error propagation is in Supplementary Section 2.2.

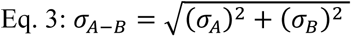

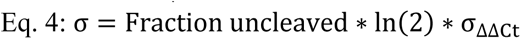

Where σ indicates standard deviation and σ*_A–B_* indicates standard deviation of ΔCt or ΔΔCt.

## RESULTS

### Overview of RT-qPCR method to measure ribozyme cleavage

Here we sought to use RT-qPCR to measure context-dependent ribozyme cleavage. To access the accuracy of such methods, we selected validation RNA sequences for which ribozyme cleavage activities *in vitro* had been previously measured using denaturing gel electrophoresis. Gel electrophoresis is an ideal orthogonal measurement for benchmarking because it does not require sample manipulation after IVT, the RNA is simply loaded on the gel for analysis. So, we take gel electrophoresis measurements as the ground truth for assessing the accuracy of RT-qPCR methods.

The validation sequences we selected are from the cotranscriptionally encoded RNA strand displacement (ctRSD) toolkit (14) (Figure 1B). ctRSD circuits are an emerging technology with potential applications spanning many environments, from cell-free biosensing (60) to cellular computation (61, 62). In ctRSD circuits, gates containing HDV-like ribozymes are designed to fold and self-cleave during transcription (Figure 1B). After ribozyme cleavage, gates can participate in strand displacement reactions programmed to process information (13), so ribozyme activity is crucial for desired performance (14). Previously, a library of ctRSD gates in different genetic contexts with both efficient and poor ribozyme cleavage in IVT have been identified (14). For our validation RNAs, we selected six ctRSD gate sequences (G1 to G6) with the same ribozyme sequence, a minimal version of the antigenomic HDV ribozyme (8) that we term Ro. Three of these gates have genetic contexts in which the Ro cleaves well (>75 % as measured by gel electrophoresis) and three have genetic contexts in which the Ro cleaves poorly (<25 % as measured by gel electrophoresis) (Figure 1C). In addition to the method validation RNAs, the ctRSD toolkit has characterized other ribozymes, such as the CPEB-3 ribozyme (48), which also allows us to evaluate generalizability.

The RT-qPCR method we employed to measure ribozyme cleavage activity is based on relative quantification methods, analogous to ΔΔCt methods, developed previously (44, 45, 57). In brief, two primer sets are used in these experiments, one set that spans the cleavage site and only amplifies uncleaved RNA (PCRu), and another set that bind downstream of the cleavage site and amplify both cleaved and uncleaved RNA (PCRo, Figure 1B). The change in cycle threshold (Ct) of the two primer sets for an RNA of interest (G) relative to a control transcript without cleavage (U) is used to determine the fraction of uncleaved RNA in a given context (Methods). For simplicity, we implemented this method using a one-step RT-qPCR reaction with SYBR Green reporting.

We designed the PCRu and PCRo primer sets for RT-qPCR, termed (Figure 1D), to bind to regions of the ctRSD gates that did not vary from gate to gate so the same primers could be used for each genetic context of our validation RNAs. PCRu and PCRo were designed with different reverse transcription primers to keep the amplicon lengths below 150 bases (63). As a reference transcript for relative quantification, we used gates with a point mutant in the ribozyme sequence that renders it incapable of cleavage (U in Figure 1D). To assess the accuracy of RT-qPCR measurements on our validation RNAs, we applied the following benchmarking metrics (Supplementary Section 3): sequences with >0.5 fraction uncleaved are considered poor cleavers and sequences with <0.1 fraction uncleaved are considered good cleavers. From the gel electrophoresis results (Figure 1C), the six validation RNAs fall into these two categories and these metrics align with previously developed fit-for-purpose heuristics for ctRSD circuits (14). Additionally, for primers with ≈100 % amplification efficiency, a fraction uncleaved of 0.5 corresponds to a ΔΔCt of 1 (Eq. 1, Methods), which is well within the sensitivity of RT-qPCR (57). In contrast, a fraction uncleaved of >0.75 corresponds to a ΔΔCt values of <0.5, which is generally considered within the acceptable variability for technical replicates (57). So, we expect measurements of sequences for which most RNA is uncleaved to have large uncertainties, justifying the metric of >0.5 fraction uncleaved as the criteria for a poor cleaving sequence. On the other hand, for sequences for which most of the RNA cleaves, the ΔΔCt values should be >2 and less sensitive to experimental noise, thus justifying the metric of <0.1 fraction uncleaved as the criteria for a sequence that cleaves well.

### Ribozyme cleavage during sample preparation confounds RT-qPCR measurements

To validate the RT-qPCR method for measuring ribozyme activity, we first produced the six validation RNAs by IVT and purified these RNAs for RT-qPCR. The six gate sequences in our validation RNAs contain the same ribozyme sequence but different upstream and downstream flanking sequences, which influence the ribozyme’s activity when produced by IVT (Figure 1D). Presumably the G2, G3, and G4 sequences have adopted folds during IVT that disrupt the ribozyme structure and interfere with cleavage activity. These misfolded structures may be metastable, so we were concerned that RNA purification or subsequent reverse transcription could unintentionally induce ribozyme cleavage and result in incorrect measurements. To prevent ribozyme cleavage during RNA purification, we supplemented the RNA purification buffers with ethylenediaminetetraacetic acid (EDTA) to chelate magnesium (Figure 2A), as magnesium is required for ribozyme activity (64).

Many RT-qPCR protocols recommend a denaturing step at ≥65 °C for RNAs with high GC content or strong secondary structure, and reverse transcription is often conducted at (42 to 60) °C to promote destabilization of RNA secondary structure. Interestingly, we found that RNAs that did not cleave when produced by IVT at 37 °C, could be induced to cleave completely when heated to 60 °C for 10 min (Figure 2B), and substantial cleavage was also observed at 45 °C (Supplementary Figure S13). The HDV ribozyme retains activity at high temperatures and under denaturing conditions (65, 66), so increasing temperature may destabilize the interactions in G2, G3, and G4 that interfere with ribozyme folding, allowing the ribozyme to adopt a catalytically active conformation. Surprisingly, we also found that ribozyme cleavage could be induced for G2, G3, and G4 after purification when incubated for 10 min at 37 °C in the RT-qPCR reaction mixture (Figure 2C). This incubation mimics the reverse transcription step of RT-qPCR, and RT-qPCR measurements of G2 cleavage under these conditions resulted in <0.5 fraction uncleaved RNA, contrasting with our gel electrophoresis measurements (Figure 2D).

To explore which components of the RT-qPCR reaction mixture could be inducing unintended ribozyme cleavage after purification, we incubated G2 with different combinations of RT-qPCR components. We found the RT-qPCR reaction buffer without reverse transcriptase or DNA polymerase was sufficient to induce cleavage of G2 at 37 °C after RNA purification. Further, addition of magnesium (to the concentration present in the RT-qPCR reaction mixture) to purified G2 in RNA storage solution also induced substantial cleavage (Supplementary Figure S14). Addition of free magnesium exceeding the concentration in the RT-qPCR reaction buffer to G2 without purification directly after transcription did not change the cleavage observed (Supplementary Figure S15). Together, these results suggest that the RNA changes conformation during the purification process and becomes capable of undergoing cleavage upon reintroduction of magnesium.

Based on the above results, we sought to prevent RNA from cleaving during the reverse transcription step of RT-qPCR. Adding EDTA to the RT-qPCR reaction mixture was not feasible, because free magnesium is required for reverse transcription and PCR (57). We opted to design a short DNA strand, termed a blocking oligonucleotide, that could hybridize to the ribozyme sequence and prevent the ribozyme from folding into a catalytically active conformation (44) (Figure 2A, inset). The blocking oligonucleotide was designed with 22 bases of sequence complementarity to the P2 helix of the ribozyme, a region necessary for cleavage (66) (Supplementary Section 1). Further, a 3’ amino modification was introduced to prevent the blocking oligonucleotide from being extended during RT-qPCR (67). After RNA purification, the blocking oligonucleotide was added to samples and a thermal annealing step was conducted to promote hybridization (Figure 2A and Methods). We found a >150-fold molar excess of blocking oligonucleotide was able to prevent cleavage in the RT-qPCR reaction mixture (Figure 2C and Supplementary Figure S16). The high concentration of blocking oligonucleotide required could be due to the EDTA added to sequester magnesium, or the strong secondary structure of the HDV ribozyme (68). For G2 hybridized with blocking oligonucleotide, the RT-qPCR measurement of ribozyme cleavage yielded >0.75 fraction uncleaved RNA, which aligned with the expected results from denaturing gel electrophoresis measurements (Figure 2D). With the blocking oligonucleotide included, RT-qPCR measurements of ribozyme cleavage yielded similar results for reverse transcription conducted at 37 °C or 50 °C (Figure 2D), and the blocking oligonucleotide prevented G2 cleavage at 65 °C (Supplementary Figure S17). These results indicate the blocking oligonucleotide prevents ribozyme cleavage during reverse transcription to enable accurate RT-qPCR measurements, and this strategy is compatible with many RT-qPCR workflows.

### Addition of a blocking oligonucleotide enables accurate RT-qPCR measurements of ribozyme cleavage in different genetic contexts

We next evaluated RT-qPCR measurements of ribozyme cleavage for the remaining five gate sequences from our validation RNAs (Figure 1D) using the RNA purification method we developed with the blocking oligonucleotide (Figure 2A). We conducted these experiments with a 37 °C reverse transcription step and ran each sample in technical triplicate, with each experiment repeated twice using RNA produced by, and purified from, IVT reactions on separate days. Based on our benchmarking metrics (Supplementary Section 3), the fractions of uncleaved RNA for these gates from RT-qPCR aligned with our denaturing gel electrophoresis results, with G1, G5, and G6 cleaving well (<0.10 fraction uncleaved) and G2, G3, and G4 cleaving poorly (>0.50 fraction uncleaved) for both independent replicates of each gate (Figure 3A).

**Figure 3:**
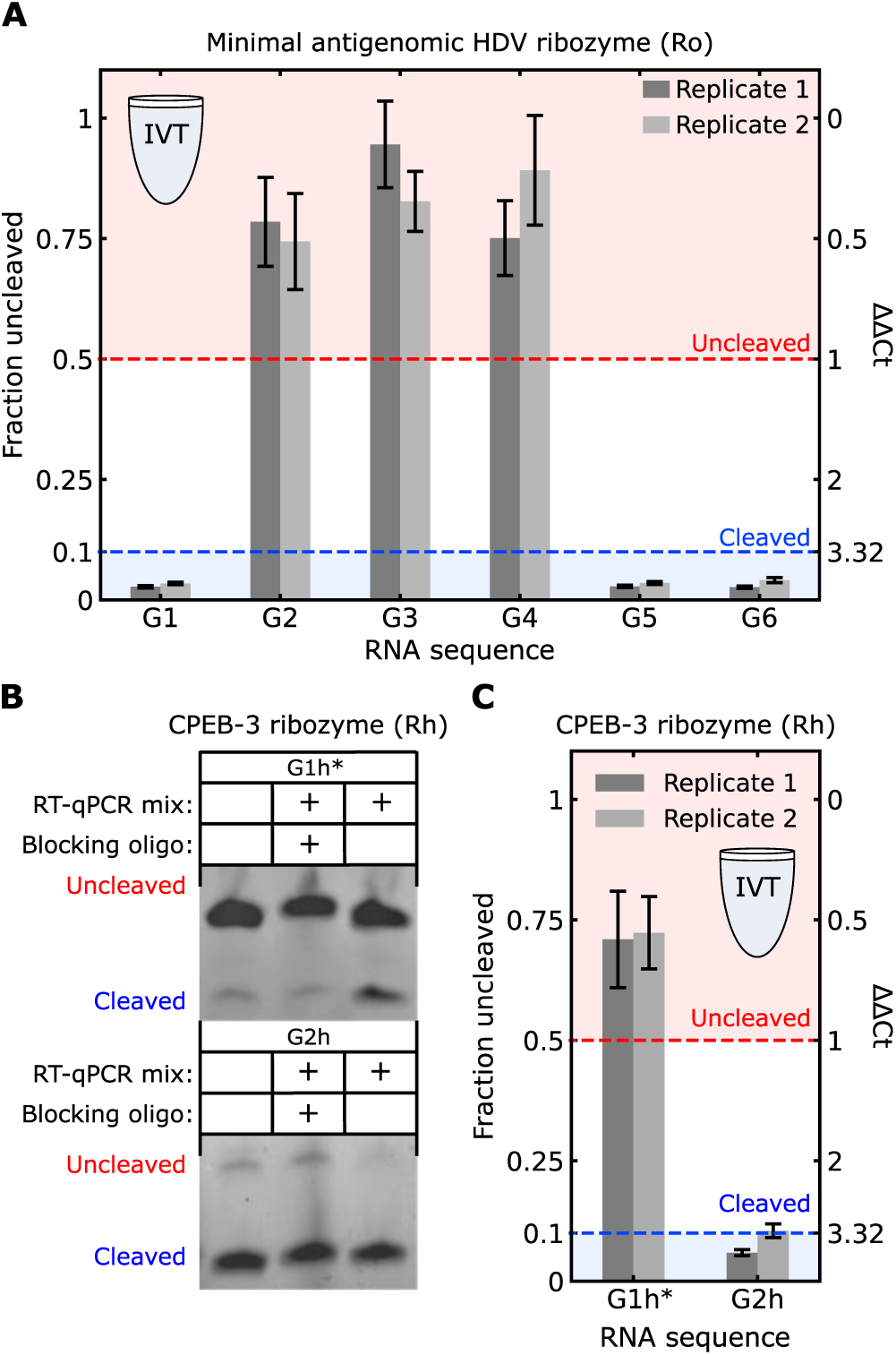
(**A**,**C**) RT-qPCR measurements of ribozyme cleavage for gates produced in IVT. In (A), reverse transcription was conducted at 37 °C. In (C), reverse transcription was conducted at 50 °C. ΔΔCt (Eq. 2, Methods) and corresponding fraction uncleaved RNA are labeled on the right and left axes, respectively. Error bars indicate standard deviation from three technical replicates. Replicate 1 and replicate 2 indicate replicates from two IVT reactions and RNA purifications conducted on different days. (**B**) Denaturing gel electrophoresis results for Rh gates after a 10 min incubation at 37 °C in RT-qPCR reaction mixture with or without BO. Prior to incubation, the RNAs were purified from IVT reactions and annealed with or without blocking oligonucleotide (Methods). G1h* RNA incubated at 50 °C was similarly protected by the blocking oligonucleotide (Supplementary Figure S18). Amplification curves and primer efficiency plots for (A) and (C) are presented in Supplementary Sections 2.4 and 2.5.

We next explored how our workflow would transfer to a different ribozyme sequence. We selected the CPEB-3 ribozyme, which is found in the human genome (69), as it has a similar double pseudoknotted fold to HDV ribozymes (66). Further, this ribozyme has previously been shown to recover the cleavage activity of both the G2 and G3 sequences (14). To adopt our method to the CPEB-3 ribozyme, which we term Rh, new primers and a new blocking oligonucleotide were designed to target similar regions of Rh as the regions targeted in Ro (Supplementary Section 1). We designed variants of the G1 and G2 gates with the CPEB-3 ribozyme (G1h* and G2h), which had different context-dependent cleavage (Figure 3B). G2h has previously been shown to cleave well in IVT by denaturing gel electrophoresis (14). We intentionally designed the G1h* sequence to have poor cleavage by changing the sequence directly downstream of the ribozyme to have complementarity with the P2 helix of Rh (Supplementary Section 1). With G1h*, we confirmed the blocking oligonucleotide sequence for Rh effectively prevented cleavage both in the RT-qPCR reaction mixture at 37 °C (Figure 3B, top) and at 50 °C (Supplementary Figure S18). We also tested RT-qPCR with reverse transcription at either 37 °C and 50 °C. RT-qPCR results were qualitatively similar at both temperatures (Supplementary Figure S10), but 37 °C reverse transcription resulted in a higher fraction uncleaved RNA for G2h than expected, possibly due to differences in reverse transcription efficiency between U and G at the lower temperature (Supplementary Figure S9, S10). With 50 °C reverse transcription, RT-qPCR measurements of Rh cleavage matched the expected cleavage based on denaturing gel electrophoresis measurements for G1h* and G2h (Figure 3C). These results support the generality of adding a blocking oligonucleotide to enable correct RT-qPCR measurements of different ribozyme sequences, and suggest that higher reverse transcription temperatures may be necessary for certain sequences.

### RT-qPCR measurements of ribozyme cleavage for RNAs produced in cells

Having applied our RT-qPCR method with the blocking oligonucleotide to eight different genetic contexts for RNAs produced by IVT (Figure 3), we next explored using the method to measure ribozyme cleavage of gates produced in *E. coli*. For these experiments, we selected two gates that cleaved well when produced by IVT, G1 and G5, and two gates that cleaved poorly when produced by IVT, G2 and G3 (Figure 3A). Each gate sequence was inserted into a plasmid downstream of a constitutively active T7 RNA polymerase (RNAP) promoter, and plasmids were individually transformed into *E. coli* BL21* (DE3). BL21* (DE3) can be induced to express T7 RNAP with isopropyl ß-D-1-thiogalactopyranoside (IPTG) to drive transcription of the gates (Figure 4A). Further, this strain has reduced RNase E activity, so it is often used for RNA expression studies (61). For each gate sequence, we also created plasmids containing the uncleaved (U) control sequence and transformed those into *E. coli* BL21* (DE3) for relative quantification.

**Figure 4:**
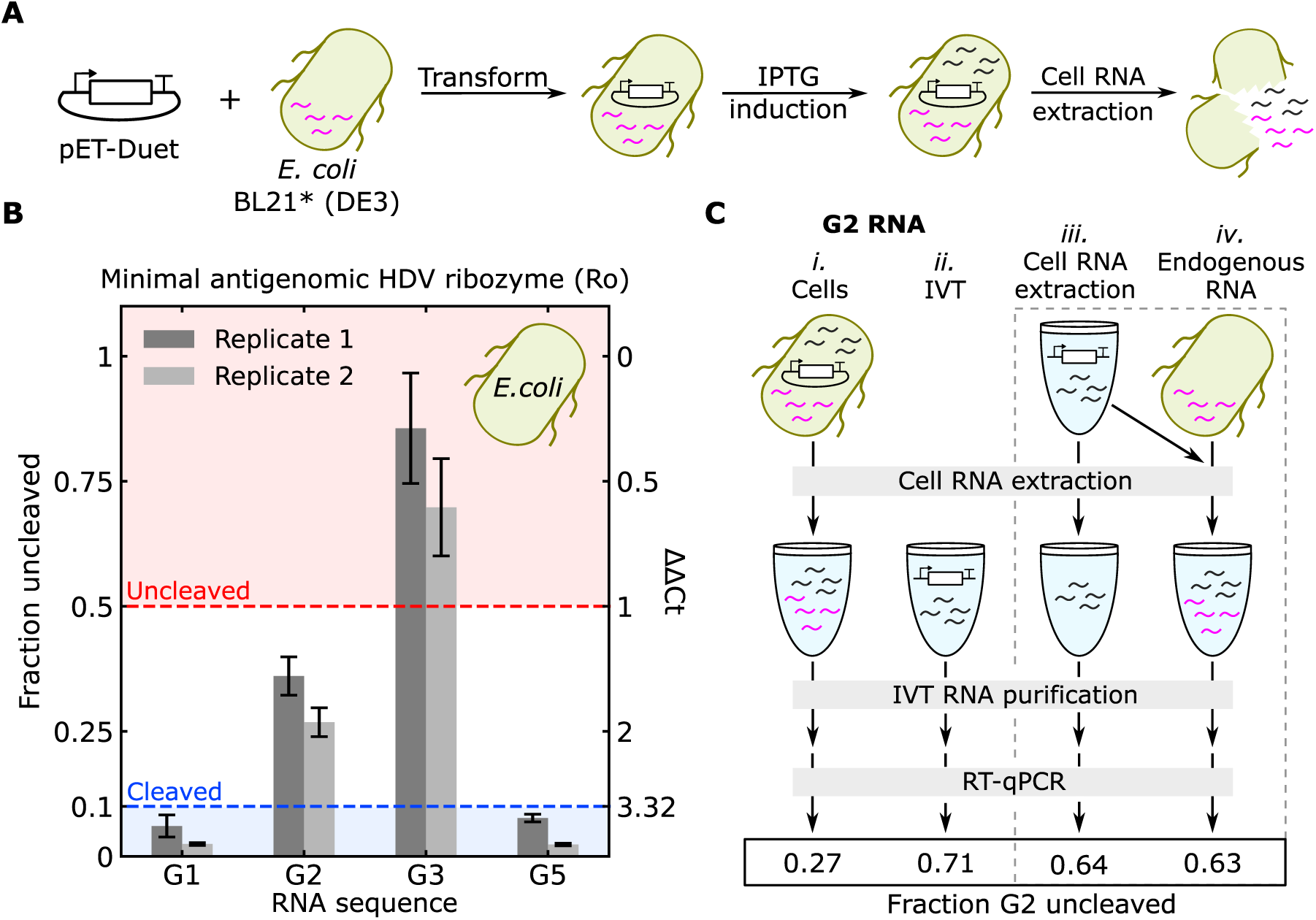
RT-qPCR measurements of ribozyme cleavage of gates produced in *E. coli*. (**A**) The workflow for expressing RNA in *E. coli*. Pink lines and gray lines inside cells indicate endogenous RNA and gate RNA, respectively. Gate sequences were cloned into a pET plasmid backbone and then transformed into *E. coli* BL21* (DE3). Cells cultured with IPTG express T7 RNAP, which transcribes the pET plasmid to produce gate RNA. Total cellular RNA was then extracted for analysis. (**B**) Fraction uncleaved RNA for gates with a minimal antigenomic HDV ribozyme produced in cells. Reverse transcription was conducted at 37 °C. Error bars indicate standard deviation from three technical replicates. Replicate 1 and replicate 2 indicate replicates from two IVT reactions and RNA purifications conducted on different days. Amplification curves and primer efficiency plots are presented in Supplementary Sections 2.4 and 2.5. (**C**) Schematic of controls to evaluate whether the cell RNA extraction protocol influenced the ribozyme cleavage measurement. In sample *iii*, IVT RNA (gray) was taken through the entire cell RNA extraction protocol. In sample *iv*, cells without the plasmid were grown and lysed to yield endogenous RNA (pink), IVT RNA (gray) was added to the lysed cells, and the cell RNA extraction protocol was followed. The numbers in the box below the RT-qPCR arrow indicate the measured fraction uncleaved. Sample *ii* through sample *iv* were all >0.5 fraction uncleaved as expected, indicating cell RNA extraction or the presence of cellular RNA cannot account for the lower fraction uncleaved measured in sample *i*.

Cells were grown to log phase in media supplemented with IPTG, then pelleted and lysed to extract RNA. After extraction, the blocking oligonucleotide protocol was performed on total RNA prior to RT-qPCR. In *E. coli*, the G1, G3, and G5 sequences had similar cleavage as they did when produced by IVT reactions. G2, however, showed a lower fraction uncleaved when produced in *E. coli* (Figure 3B and Figure 4B). It is possible that the additional steps required to extract RNA from cells could confound the RT-qPCR measurement of G2. For example, cell lysis could induce cleavage, or the additional endogenous RNA present in the reaction (we used 50-fold more total RNA by mass from cells than from IVT) could interfere with RT-qPCR, or cause G2 to cleave. To assess these possibilities, we conducted control experiments in which G2 RNA produced by IVT was either taken through the full cell lysis and extraction protocol (Figure 4C, sample *iii*) or added to total RNA extracted from *E. coli* lacking a gate plasmid (Figure 4C, sample *iv*) prior to RT-qPCR. Both controls yielded fractions uncleaved of G2 >0.5 compared to a fraction uncleaved <0.3 for G2 produced in *E. coli*, suggesting G2 does indeed cleave to a greater extent in the cellular environment.

Enhanced ribozyme activity in cells, or environments mimicking the cellular cytoplasm, is often reported compared to activity in IVT reactions, possibly due to differences in ionic conditions or the presence of proteins (16, 17, 70–72). Of our validation RNAs, G2 was the most sensitive to changing conditions, showing substantial cleavage at lower temperatures (Supplementary Figure S13) and cleaving to a greater extent in RT-qPCR reaction mixture without blocking oligonucleotide than G3 and G4 (Figure 2C). It is also possible that differences in degradation rates between cleaved and uncleaved gates within cells influences the ratio of these products. These results show ribozyme cleavage can differ across both genetic and environmental contexts, highlighting the importance of accurate measurement techniques that can be applied across many applications.

## DISCUSSION

Here we developed an RT-qPCR-based method for measuring ribozyme cleavage in different genetic contexts and transcription environments and validated the method using orthogonal measurements of cleavage for RNA produced *in vitro*. The method we developed to prevent ribozyme cleavage with a blocking oligonucleotide during sample preparation and reverse transcription should be compatible with most RT-qPCR kits and protocols. We focused on a general one-step method with SYBR green reporting, but the blocking oligonucleotide method could be adopted for two-step protocols and other probe chemistries (57), or with digital droplet PCR if more sensitive measurements are required (73). We also found the blocking oligonucleotide stabilized ribozymes at 65 °C (Supplementary Figure S17), indicating denaturation steps or higher reverse transcription temperatures could likely be used. However, we found assay performance could vary with different reverse transcription temperatures, depending on the primer set and ribozyme sequence (Supplementary Figures S9, S10), so careful analysis and optimization is warranted when applying the method to new sequences.

Perhaps the most important finding in this study was that misfolded ribozymes with low catalytic activity could be induced to exhibit activity during RNA purification/extraction and reverse transcription, resulting in RT-qPCR measurements overestimating ribozyme activity in the absence of the blocking oligonucleotide. These results highlight the importance of validating new methods with orthogonal measurements. If we had not validated the RT-qPCR method on RNA produced by IVT and introduced the blocking oligonucleotide before applying the method to RNA produced in cells, we would have incorrectly observed substantial cleavage for G2, G3, and G4 in cells – an artifact of sample preparation. These findings have implications for other ribozyme cleavage measurements that require RNA purification/extraction and reverse transcription, such as RNA-seq (32, 38–43). In particular, reports of context-dependent ribozyme cleavage that used different measurement techniques *in vitro* and in cells and found differences in activity may warrant additional examination (25, 45, 46). It is worth noting that HDV-like ribozymes are very stable and retain catalytic activity at high temperatures, so they may be uniquely susceptible to unwanted activity during sample denaturation and reverse transcription compared to other ribozyme folds (48, 68).

More broadly, our findings suggest careful analysis of experimental design when applying any method for RNA analysis that requires RNA extraction or reverse transcription, particularly for structure and function analysis. For example, chemical probing techniques of RNA structure (74), such as selective 2’-hydroxyl acylation analyzed by primer extension sequencing (SHAPE-seq) or dimethyl sulfate probing followed by RNA sequencing (DMS-seq), require RNA extraction and reverse transcription. SHAPE-seq and DMS-seq methods are often conducive with chemical probing *in situ* (75–77), which should provide an accurate picture of the RNA structure irrespective of any rearrangements in subsequent extraction and reverse transcription. But these sequencing-based methods could also be used to obtain both the structure and catalytic activity of a ribozyme, and this could lead to erroneous results if ribozyme activity is not inhibited through cDNA synthesis. In the case of G2, the structure probing results would likely indicate that the ribozyme has adopted a non-native fold, but the ribozyme cleavage results would indicate a high fraction of cleaved RNA, leading to the incorrect conclusion that the non-native fold is catalytically active.

Based on the above observations, the blocking oligonucleotide method we developed should enable accurate measurements of RNA function for a broad range of methods and applications. We only studied ribozymes with HDV-like folds, but the method should apply to other ribozyme classes (64), assuming a suitable blocking oligonucleotide can be designed. In fact, a similar technique has been demonstrated for the hammerhead ribozyme, although the blocking oligonucleotide was removed prior to reverse transcription (44). Our method could also be applied for measurements of aptaswitch activity (6, 34, 36, 50), another important class of RNAs that often use context-dependent cleavage and could be susceptible to structural rearrangements during RNA extraction and reverse transcription. More generally, the method of blocking catalytic activity during sample preparation could be applied to other catalytic nucleic acids, such as ribozymes that perform ligation (51–55) or DNAzymes (49). Accurate measurements of context-dependent RNA catalysis will ultimately support the discovery of new biology and innovations in RNA synthetic biology, biotechnology, and biomanufacturing.

## DATA AVAILABILITY

All MIQE information is provided in the associated methods, protocols, supplementary files, and associated data files. The primer and transcription template sequences used in this study are in Supplementary File S1. The full plasmid sequences for the experiments in *E. coli* are in Supplementary File S2. Uncropped gel images and RT-qPCR data along with the Python analysis code for producing the figures are in Supplementary File S3.

## SUPPLEMENTARY DATA

[Supplementary Information] pdf file containing additional schematics and experimental results

[Supplementary Methods] pdf file containing detailed, step-by-step protocols

[Supplementary File S1] Excel file containing nucleic acid sequences

[Supplementary File S2] GenBank files containing plasmid sequences

[Supplementary File S3] Uncropped gel images, raw RT-qPCR data, and data analysis code

## Supporting information

DNA oligonucleotide and template sequences

Plasmid sequences

Uncropped gel images, raw data, and analysis code

## ACKNOWLEDGMENTS

*Author contributions:* S.W.S conceived the project, designed experiments, and conducted data analysis. N.Y.A. designed and conducted RT-qPCR experiments. O.B.V. conducted plasmid cloning experiments. S.W.S wrote the manuscript with input from all authors.

The authors would like to thank Jason Kralj, David Ross, Elizabeth Strychalski, Molly Wintenberg, and Christina Bergonzo for insightful discussions throughout the project and manuscript preparation.

*Disclaimer:* Certain commercial entities, equipment, or materials may be identified in this document to describe an experimental procedure or concept adequately. Such identification is not intended to imply recommendation or endorsement by the National Institute of Standards and Technology, nor is it intended to imply that the entities, materials, or equipment are necessarily the best available for the purpose. Official contribution of the National Institute of Standards and Technology; not subject to copyright in the United States.

## FUNDING

This work was supported by a National Research Council Postdoctoral Fellowship to S.W.S.

*Conflict of interest statement:* S.W.S has intellectual property related to cotranscriptionally encoded RNA strand displacement circuits (International application number: PCT/US2022/053229). The authors declare no other competing interests.

## Notes

### Competing Interest Statement

The authors have declared no competing interest.

## REFERENCES

1. Webb, C.-H.T., Riccitelli, N.J., Ruminski, D.J. and Lupták, A. (2009) Widespread occurrence of self-cleaving ribozymes. Science, 326, 953–953.

2. Weinberg, C.E., Weinberg, Z. and Hammann, C. (2019) Novel ribozymes: discovery, catalytic mechanisms, and the quest to understand biological function. Nucleic Acids Research, 47, 9480–9494.

3. Jimenez, R.M., Polanco, J.A. and Lupták, A. (2015) Chemistry and biology of self-cleaving ribozymes. Trends Biochem Sci, 40, 648–661.

4. E. Weinberg, C. (2021) Biological Roles of Self-Cleaving Ribozymes. In Ribozymes.pp. 23–53.

5. Lou, C., Stanton, B., Chen, Y.-J., Munsky, B. and Voigt, C.A. (2012) Ribozyme-based insulator parts buffer synthetic circuits from genetic context. Nature Biotechnology, 30, 1137–1142.

6. Dykstra, P.B., Kaplan, M. and Smolke, C.D. (2022) Engineering synthetic RNA devices for cell control. Nat. Rev. Genet., 23, 215–228.

7. Park, S.V., Yang, J.-S., Jo, H., Kang, B., Oh, S.S. and Jung, G.Y. (2019) Catalytic RNA, ribozyme, and its applications in synthetic biology. Biotechnology Advances, 37, 107452.

8. Schürer, H., Lang, K., Schuster, J. and Mörl, M. (2002) A universal method to produce in vitro transcripts with homogeneous 3’ ends. Nucleic Acids Res., 30, e56.

9. Ryczek, M., Pluta, M., Błaszczyk, L. and Kiliszek, A. (2022) Overview of Methods for Large-Scale RNA Synthesis. Applied Sciences, 12.

10. Chen, Y., Cheng, Y. and Lin, J. (2022) A scalable system for the fast production of RNA with homogeneous terminal ends. RNA Biology, 19, 1077–1084.

11. Feng, R., Patil, S., Zhao, X., Miao, Z. and Qian, A. (2021) RNA Therapeutics - Research and Clinical Advancements. Frontiers in Molecular Biosciences, 8.

12. Zhu, Y., Zhu, L., Wang, X. and Jin, H. (2022) RNA-based therapeutics: an overview and prospectus. Cell Death & Disease, 13, 644.

13. Schaffter, S.W. and Strychalski, E.A. (2022) Cotranscriptionally encoded RNA strand displacement circuits. Sci. Adv., 8, eabl4354.

14. Schaffter, S.W., Wintenberg, M.E., Murphy, T.M. and Strychalski, E.A. (2023) Design Approaches to Expand the Toolkit for Building Cotranscriptionally Encoded RNA Strand Displacement Circuits. ACS Synth. Biol., 12, 1546–1561.

15. Bae, W., Stan, G.-B.V. and Ouldridge, T.E. (2021) In situ generation of RNA complexes for synthetic molecular strand-displacement circuits in autonomous systems. Nano Lett., 21, 265–271.

16. Yamagami, R., Sieg, J.P. and Bevilacqua, P.C. (2021) Functional Roles of Chelated Magnesium Ions in RNA Folding and Function. Biochemistry, 60, 2374–2386.

17. Yamagami, R., Bingaman, J.L., Frankel, E.A. and Bevilacqua, P.C. (2018) Cellular conditions of weakly chelated magnesium ions strongly promote RNA stability and catalysis. Nat. Commun., 9, 2149.

18. Sieg, J.P., McKinley, L.N., Huot, M.J., Yennawar, N.H. and Bevilacqua, P.C. (2022) The Metabolome Weakens RNA Thermodynamic Stability and Strengthens RNA Chemical Stability. Biochemistry, 61, 2579–2591.

19. Kato, Y., Kuwabara, T., Warashina, M., Toda, H. and Taira, K. (2001) Relationships between the Activities in Vitro and in Vivo of Various Kinds of Ribozyme and Their Intracellular Localization in Mammalian Cells*. Journal of Biological Chemistry, 276, 15378–15385.

20. Brown, A.L., Perrotta, A.T., Wadkins, T.S. and Been, M.D. (2008) The poly(A) site sequence in HDV RNA alters both extent and rate of self-cleavage of the antigenomic ribozyme. Nucleic Acids Research, 36, 2990–3000.

21. Wang, Y., Wang, Z., Liu, T., Gong, S. and Zhang, W. (2018) Effects of flanking regions on HDV cotranscriptional folding kinetics. RNA, 24, 1229–1240.

22. Chadalavada, D.M., Knudsen, S.M., Nakano, S. and Bevilacqua, P.C. (2000) A role for upstream RNA structure in facilitating the catalytic fold of the genomic hepatitis delta virus ribozyme11Edited by J. A. Doudna. Journal of Molecular Biology, 301, 349–367.

23. Diegelman-Parente, A. and Bevilacqua, P.C. (2002) A Mechanistic Framework for Co-transcriptional Folding of the HDV Genomic Ribozyme in the Presence of Downstream Sequence. Journal of Molecular Biology, 324, 1–16.

24. Wurmthaler, L.A., Klauser, B. and Hartig, J.S. (2018) Highly motif- and organism-dependent effects of naturally occurring hammerhead ribozyme sequences on gene expression. RNA Biology, 15, 231–241.

25. McKinley, L.N., Kern, R.G., Assmann, S.M. and Bevilacqua, P.C. (2023) Flanking Sequence Cotranscriptionally Regulates Twister Ribozyme Activity. Biochemistry, 10.1021/acs.biochem.3c00506.

26. Zingler, N. (2014) The Kinetics of Ribozyme Cleavage: A Tool to Analyze RNA Folding as a Function of Catalysis. In Waldsich, C. (ed), RNA Folding: Methods and Protocols. Humana Press, Totowa, NJ, pp. 209–224.

27. Koseki Shiori, Tanabe Tsuyoshi, Tani Kenzaburo, Asano Shigetaka, Shioda Tatsuo, Nagai Yoshiyuki, Shimada Takashi, Ohkawa Jun, and Taira Kazunari (1999) Factors Governing the Activity In Vivo of Ribozymes Transcribed by RNA Polymerase III. Journal of Virology, 73, 1868–1877.

28. Filonov, G.S., Kam, C.W., Song, W. and Jaffrey, S.R. (2015) In-Gel Imaging of RNA Processing Using Broccoli Reveals Optimal Aptamer Expression Strategies. Chemistry & Biology, 22, 649–660.

29. Andreasson, J.O.L., Savinov, A., Block, S.M. and Greenleaf, W.J. (2020) Comprehensive sequence-to-function mapping of cofactor-dependent RNA catalysis in the glmS ribozyme. Nature Communications, 11, 1663.

30. Singh, K.K., Parwaresch, R. and Krupp, G. (1999) Rapid kinetic characterization of hammerhead ribozymes by real-time monitoring of fluorescence resonance energy transfer (FRET). RNA, 5, 1348–1356.

31. Zhuang, X., Bartley, L.E., Babcock, H.P., Russell, R., Ha, T., Herschlag, D. and Chu, S. (2000) A Single-Molecule Study of RNA Catalysis and Folding. Science, 288, 2048–2051.

32. Xiang, J.S., Kaplan, M., Dykstra, P., Hinks, M., McKeague, M. and Smolke, C.D. (2019) Massively parallel RNA device engineering in mammalian cells with RNA-Seq. Nature Communications, 10, 4327.

33. Aroonsri, A., Kongsee, J., Gunawan, J.D., Aubry, D.A. and Shaw, P.J. (2021) A cell-based ribozyme reporter system employing a chromosomally-integrated 5′ exonuclease gene. BMC Molecular and Cell Biology, 22, 20.

34. Nomura, Y., Zhou, L., Miu, A. and Yokobayashi, Y. (2013) Controlling Mammalian Gene Expression by Allosteric Hepatitis Delta Virus Ribozymes. ACS Synth. Biol., 2, 684–689.

35. Townshend, B., Kennedy, A.B., Xiang, J.S. and Smolke, C.D. (2015) High-throughput cellular RNA device engineering. Nature Methods, 12, 989–994.

36. Wieland, M. and Hartig, J.S. (2008) Improved Aptazyme Design and In Vivo Screening Enable Riboswitching in Bacteria. Angewandte Chemie International Edition, 47, 2604–2607.

37. Ogawa, A. and Maeda, M. (2007) Development of a New-type Riboswitch Using an Aptazyme and an anti-RBS Sequence. Nucleic Acids Symposium Series, 51, 389–390.

38. Yokobayashi, Y. (2020) High-Throughput Analysis and Engineering of Ribozymes and Deoxyribozymes by Sequencing. Acc. Chem. Res., 53, 2903–2912.

39. Kobori, S., Nomura, Y., Miu, A. and Yokobayashi, Y. (2015) High-throughput assay and engineering of self-cleaving ribozymes by sequencing. Nucleic Acids Research, 43, e85–e85.

40. Roberts, J.M., Beck, J.D., Pollock, T.B., Bendixsen, D.P. and Hayden, E.J. (2023) RNA sequence to structure analysis from comprehensive pairwise mutagenesis of multiple self-cleaving ribozymes. eLife, 12, e80360.

41. Olzog, V.J., Gärtner, C., Stadler, P.F., Fallmann, J. and Weinberg, C.E. (2021) cyPhyRNA-seq: a genome-scale RNA-seq method to detect active self-cleaving ribozymes by capturing RNAs with 2ʹ, 3ʹ cyclic phosphates and 5ʹ hydroxyl ends. RNA Biology, 18, 818–831.

42. Peach, S.E., York, K. and Hesselberth, J.R. (2015) Global analysis of RNA cleavage by 5′-hydroxyl RNA sequencing. Nucleic Acids Research, 43, e108–e108.

43. Espah Borujeni, A., Zhang, J., Doosthosseini, H., Nielsen, A.A.K. and Voigt, C.A. (2020) Genetic circuit characterization by inferring RNA polymerase movement and ribosome usage. Nature Communications, 11, 5001.

44. Kim, M.W., Sun, G., Lee, J.H. and Kim, B. (2018) Development of Quenching-qPCR (Q-Q) assay for measuring absolute intracellular cleavage efficiency of ribozyme. Analytical Biochemistry, 550, 27–33.

45. Vlková, M., Morampalli, B.R. and Silander, O.K. (2021) Efficiency of the synthetic self-splicing RiboJ ribozyme is robust to cis- and trans-changes in genetic background. MicrobiologyOpen, 10, e1232.

46. Roth, A., Weinberg, Z., Chen, A.G.Y., Kim, P.B., Ames, T.D. and Breaker, R.R. (2014) A widespread self-cleaving ribozyme class is revealed by bioinformatics. Nature Chemical Biology, 10, 56–60.

47. He, S.L. and Green, R. (2013) Chapter Three - Northern Blotting. In Lorsch, J. (ed), Methods in Enzymology. Academic Press, Vol. 530, pp. 75–87.

48. Webb, C.-H.T. and Lupták, A. (2011) HDV-like self-cleaving ribozymes. RNA Biol., 8, 719–727.

49. Zimmermann, A.C., White, I.M. and Kahn, J.D. (2020) Nucleic acid-cleaving catalytic DNA for sensing and therapeutics. Talanta, 211, 120709.

50. Felletti, M. and Hartig, J.S. (2017) Ligand-dependent ribozymes. WIREs RNA, 8, e1395.

51. Gambill, L., Staubus, A., Mo, K.W., Ameruoso, A. and Chappell, J. (2023) A split ribozyme that links detection of a native RNA to orthogonal protein outputs. Nature Communications, 14, 543.

52. Hieronymus, R., Zhu, J. and Müller, S. (2022) RNA self-splicing by engineered hairpin ribozyme variants. Nucleic Acids Research, 50, 368–377.

53. Hausner, G., Hafez, M. and Edgell, D.R. (2014) Bacterial group I introns: mobile RNA catalysts. Mobile DNA, 5, 8.

54. Hedberg, A. and Johansen, S.D. (2013) Nuclear group I introns in self-splicing and beyond. Mobile DNA, 4, 17.

55. Kalvapalle, P.B., Staubus, A., Dysart, M.J., Gambill, L., Reyes Gamas, K., Chieh Lu, L., Silberg, J.J., Stadler, L.B. and Chappell, J. (2023) Information storage across a microbial community using universal RNA memory. bioRxiv, 10.1101/2023.04.16.536800.

56. Gibson, D.G., Young, L., Chuang, R.-Y., Venter, J.C., Hutchison, C.A. and Smith, H.O. (2009) Enzymatic assembly of DNA molecules up to several hundred kilobases. Nat. Methods, 6, 343–345.

57. Bustin, S.A. (2004) A-Z of Quantitative PCR International University Line.

58. Pfaffl, M.W. (2001) A new mathematical model for relative quantification in real-time RT–PCR. Nucleic Acids Research, 29, e45–e45.

59. Taylor, J.R. (2022) An Introduction to Error Analysis: The Study of Uncertainties in Physical Measurements University Science Books.

60. Jung, J.K., Archuleta, C.M., Alam, K.K. and Lucks, J.B. (2022) Programming cell-free biosensors with DNA strand displacement circuits. Nat. Chem. Biol., 18, 385–393.

61. Green, A.A., Kim, J., Ma, D., Silver, P.A., Collins, J.J. and Yin, P. (2017) Complex cellular logic computation using ribocomputing devices. Nature, 548, 117–121.

62. Chappell, J., Westbrook, A., Verosloff, M. and Lucks, J.B. (2017) Computational design of small transcription activating RNAs for versatile and dynamic gene regulation. Nat. Commun., 8, 1051.

63. Udvardi, M.K., Czechowski, T. and Scheible, W.-R. (2008) Eleven Golden Rules of Quantitative RT-PCR. The Plant Cell, 20, 1736–1737.

64. Ferré-D’Amaré, A.R. and Scott, W.G. (2010) Small Self-cleaving Ribozymes. Cold Spring Harbor Perspectives in Biology, 2.

65. Smith, J.B. and Dinter-Gottlieb, G. (1991) Antigenomic Hepatitis delta virus ribozymes self-cleave in 18 M formamide. Nucleic Acids Research, 19, 1285–1289.

66. Riccitelli, N. and Lupták, A. (2013) HDV family of self-cleaving ribozymes. In Soukup, G.A. (ed), Progress in Molecular Biology and Translational Science. Elsevier, Amsterdam, The Netherlands, Vol. 120, pp. 123–171.

67. Mizuno, Y., Carninci, P., Okazaki, Y., Tateno, M., Kawai, J., Amanuma, H., Muramatsu, M. and Hayashizaki, Y. (1999) Increased specificity of reverse transcription priming by trehalose and oligo-blockers allows high-efficiency window separation of mRNA display. Nucleic Acids Research, 27, 1345–1349.

68. Duhamel, J., Liu, D.M., Evilia, C., Fleysh, N., Dinter-Gottlieb, G. and Lu, P. (1996) Secondary structure content of the HDV ribozyme in 95% formamide. Nucleic Acids Res., 24, 3911–3917.

69. Salehi-Ashtiani, K., Lupták, A., Litovchick, A. and Szostak, J.W. (2006) A Genomewide Search for Ribozymes Reveals an HDV-Like Sequence in the Human CPEB3 Gene. Science, 313, 1788–1792.

70. Brown, T.S., Chadalavada, D.M. and Bevilacqua, P.C. (2004) Design of a Highly Reactive HDV Ribozyme Sequence Uncovers Facilitation of RNA Folding by Alternative Pairings and Physiological Ionic Strength. Journal of Molecular Biology, 341, 695–712.

71. Herschlag, D., Khosla, M., Tsuchihashi, Z. and Karpel, R.L. (1994) An RNA chaperone activity of non-specific RNA binding proteins in hammerhead ribozyme catalysis. The EMBO Journal, 13, 2913–2924.

72. Ruminski, D.J., Watson, P.Y., Mahen, E.M. and Fedor, M.J. (2016) A DEAD-box RNA helicase promotes thermodynamic equilibration of kinetically trapped RNA structures in vivo. RNA, 22, 416–427.

73. Taylor, S.C., Laperriere, G. and Germain, H. (2017) Droplet Digital PCR versus qPCR for gene expression analysis with low abundant targets: from variable nonsense to publication quality data. Scientific Reports, 7, 2409.

74. Spitale, R.C. and Incarnato, D. (2023) Probing the dynamic RNA structurome and its functions. Nature Reviews Genetics, 24, 178–196.

75. Watters, K.E., Yu, A.M., Strobel, E.J., Settle, A.H. and Lucks, J.B. (2016) Characterizing RNA structures in vitro and in vivo with selective 2′-hydroxyl acylation analyzed by primer extension sequencing (SHAPE-Seq). Methods, 103, 34–48.

76. Zubradt, M., Gupta, P., Persad, S., Lambowitz, A.M., Weissman, J.S. and Rouskin, S. (2017) DMS-MaPseq for genome-wide or targeted RNA structure probing in vivo. Nat. Methods, 14, 75–82.

77. Smola, M.J. and Weeks, K.M. (2018) In-cell RNA structure probing with SHAPE-MaP. Nature Protocols, 13, 1181–1195.

