## Supplementary Information for "Prevention of ribozyme catalysis through cDNA synthesis enables accurate RT-qPCR measurements of context-dependent ribozyme activity"

### 1 Sequences and schematics

#### 1.1 RNA sequences and schematics

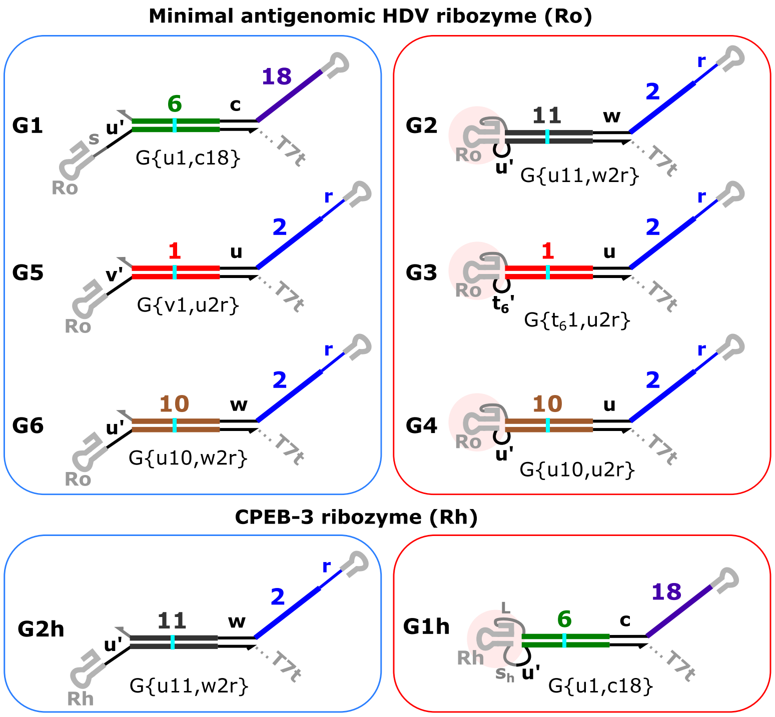

**Supplementary Figure S1:** Schematics of the ctRSD gates used for RT-qPCR method validation. The schematics follow ctRSD toolkit domain naming conventions (1). The full domain specific names of the gates from the ctRSD toolkit are shown below the gates. Sequence schematics shown below in Supplementary Figure S2.

**Supplementary Table S1:** Annotated ctRSD gate RNA sequences. Red text corresponds to the portion of the ribozyme sequence that the blocking oligo binds. Underlined text represents the binding sites for PCRu (pink underlines) and PCRo (black underlines) primers. Primer sequences are in Supplementary Table S2. *Italic text* represents the portions of upstream (US) and downstream (DS) sequences that vary from gate to gate. For the uncleaved control transcripts (U), the **bold red C** was mutated to a U base. For the cleaved control transcripts (C), the upstream (US) sequence was removed. Supplementary Figure S2 shows the annotated gate secondary structures.

| Gate | RNA sequence |
|---|---|
| | Minimal antigenomic HDV ribozyme (Ro) |
| **G1**<br>G{u6, c18} | US:GGGCUCGAUCACUAAUCUGAUCGAGACA*AACAUACCUACCUUCAUUAUCUUACUUAC*<br>Ro:GGGUCGGCAUGGCAUCUCCACCUCCUCGCGGUCCGACCUGGGCUACUUCGGUAGG**C**UAAGGGAGAG<br>DS:*AGAGUAUGAUGUAGUAAGGUAAUGAAGG*AGCAUAACCCCUUGGGGCCUCUAAACGGGUCUUGAGGGGUUUUUUG<br>PCRu: 66 bases, PCRo: 112 bases |
| **G2**<br>G{u11, w2r} | US:GGGCUCGAUCACUAAUCUGAUCGAGACA*CUACAUCCACAUACUAAUUAACAUACCUCUUAUUC*<br>Ro:GGGUCGGCAUGGCAUCUCCACCUCCUCGCGGUCCGACCUGGGCUACUUCGGUAGG**C**UAAGGGAGAG<br>DS:*UGAUGUUAAGAGGUGUGUUAAU*AGCAUAACCCCUUGGGGCCUCUAAACGGGUCUUGAGGGGUUUUUUG<br>PCRu: 72 bases, PCRo: 106 bases |
| **G3**<br>G{$t_{6}$1, u2r} | US:GGGCUCGAUCACUAAUCUGAUCGAGACA*CUACAUCCACAUACUAACAUCAUUACUUCACAUUC*<br>Ro:GGGUCGGCAUGGCAUCUCCACCUCCUCGCGGUCCGACCUGGGCUACUUCGGUAGG**C**UAAGGGAGAG<br>DS:*GAGAGUUGUGAAGUGAUGAUGU*AGCAUAACCCCUUGGGGCCUCUAAACGGGUCUUGAGGGGUUUUUUG<br>PCRu: 72 bases, PCRo: 106 bases |
| **G4**<br>G{u10, u2r} | US:GGGCUCGAUCACUAAUCUGAUCGAGACA*CUACAUCCACAUACUAACAUCACUUACUCCUAUUC*<br>Ro:GGGUCGGCAUGGCAUCUCCACCUCCUCGCGGUCCGACCUGGGCUACUUCGGUAGG**C**UAAGGGAGAG<br>DS:*UGAUGUUAGGAGUAGGUGAUGU*AGCAUAACCCCUUGGGGCCUCUAAACGGGUCUUGAGGGGUUUUUUG<br>PCRu: 72 bases, PCRo: 106 bases |
| **G5**<br>G{v1, u2r} | US:GGGCUCGAUCACUAAUCUGAUCGAGACA*CUACAUCCACAUACUAACAUCAUUACUUCACAUUC*<br>Ro:GGGUCGGCAUGGCAUCUCCACCUCCUCGCGGUCCGACCUGGGCUACUUCGGUAGG**C**UAAGGGAGAG<br>DS:*AGGAUUUGUGAAGUGAUGAUGU*AGCAUAACCCCUUGGGGCCUCUAAACGGGUCUUGAGGGGUUUUUUG<br>PCRu: 72 bases, PCRo: 106 bases |
| **G6**<br>G{u10, w2r} | US:GGGCUCGAUCACUAAUCUGAUCGAGACA*CUACAUCCACAUACUAAUUAACCUUACUCCUAUUC*<br>Ro:GGGUCGGCAUGGCAUCUCCACCUCCUCGCGGUCCGACCUGGGCUACUUCGGUAGG**C**UAAGGGAGAG<br>DS:*UGAUGUUAGGAGUAGGGUUAAU*AGCAUAACCCCUUGGGGCCUCUAAACGGGUCUUGAGGGGUUUUUUG<br>PCRu: 72 bases, PCRo: 106 bases |
| | CPEB-3 ribozyme (Rh) |
| **G1h***<br>G{u6, c18} | US:GGGCUCGAUCACUAAUCUGAUCGAGACA*AACAUACCUACCUUCAUUAUCUUACUUAC*<br>Rh:GGGGGCCACAGCAGAAGCGUUCACGUCGCAGCCCCUGUCAGAUUCUGGUGAAUCUG**C**GAAUUCUGCU<br>DS:*GCUGUUGAUGUAGUAAGGUAAUGAAGG*AGCAUAACCCCUUGGGGCCUCUAAACGGGUCUUGAGGGGUUUUUUG<br>PCRu: 67 bases, PCRo: 112 bases |
| **G2h**<br>G{u11, w2r} | US:GACUCGAUCACUAAUCUGAUCGAGACA*CUACAUCCACAUACUAAUUAACAUACCUCUUAUUC*<br>Rh:GGGGGCCACAGCAGAAGCGUUCACGUCGCAGCCCCUGUCAGAUUCUGGUGAAUCUG**C**GAAUUCUGCU<br>DS:*UGAUGUUAAGAGGUGUGUUAAU*AGCAUAACCCCUUGGGGCCUCUAAACGGGUCUUGAGGGGUUUUUUG<br>PCRu: 66 bases, PCRo: 107 bases |

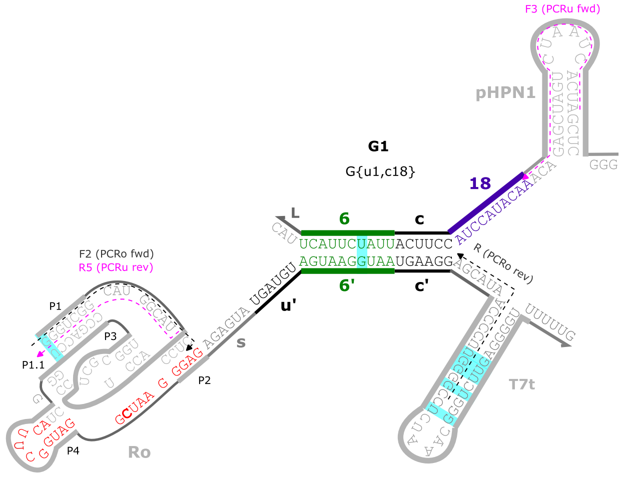
G1
G{u1,c18}
F3 (PCRu fwd)
pHPN1
F2 (PCRo fwd)
R5 (PCRu rev)
R (PCRo rev)
P1
P1.1
P2
P3
P4
Ro
T7t
L
6
c
18
6'
c'
u'
s
UCAUUCUAUUACUUCC
AGUAAGGUAAUGAAGG
AUCCAUACAA
UGAUGU
AGAGUA
GGG
UUUUG

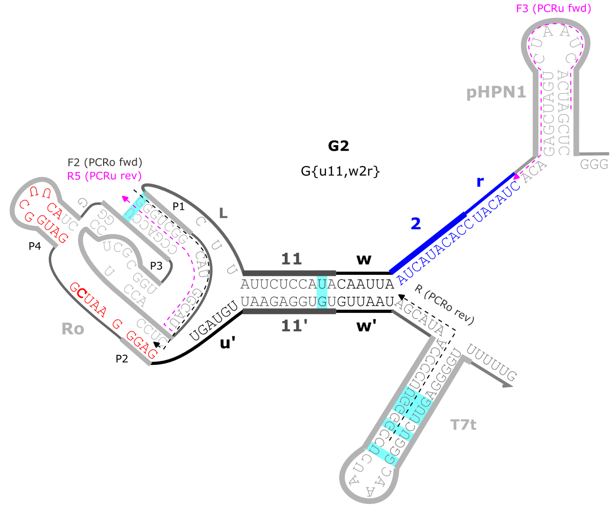
G2
G{u11,w2r}
F3 (PCRu fwd)
pHPN1
F2 (PCRo fwd)
R5 (PCRu rev)
R (PCRo rev)
P1
P2
P3
P4
Ro
T7t
L
11
w
2
r
11'
w'
u'
AUUCUCCAUACAAUUA
UAAGAGGUGUGUUAAU
AUCAUACACCUACAUC
UGAUGU
GGG
UUUUG

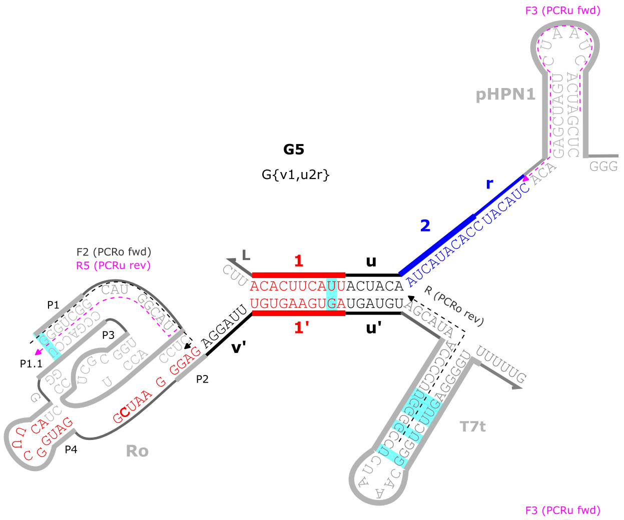
G5
G{v1,u2r}
F3 (PCRu fwd)
pHPN1
F2 (PCRo fwd)
R5 (PCRu rev)
R (PCRo rev)
P1
P1.1
P2
P3
P4
Ro
T7t
L
1
u
2
r
1'
u'
v'
ACACUUCAUUACUACA
UGUGAAGUGAUGAUGU
AUCAUACACCUACAUC
AGGAUU
GGG
UUUUG

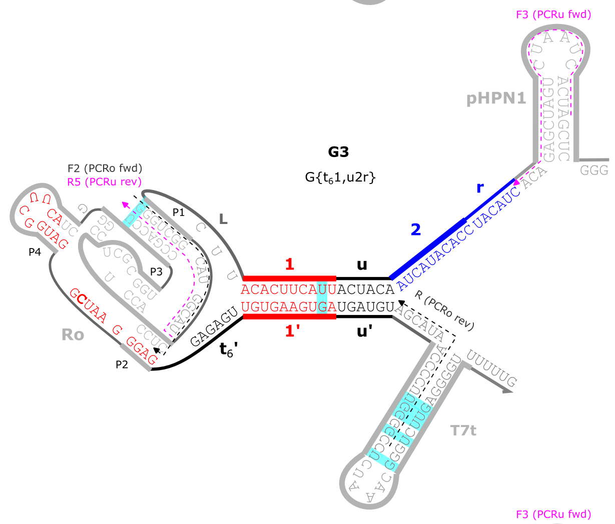
G3
G{t61,u2r}
F3 (PCRu fwd)
pHPN1
F2 (PCRo fwd)
R5 (PCRu rev)
R (PCRo rev)
P1
P2
P3
P4
Ro
T7t
L
1
u
2
r
1'
u'
t6'
ACACUUCAUUACUACA
UGUGAAGUGAUGAUGU
AUCAUACACCUACAUC
GAGAGU
GGG
UUUUG

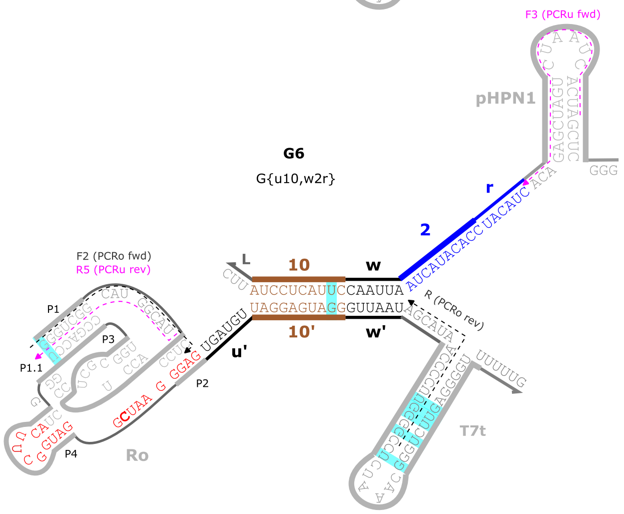
G6
G{u10,w2r}
F3 (PCRu fwd)
pHPN1
F2 (PCRo fwd)
R5 (PCRu rev)
R (PCRo rev)
P1
P1.1
P2
P3
P4
Ro
T7t
L
10
w
2
r
10'
w'
u'
AUCCUCAUUCCAAUUA
UAGGAGUAGGGUUAAU
AUCAUACACCUACAUC
UGAUGU
GGG
UUUUG

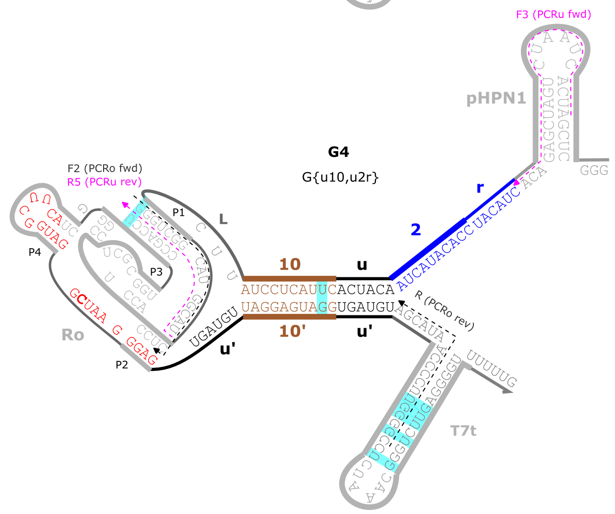
G4
G{u10,u2r}
F3 (PCRu fwd)
pHPN1
F2 (PCRo fwd)
R5 (PCRu rev)
R (PCRo rev)
P1
P2
P3
P4
Ro
T7t
L
10
u
2
r
10'
u'
AUCCUCAUUCACUACA
UAGGAGUAGGUGAUGU
AUCAUACACCUACAUC
UGAUGU
GGG
UUUUG

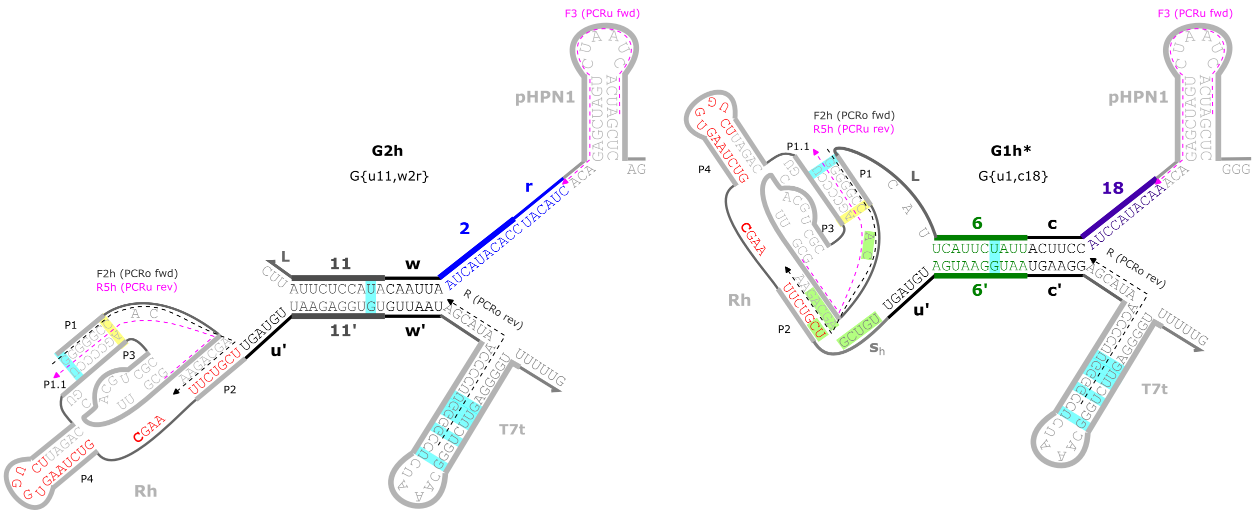

**Supplementary Figure S2:** Gate secondary structure schematics. As the exact structures are not known, the intended secondary structure is shown for the gates that cleave well and an uncleaved ribozyme is shown for the gates that do not cleave well. Presumably, the gates that do not cleave well have folds that disrupt the ribozyme structure. Red bases in the ribozyme sequence indicate the sequence the blocking oligo binds to within the ribozyme. The bold C base in the ribozyme sequence indicates the base that was changed to a U base to abolish cleavage in the uncleaved control RNAs. Pink and gray dashed lines indicate the sequences the PCRu and PCRo bind to, respectively. Cyan highlighted bases indicate GU wobble base pairs. The green highlighted bases in Rh of G1h* represent alternative base pairings that were designed to cause the ribozyme to misfold.

#### 1.2 DNA templates

DNA templates encoding for G, U, and C RNAs were ordered as eBlocks from IDT. Example annotated templates are shown below. To meet the length requirements for ordering (300 bases), the lowercase gray flanking sequence was appended to the 3' end of the sequences. This flanking sequence was not amplified in subsequent PCRs. All the DNA sequences from this study are in Supporting File S3.

Underlined regions indicate primer binding sequences for PCR amplification of templates. The primers specific to the T7 promoter and T7 terminator were used to prepare DNA templates for *in vitro* transcription. The primers specific to the ga4 and pET BB homology domains were used to prepare DNA insert for cloning into plasmids.

Legend:
ga4 homology domain, T7 promoter, Ribozyme, pET BB homology domain, *T7 terminator*,
5' HAIRPIN, ctRSD output domain, ctRSD input domains, **ctRSD toehold domains**, excess

**G1:** G{u6,c18} eBlock DNA

5'gaagtctaacgctgctctgggctaactgtcGCGCTAATACGACTCACTATAGGGCTCGATCACTAATCTGATCGA
GACAAACATACCTA**CCTTCA**TTATCTTACTTACGGGTCGGCATGGCATCTCCACCTCCTCGCGGTCCGACCTGGGCT
ACTTCGGTAGGCTAAGGGAGAGAGTA**TGATGT**AGTAAGGTAA**TGAAGG***AGCATAACCCCTTGGGGCCTCTAAACGGG*
*TCTTGAGGGGTTTTTTG*ctgaaaggaggaactatatccggattggcgctgaaacctcaggcatttgagaag

**U1:** U{u6,c18} eBlock DNA

5'gaagtctaacgctgctctgggctaactgtcGCGCTAATACGACTCACTATAGGGCTCGATCACTAATCTGATCGA
GACAAACATACCTACCTTCATTATCTTACTTACGGGTCGGCATGGCATCTCCACCTCCTCGCGGTCCGACCTGGGCT
ACTTCGGTAGG**T**TAAGGGAGAGAGTATGATGTAGTAAGGTAATGAAGG*AGCATAACCCCTTGGGGCCTCTAAACGGG*
*TCTTGAGGGGTTTTTTG*ctgaaaggaggaactatatccggattggcgctgaaacctcaggcatttgagaag

**C1:** C{u6,c18} eBlock DNA

5'gaagtctaacgctgctctgggctaactgtcGCGCTAATACGACTCACTATAGGGTCGGCATGGCATCTCCACCTC
CTCGCGGTCCGACCTGGGCTACTTCGGTAGGCTAAGGGAGAGAGTATGATGTAGTAAGGTAATGAAGG*AGCATAACC*
*CCTTGGGGCCTCTAAACGGGTCTTGAGGGGTTTTTTG*ctgaaaggaggaactatatccggattggcgctgaaacctc
aggcatttgagaagcacacgactgtgtacttgttataacatctgacagttaaagtcgggagaataggagcc

#### 1.3 RT-qPCR primers and blocking oligos

**Supplementary Table S2:** RT-qPCR primers and blocking oligos used in this study. The blocking oligos have a 3' amino modification (/3AmMO/) to prevent extension during RT-qPCR. The blocking oligos were used for the gate RNA and the uncleaved and cleaved control RNAs. Because the uncleaved control RNA has a cytosine to uracil mutation to abolish activity, the blocking oligos will have a GU wobble base pair at this mutation site (highlighted) when bound to the uncleaved control RNAs rather than the GC base pair present when bound to the gate RNA.

| **RT-qPCR primers** | | |
|---|---|---|
| F3 | 5'ATCACTAATCTGATCGAGACA | PCRu_Ro_fwd, PCRu_Rh_fwd |
| R5 | 5'GATGCCATGCCGACCC | PCRu_Ro_rev |
| F2 | 5'GGGTCGGCATGGCATC | PCRo_Ro_fwd |
| R | 5'GCCCCAAGGGGTTATGCT | PCRo_Ro_rev, PCRo_Rh_rev |
| F2h | 5'GGGGGCCACAGCAGAAG | PCRo_Rh_fwd |
| R5h | 5'CTTCTGCTGTGGCCCCC | PCRu_Rh_rev |
| **Blocking Oligos** | | |
| BO_Ro | 5'CTCCCTTA**G**CCTACCGAAGT/3AmMO/ | Blocking oligo for Ro |
| BO_Rh | 5'AGCAGAATTC**G**CAGATTCACCAGT/3AmMO/ | Blocking oligo for Rh |

### 2 Additional RT-qPCR results and data analysis details

#### 2.1 Plate layout and experimental setup

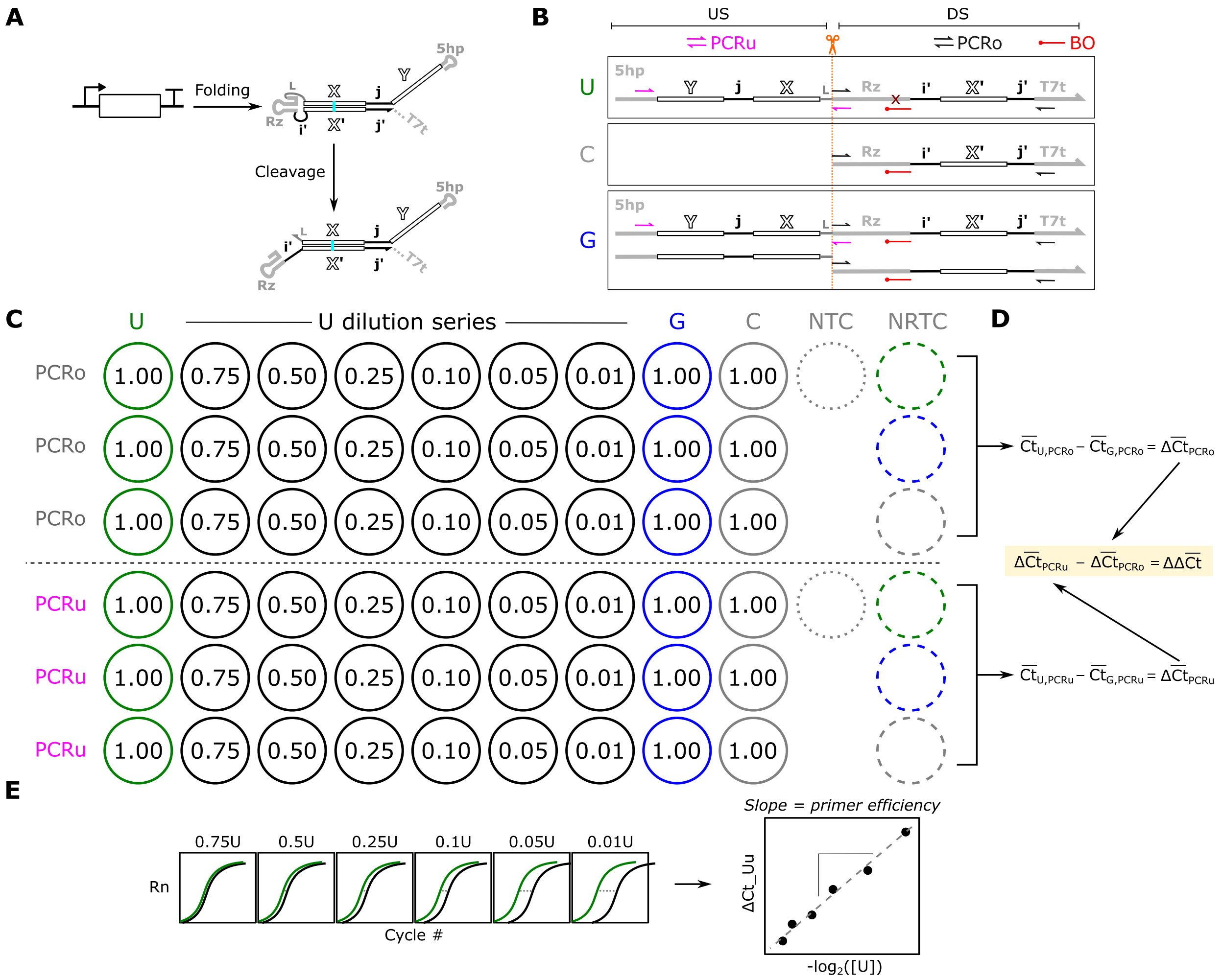

**Supplementary Figure S3:** Experimental setup and analysis overview. (**A**) ctRSD gate schematic. (**B**) PCR primer layout for a ctRSD gate (G) and corresponding uncleaved (U) and cleaved (C) control transcripts. (**C**) A typical plate layout for an RT-qPCR experiment for a single gate sequence. The first three rows are technical replicates for PCRo and the second three rows are technical replicates for PCRu. The 1$^{st}$, 8$^{th}$, and 9$^{th}$ columns are U, G, and C RNAs at 1x concentration (0.08 pg/µL). U and G samples are used to determine ΔΔCt (panel D). The 2$^{nd}$ through 7$^{th}$ columns are dilution series of U used to determine primer amplification efficiency (panel E). NTC is a no template (no RNA) control for the two primer sets. NRTC is a no reverse transcriptase control for each of the RNAs to assess potential DNA contamination. (**D**) ΔΔCt analysis. The average Ct of U and G for PCRo were used to obtain an average ΔCt for PCRo. The average Ct of U and G for PCRu were used to obtain an average ΔCt for PCRu. These average ΔCt values were then used to compute the average ΔΔCt used to compute fraction uncleaved gate. (**E**) Illustration of U dilution series used to assess primer amplification efficiency. The ΔCt of dilutions of U compared to 1x U were plotted against the negative log base 2 of the dilution fraction. A line was then fit to this data with a slope of 1 indicating 100% amplification efficiency. Two types of U dilution series were used in this study: dilutions with C RNA solutions or dilutions with RNA storage solution (ThermoFisher, AM7001). For dilutions with C RNA, samples of U RNA and C RNA at 1 pg/µL were mixed at the appropriate volume ratios to reach the desired U dilution. For example, a 0.75 U sample was prepared by mixing 75 µL of 1 pg/µL U RNA was mixed with 25 µL of 1 pg/µL C RNA. Likewise, for the 0.01 U sample, 1 µL of 1 pg/µL U RNA was mixed with 99 µL of 1 pg/µL C RNA. For dilutions without C RNA, the same dilutions of 1 pg/µL U RNA were conducted with RNA storage solution rather than C RNA solutions.

#### 2.2 Detailed data analysis and error propagation

***Comparison to conventional* $\Delta\Delta$*Ct methods*:** Our analysis is based on a relative quantification technique termed the ΔΔCt method (2–4). In conventional ΔΔCt analysis the change in expression of a gene of interest in a treated and untreated sample is compared to the change in expression of a reference gene that should have constant expression across treated and untreated samples. Inclusion of the reference gene controls for differences in the amount of total RNA added between the treated and untreated samples. Our method is analogous (Supplementary Table S3), but instead of a reference gene it uses a reference PCR (PCRo) that amplifies both uncleaved and cleaved RNA. PCRo should have the same Ct values between the G and U samples if the same amount of RNA is added. Similarly, instead of untreated and treated samples, our method has uncleaved (U) and unknown cleavage (G) samples, which are tested using primers that only amplify uncleaved RNA (PCRu).

**Supplementary Table S3:** Comparison of samples in a conventional ΔΔCt to samples in the method presented here.

| Conventional ΔΔCt method | Our method |
|---|---|
| Reference gene, untreated (Ru) | U, PCRo (Uo) |
| Reference gene, treated (Rt) | G, PCRo (Go) |
| Gene of interest, untreated (Iu) | U, PCRu (Uu) |
| Gene of interest, treated (It) | G, PCRu (Gu) |

Convential: $\Delta\Delta\mathrm{Ct} = (\mathrm{Ct}_{\mathrm{Iu}} - \mathrm{Ct}_{\mathrm{It}}) - (\mathrm{Ct}_{\mathrm{Ru}} - \mathrm{Ct}_{\mathrm{Rt}}) = \Delta\mathrm{Ct}_{\mathrm{I}} - \Delta\mathrm{Ct}_{\mathrm{R}}$

Our method: $\Delta\Delta\mathrm{Ct} = (\mathrm{Ct}_{\mathrm{U,PCRu}} - \mathrm{Ct}_{\mathrm{G,PCRu}}) - (\mathrm{Ct}_{\mathrm{U,PCRo}} - \mathrm{Ct}_{\mathrm{G,PCRo}}) = \Delta\mathrm{Ct}_{\mathrm{PCRu}} - \Delta\mathrm{Ct}_{\mathrm{PCRo}}$

As described in the Methods of the main text, Eq. 1 and Eq. 2 below were used to calculate the fraction uncleaved of a given gate sequence.

$$\text{Eq. 1: Fraction uncleaved} = 2^{-\Delta\Delta\overline{\mathrm{Ct}}}$$

$$\text{Eq. 2: } \Delta\Delta\overline{\mathrm{Ct}} = (\overline{\mathrm{Ct}}_{\mathrm{U,PCRu}} - \overline{\mathrm{Ct}}_{\mathrm{G,PCRu}}) - (\overline{\mathrm{Ct}}_{\mathrm{U,PCRo}} - \overline{\mathrm{Ct}}_{\mathrm{G,PCRo}}) = \overline{\Delta\mathrm{Ct}}_{\mathrm{PCRu}} - \overline{\Delta\mathrm{Ct}}_{\mathrm{PCRo}}$$

***Error propagation*:** Error in the final fraction uncleaved measurements were obtained as described below based on Ref (5).

In Eq. 2, both $\overline{\mathrm{Ct}}_{\mathrm{U,PCRu}}$ and $\overline{\mathrm{Ct}}_{\mathrm{G,PCRu}}$ have associated uncertainties, *i.e.*, standard deviation of the mean from the technical replicates. So, the standard deviation for $\overline{\Delta\mathrm{Ct}}_{\mathrm{PCRu}}$ is given by:

$$\sigma_{\overline{\Delta\mathrm{Ct}}_{\mathrm{PCRu}}} = \sqrt{\left(\sigma_{\overline{\mathrm{Ct}}_{\mathrm{U,PCRu}}}\right)^2 + \left(\sigma_{\overline{\mathrm{Ct}}_{\mathrm{G,PCRu}}}\right)^2}$$

Likewise, both $\overline{\mathrm{Ct}}_{\mathrm{U,PCRo}}$ and $\overline{\mathrm{Ct}}_{\mathrm{G,PCRo}}$ have associated uncertainties, *i.e.*, standard deviation of the mean from the technical replicates. So, the standard deviation for $\overline{\Delta\mathrm{Ct}}_{\mathrm{PCRo}}$ is given by:

$$\sigma_{\overline{\Delta\mathrm{Ct}}_{\mathrm{PCRo}}} = \sqrt{\left(\sigma_{\overline{\mathrm{Ct}}_{\mathrm{U,PCRo}}}\right)^2 + \left(\sigma_{\overline{\mathrm{Ct}}_{\mathrm{G,PCRo}}}\right)^2}$$

Then the standard deviation for ΔΔCt is obtained from the standard deviations of $\overline{\Delta Ct}_{PCRu}$ and $\overline{\Delta Ct}_{PCRo}$ as:

$$\sigma_{\Delta\Delta Ct} = \sqrt{\left(\sigma_{\ \overline{\Delta Ct}_{PCRu}}\right)^2 + \left(\sigma_{\ \overline{\Delta Ct}_{PCRo}}\right)^2}$$

The final uncertainty in the value of fraction uncleaved gate in Eq. 1 is obtained from:

$$\sigma = \text{ Fraction uncleaved} * \ln(2) * \sigma_{\Delta\Delta\overline{Ct}}$$

***Validity of data analysis method:*** The base of 2 in Eq. 1 assumes 100% primer amplification efficiency, which may not always be the case in practice. Using dilutions of U (Supplementary Figure S3), we measured primer amplification efficiencies for the six genetic contexts in our Ro test set, all of which use the same primers. We found amplification efficiencies varied between 94% and 104% across samples and independent replicates with an average of 100% (Supplementary Figure S9). Based on these results, we assumed the overall amplification efficiency for these primers was near 100% and Eq. 1 was valid to use.

Additionally, for the ΔΔCt analysis in Eq. 2 to be valid, three criteria need to be met: (1) the PCRu and PCRo primers should have similar amplification efficiencies, (2) the reverse transcription efficiency, and (3) PCR amplification efficiencies of the PCRo need to be the same for the uncleaved gate, cleaved gate, and the uncleaved control (U) RNA. If these criteria are met, then any differences in Ct values between G and U samples in PCRo can be attributed to differences in total RNA concentration between the two samples. Shifting $\Delta Ct_{PCRu}$ by $\Delta Ct_{PCRo}$ accounts for this difference. If these criteria are not met, then $\Delta Ct_{PCRo}$ could erroneously shift the ΔΔCt value even if the same concentration of U and G are used. Because the U control transcript only differs from G by a single base mutation, we expected these two samples to have similar reverse transcription and amplification efficiencies for a given primer set.

To explore whether the above criteria were met for our system, we prepared a series of dilutions of U1, C1, and U1+C1 and evaluated primer amplification efficiencies with reverse transcription at 37 °C. For the Ro ribozyme, primer efficiencies were between 92% and 105% for all dilution series (Supplementary Figure 7A). Further, U1, G1, and C1 RNAs had similar Ct values for PCRo (Supplementary Figure 7B), suggesting reverse transcription efficiencies are similar for U, G, and C RNAs. For the Rh ribozyme, primer amplification efficiencies were near 100%, but the Ct values of U and G for PCRo differed greatly with reverse transcription at 37 °C (Supplementary Figure S8). This resulted in a fraction uncleaved for G2h that was lower than expected (Supplementary Figure S10). Repeating this experiment with a 50 °C reverse transcription step alleviated these issues (Supplementary Figures S8 and S10). These results suggest there is a difference in reverse transcription efficiency between the U2h and G2h at 37 °C, which alters ΔΔCt more than can be attributed to a difference in concentration alone.

#### 2.3 RT-qPCR controls and melt curve analysis

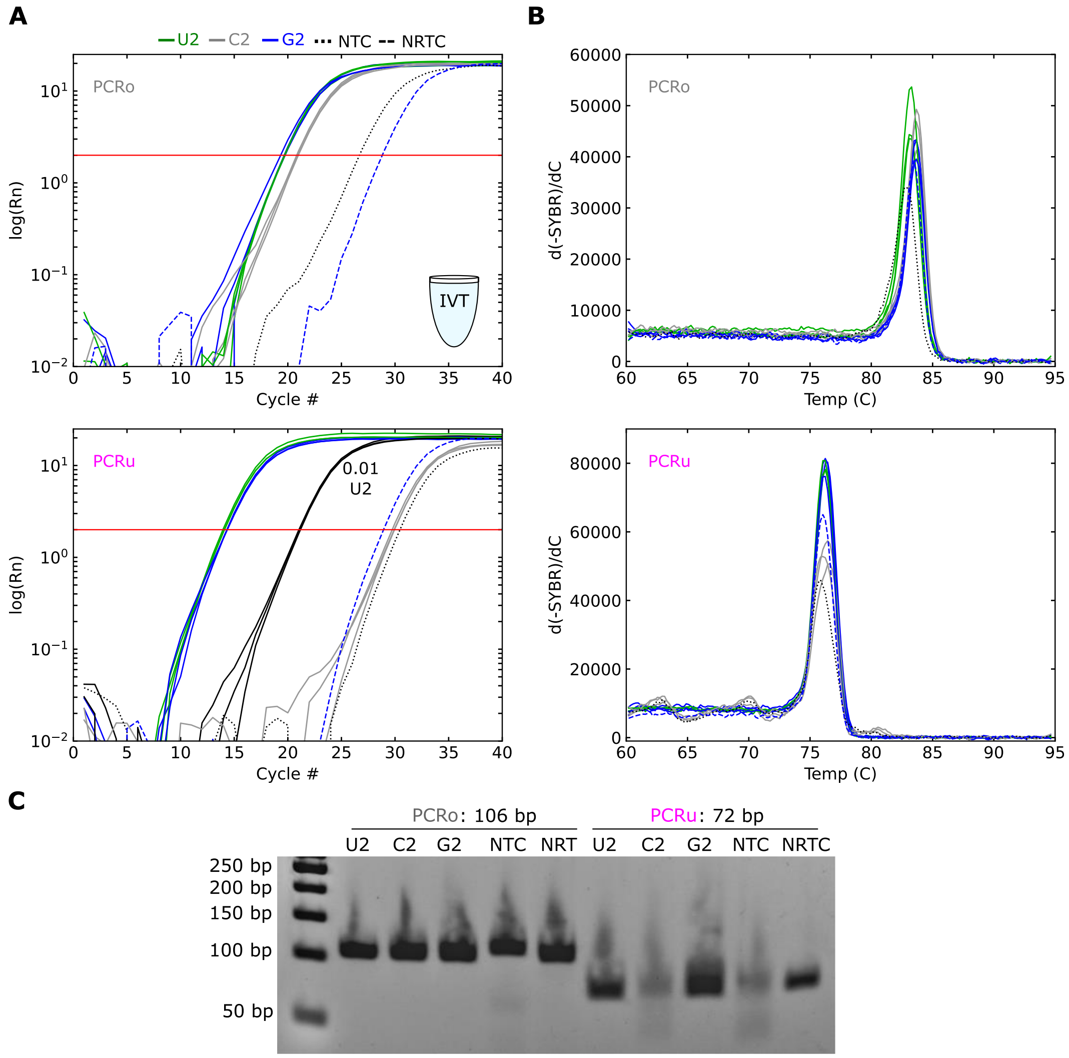

**Supplementary Figure S4:** (**A**) Normalized amplification curves for Replicate 2 of IVT G2 (also shown in Supplementary Figure S9). Lines of the same color represent technical replicates (N=3). The red horizontal lines indicate the threshold value used in this experiment. The solid black curves for PCRu are replicates with 0.01 U2 relative to the green U2 curves. The dashed blue curves represent controls in which reverse transcriptase was left out of the gate samples (NRTC). The dotted black curves represent controls in which the RNA was left out of the reactions (NTC). Although amplification is seen for these controls, the Ct values are nearly 10 cycles or more greater than Ct values for the experimental samples, indicating our results are not the result of DNA contamination or nonspecific primer amplification. Note for PCRu, the cleaved control sample (C1) does not amplify more than the background negative controls as expected. (**B**) Melt curves for the samples in (A). (**C**) Native gel electrophoresis results of the RT-qPCR products indicated above the gel. RNA was produced by *in vitro* transcription and gel electrophoresis was conducted with a 4% agarose E-gels prestained with SYBR Safe.

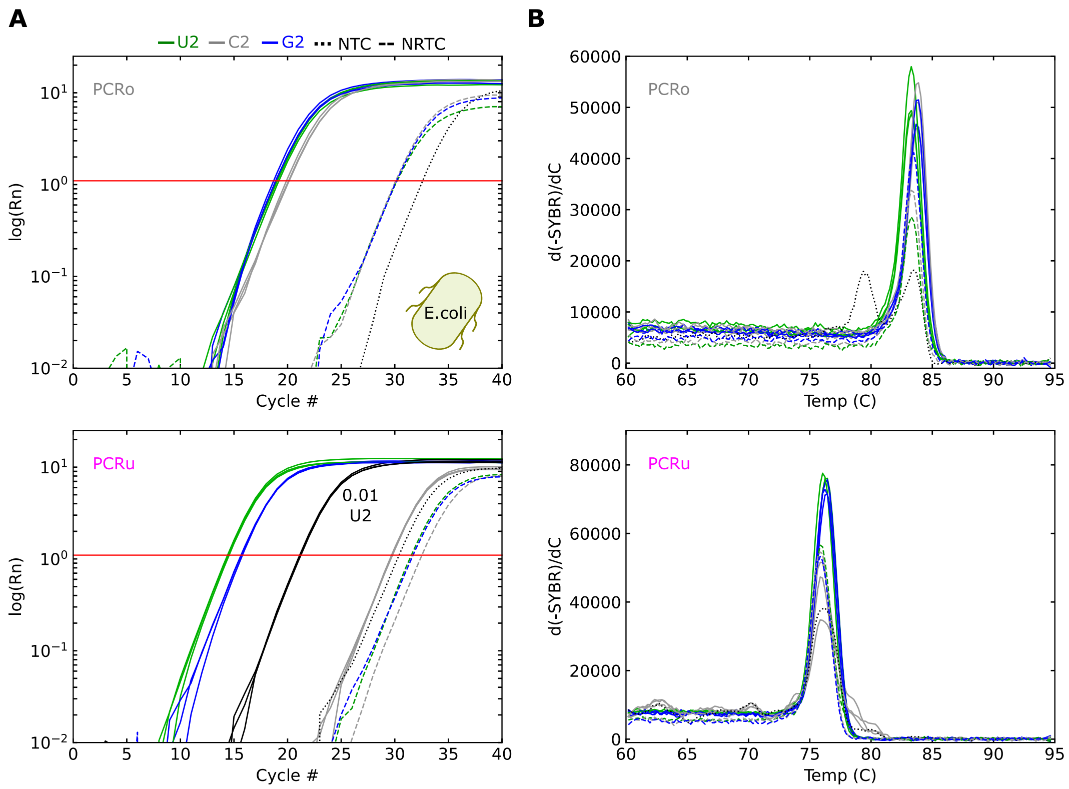

**Supplementary Figure S5:** (**A**) Amplification curves for Replicate 1 of G2 produced in *E. coli* BL21* (DE3) (also shown in Supplementary Figure S11). Lines of the same color represent technical replicates (N=3). The red horizontal lines represent the threshold value used in this experiment. The solid black curves for PCRu are replicates with 0.01 U2 relative to the green U2 curves. The dashed blue curves represent controls in which reverse transcriptase was left out of the gate samples (NRTC). The dotted black curves represent controls in which the RNA was left out of the reactions (NTC). Although amplification is seen for these controls, the Ct values are nearly 10 cycles or more greater than Ct values for the experimental samples, indicating our results are not the result of DNA contamination or nonspecific primer amplification. Note for PCRu, the cleaved control sample (C2) does not amplify more than the background negative controls as expected. (**B**) Melt curves for the samples in (A).

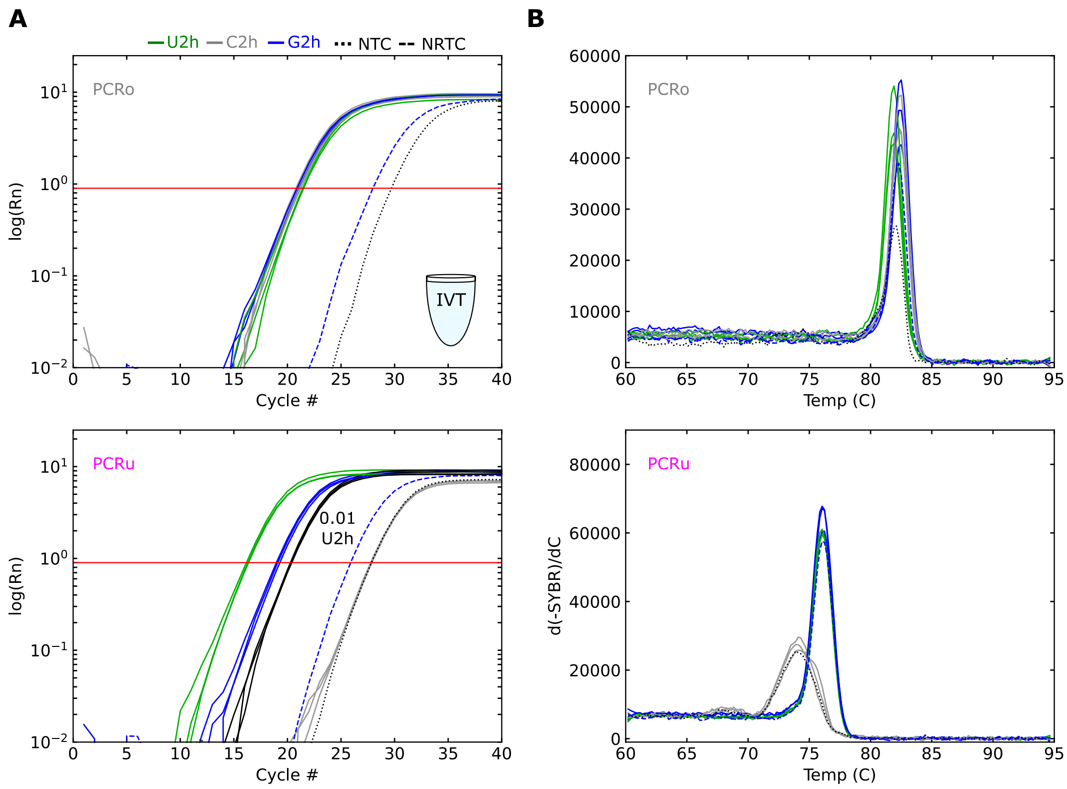

**Supplementary Figure S6:** (**A**) Normalized amplification curves for Replicate 2 of IVT G2h with reverse transcription conducted at 50 °C (also shown in Supplementary Figure S10). Lines of the same colore represent technical replicates (N=3). The red horizontal line represents the threshold value used in this experiment. The solid black curves for PCRu are replicates with 0.01 U2h relative to the green U2h curves. The dashed blue curves represent controls in which reverse transcriptase was left out of the gate samples (NRTC). The dotted black curves represent controls in which the RNA was left out of the reactions (NTC). Although amplification is seen for these controls, the Ct values are nearly 10 cycles or more greater than Ct values for the experimental samples, indicating our results are not the result of DNA contamination or nonspecific primer amplification. Note for PCRu, the cleaved control sample (C2h) does not amplify more than the background negative controls as expected. (**B**) Melt curves for the samples in (A).

#### 2.4 Primer efficiency analysis

In addition to the data below, primer amplification efficiency experiments were conducted for each replicate of each gate sequence in this study.

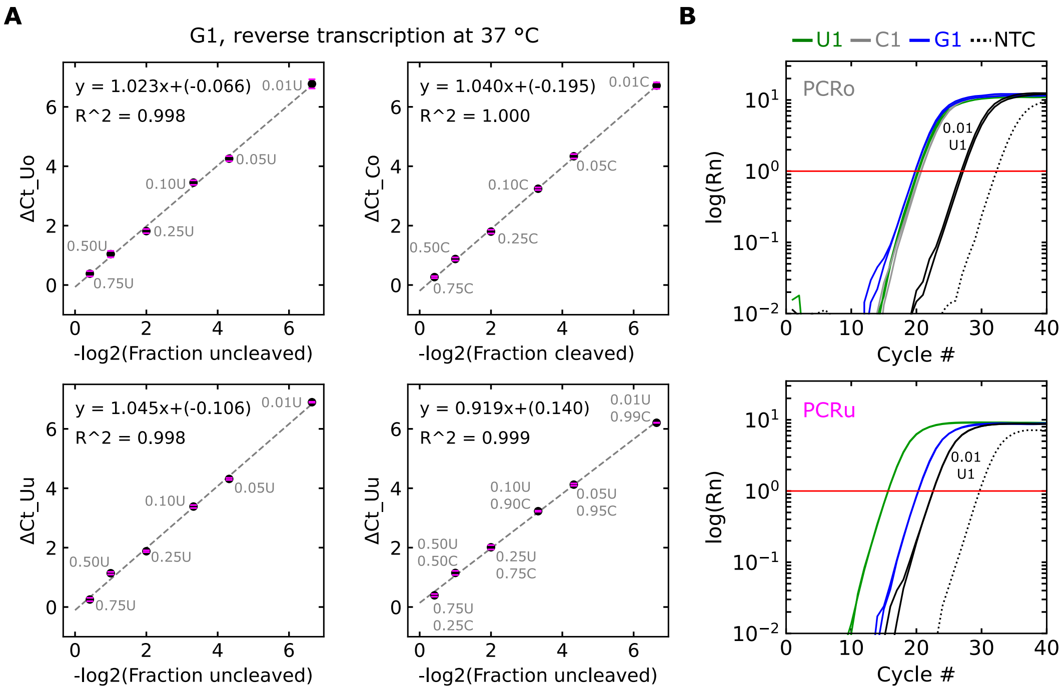

**Supplementary Figure S7:** (**A**) Testing PCR amplification efficiencies of PCRu and PCRo primers for G1 uncleaved (U1) and cleaved (C1) control RNAs. Subscripts u and o indicate PCRu and PCRo primers, respectively. PCRo has similar amplification efficiency for both U1 and C1 (top left and top right, respectively), suggesting there isn't a bias in amplification between the two types of gate products. PCRu has >90% amplification efficiency (100*slope of fit lines) with the U1 control RNA both with and without C1 present (bottom left and bottom right, respectively), suggesting the presence of C1 does not interfere with PCRu. Pink error bars indicate ± one standard deviation of the mean from two technical replicates. (**B**) Normalized amplification curves for PCRo and PCRu. The solid black curves are replicates with 0.01 U1 relative to the green U1 curves. NTC is a no template control. Lines of the same color represent technical replicates (N=2). The red horizontal lines indicate the threshold value used in the experiments. Note for PCRo that U1, C1, G1 cross the threshold at similar cycle numbers, consistent with these RNAs having similar reverse transcription efficiencies.

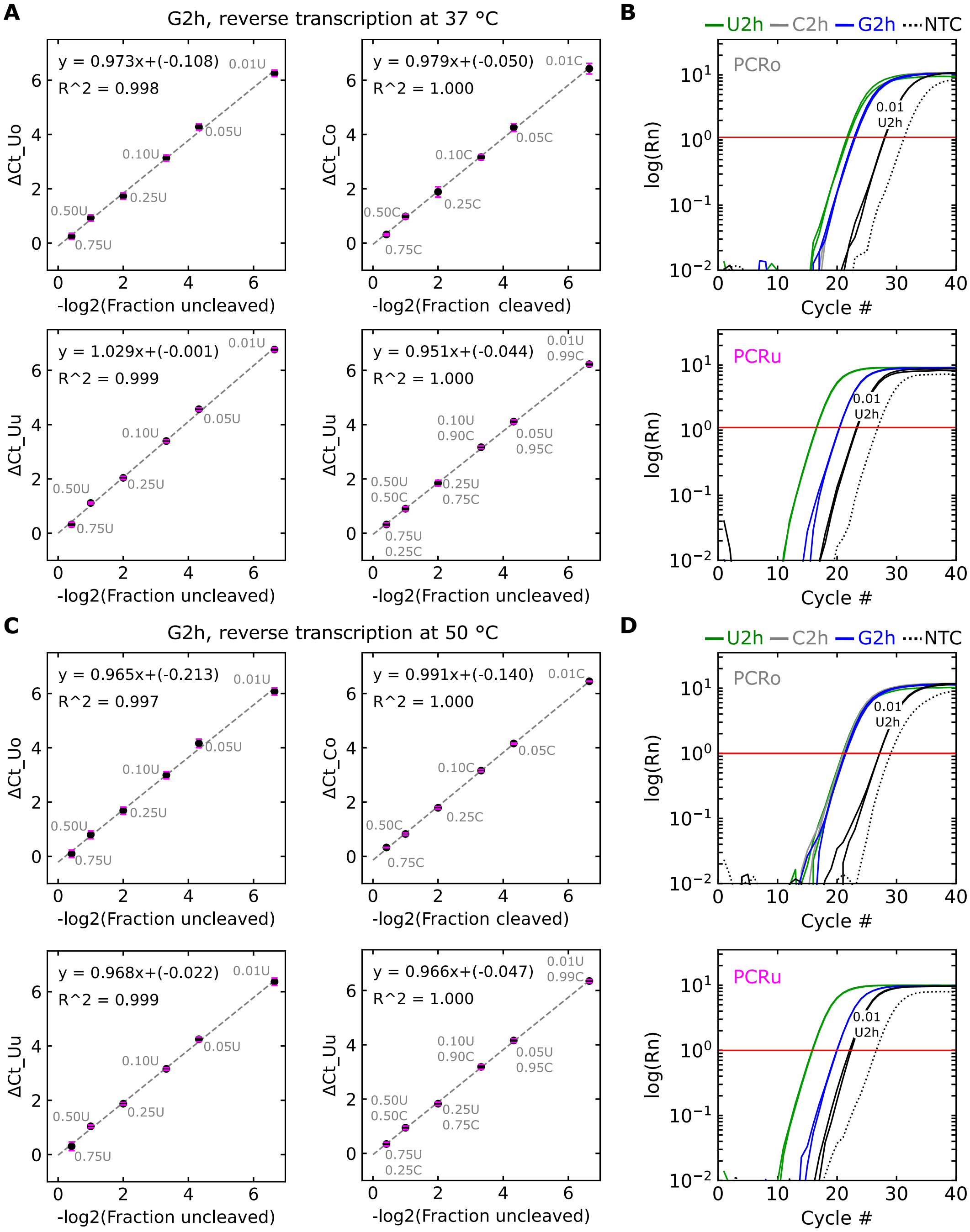

**Supplementary Figure S8:** Analysis of Rh primers with reverse transcription (RT) at 37 °C (**A**,**B**) or 50 °C (**C**,**D**). (**A**,**C**) Primer amplification efficiencies for PCRo (top panel) and PCRu (bottom panel) for dilutions indicated within the plots. Both primer sets have >95% amplification efficiency at irrespective of reverse transcription temperature. Pink error bars indicate ± one standard deviation of the mean from two technical replicates. (**B**,**D**) Normalized amplification curves for PCRo and PCRu with different reverse transcription temperatures. At 37 °C PCRo crosses the threshold at different cycles for U2h and G2h but at 50 °C these samples cross the threshold at similar cycles. This suggests the reverse transcription efficiency of U2h and G2h are different at 37 °C but not at 50 °C. Experiments at 37 °C or 50 °C were conducted with RNA from the same IVT and blocking oligo preparation. The solid black curves are replicates with 0.01 U2h relative to the green U2h curves. NTC is a no template control. Lines of the same color represent technical replicates (N=2). The red horizontal lines indicate the threshold value used in the experiments.

#### 2.5 Amplification curves and primer efficiency plots for all samples

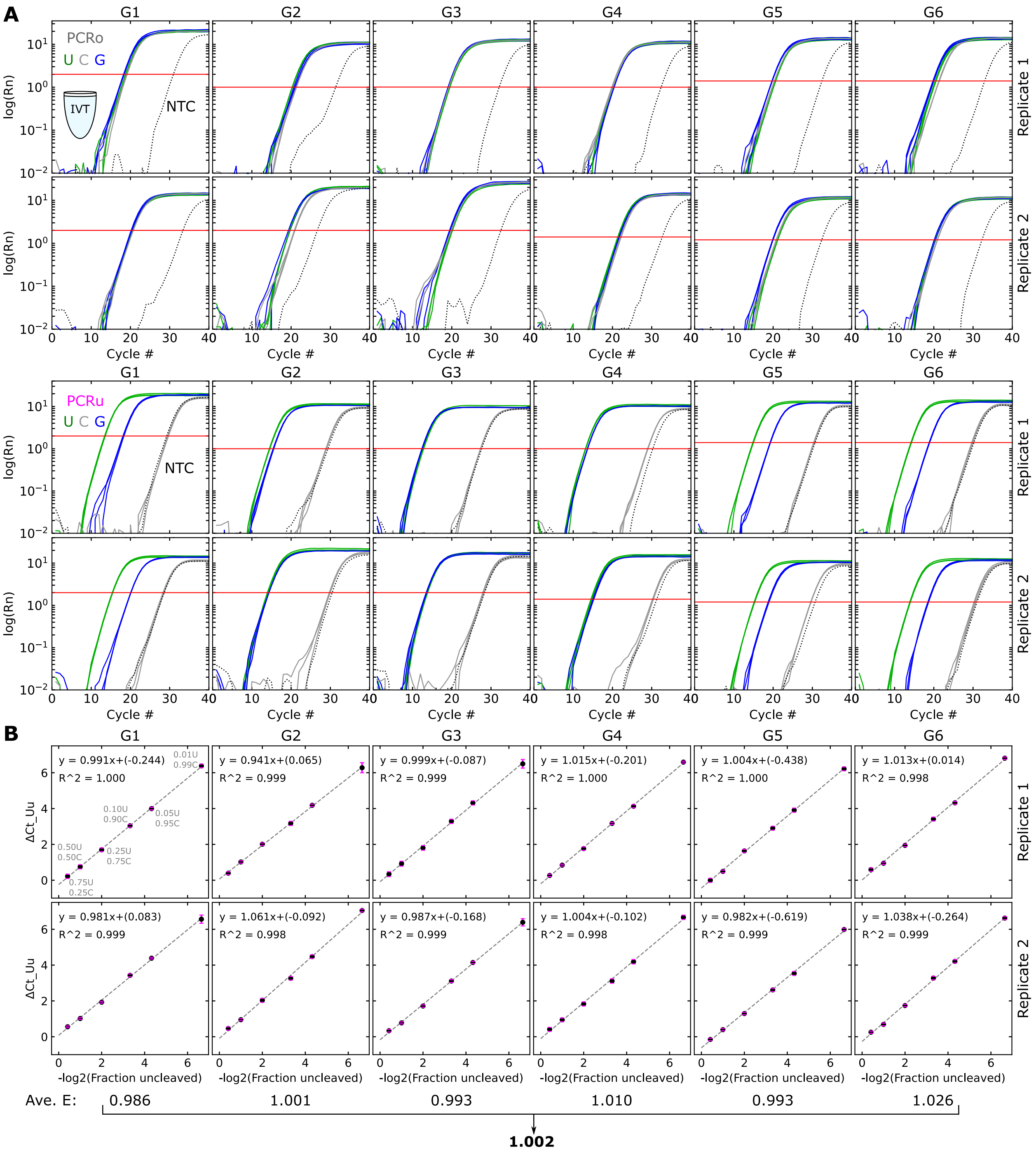

**Supplementary Figure S9:** Normalized amplification curves (**A**) and PCRu primer amplification efficiency curves (**B**) for all six validation RNA sequences produced by IVT. In (A), multiple lines of the same color of lines represent three technical replicates. The red horizontal lines indicate the threshold value for each experiment. The dotted black curves represent controls in which the RNA was left out of the reactions (NTC). In (B), dilutions of U were prepared with C (Supplementary Figure S3C). Pink bars indicate standard deviation of technical replicates. Note the average PCRu amplification efficiency across all gates and replicates is nearly 1 (100%).

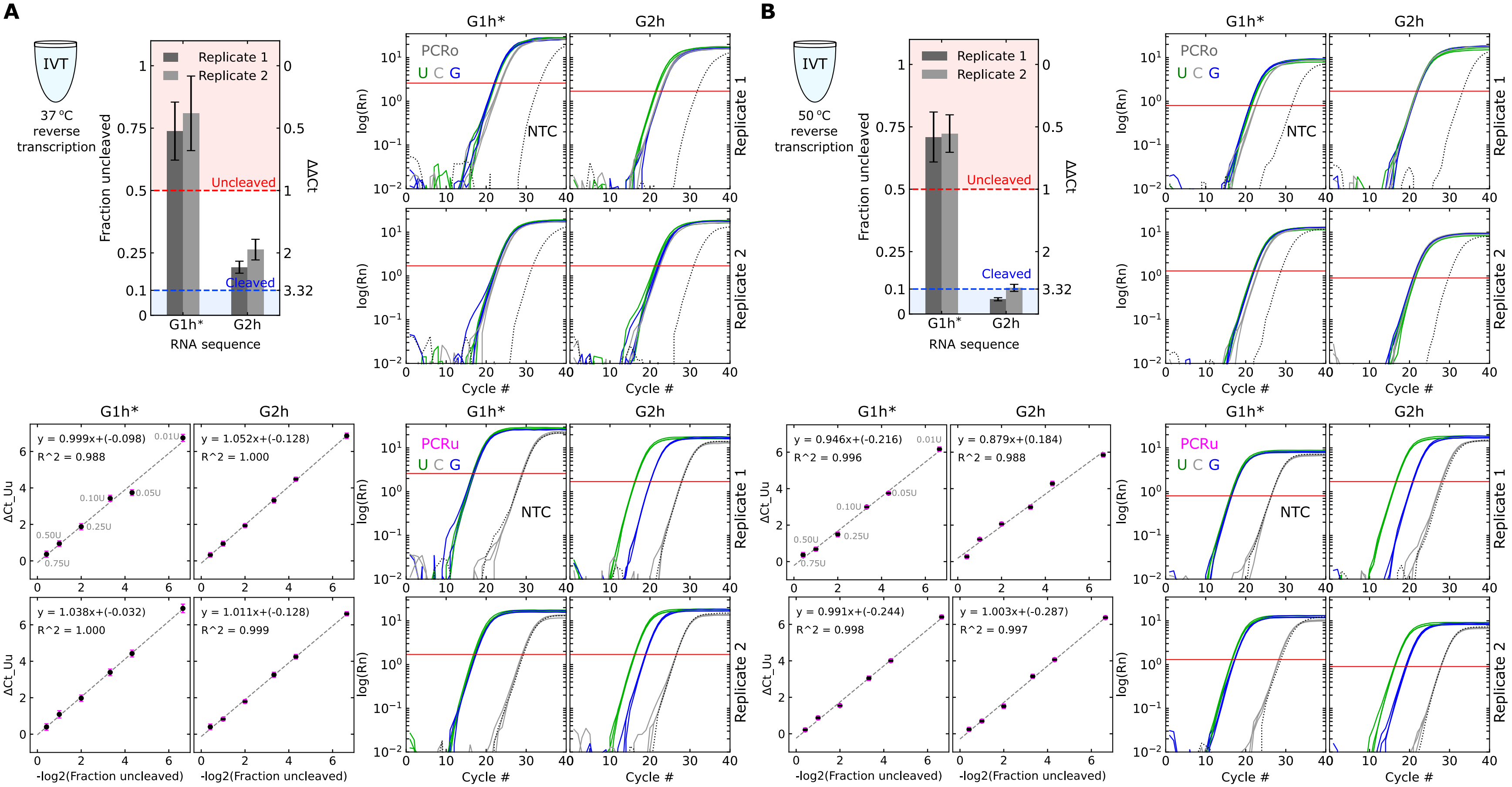

**Supplementary Figure S10:** Fraction uncleaved (top left quadrant), PCRu primer amplification efficiency curves (bottom left quadrant), and normalized amplification curves (right quadrants) for the two IVT Rh gates with 37 °C (**A**) or 50 °C (**B**) reverse transcription steps. A difference in PCRo Ct values between U and G samples likely leads to a higher than expected fraction uncleaved for G2h at 37 °C. Experiments at both temperatures were done with RNA from the same IVT and blocking oligo preparation, so it is unlikely that differences seen in the PCRo Ct values for U and G are due to incorrect concentration measurements or mixing of the U and G samples. For the primer amplification efficiency curves, U dilutions were done in RNA storage solution without C present (Supplementary Figure S3E). Pink bars indicate standard deviation of three technical replicates. In the amplification curve plots, multiple lines of the same color represent three technical replicates. The red horizontal lines indicate the threshold value for each experiment. The dotted black curves represent controls in which the RNA was left out of the reactions (NTC). The fraction uncleaved results in (B) are also shown in Figure 3 of the main text.

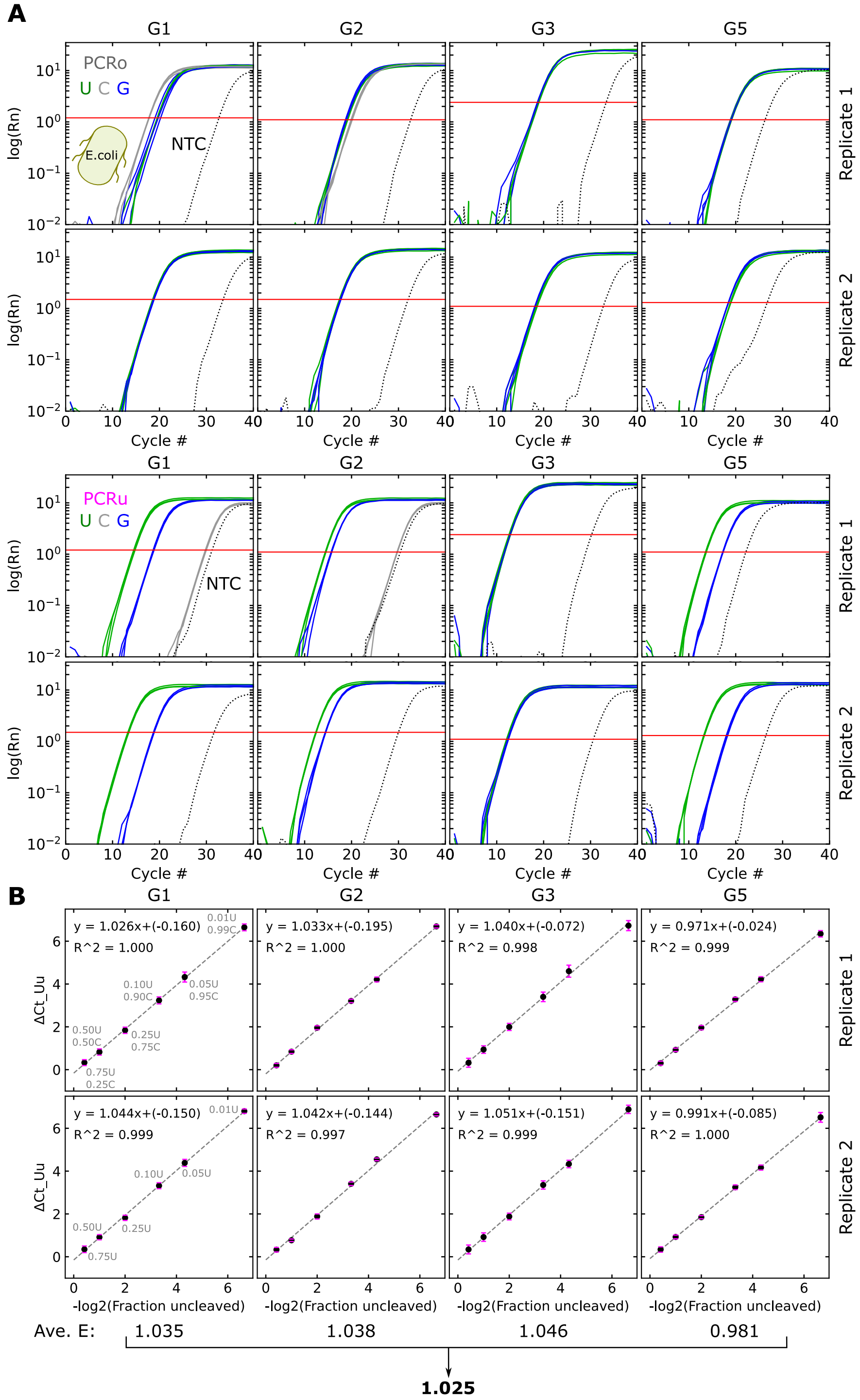

**Supplementary Figure S11:** Normalized amplification curves (**A**) and PCRu primer amplification efficiency curves (**B**) for all Ro gates tested in *E. coli* BL21* (DE3). In (A), multiple lines of the same color of lines represent three technical replicates. The red horizontal lines indicate the threshold value for each experiment. The dotted black curves represent controls in which the RNA was left out of the reactions (NTC). Replicate 1 of G1 and G2 had the C control RNA but the rest of the experiments did not include this control. In (B), dilutions of U were prepared with C only for Replicate 1 of G1. The rest of the U dilutions were prepared with RNA storage solution (Supplementary Figure S3E). Pink bars indicate standard deviation of three technical replicates. Note the average PCRu amplification efficiency across all gates and replicates is nearly 1 (100%).

#### 3 Benchmarking metrics for comparing gel electrophoresis and RT-qPCR results

To validate our RT-qPCR method, we selected six validation RNA sequences (Supplementary Section 1) for which we had orthogonal *in vitro* measurements of cleavage (Figure 1C of the main text). From denaturing gel electrophoresis measurements, three of these RNAs (G1, G5, G6) showed the majority of the RNA cleaved and the other three RNAs (G2, G3, G4), the majority of the RNA did not cleave.

To access whether our RT-qPCR could recapitulate these results, we developed as set of benchmarking metrics that considered the limitations of the both the gel electrophoresis measurements and RT-qPCR measurements. For the RNAs that are a majority uncleaved when produced by IVT (G2, G3, G4), the fraction uncleaved should be >0.75 based on gel electrophoresis results (Figure 1C and Supplementary Figure S12A). However, this corresponds to a ΔΔCt of <0.5 when measured by RT-qPCR (Supplementary Figure S12B), which is generally considered to be within the accepted range for technical replicates (4). In other words, RT-qPCR measurements of <0.5 ΔΔCt are susceptible to experimental noise and will have large uncertainty. Thus, we selected a ΔΔCt <1 as our metric of validation of RT-qPCR measurements for RNAs G2, G3, and G4 (red dashed line and red shaded region in Supplementary Figure S12B). This corresponds to a fraction uncleaved of >0.5, *i.e.*, a majority of the RNA, and should be measurable with a high degree of confidence with RT-qPCR. For the RNAs that are a majority uncleaved when produced by IVT (G1, G5, G6), the fraction uncleaved should be <0.10 based on gel electrophoresis results (Figure 1C and Supplementary Figure S12A). Our RT-qPCR measurements in this regime should be much less susceptible to experimental noise because the corresponding ΔΔCt values are >3. However, with our gel electrophoresis measurements it is difficult to quantitate the fraction of uncleaved RNA below 0.1. Therefore, we selected ΔΔCt >3.32, corresponding to fraction uncleaved of <0.1, as our metric of validation of RT-qPCR measurements for G1, G5, and G6. None of the six validation RNAs should result in 1<ΔΔCt <3.32 (Supplementary Figure S12B), as this would correspond to fractions uncleaved inconsistent with the majority of RNA being uncleaved or cleaved as we measured with gel electrophoresis.

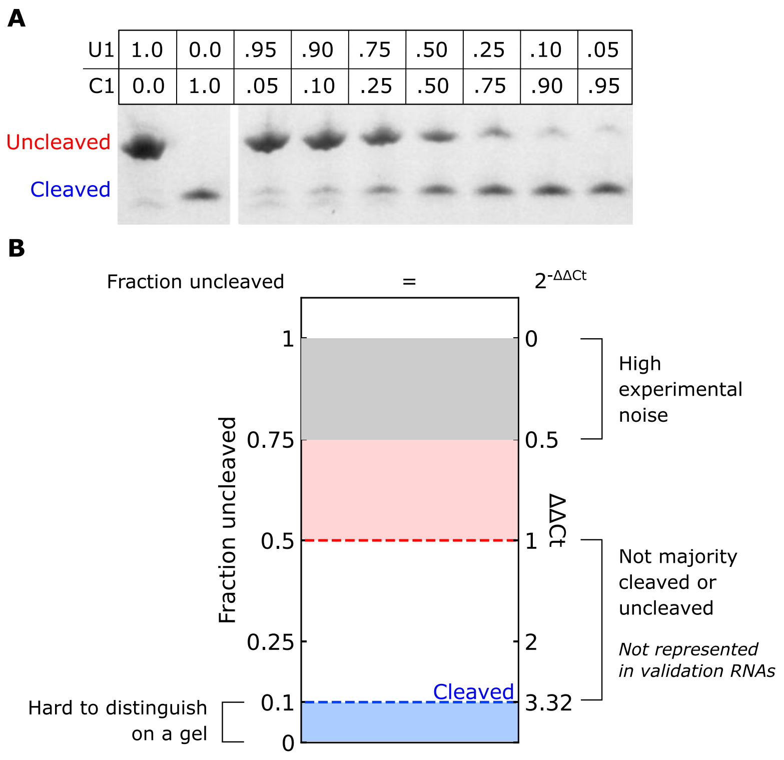

**Supplementary Figure S12:** Rational for the benchmarking metrics developed for (**A**) Denaturing gel electrophoresis results for different fractional mixtures of U1 and C1 RNAs. The fractional mixtures were obtained by *in vitro* transcribing the ratios of U1 and C1 DNA templates specified above the gel (total template concentration of 25 nmol/L across all samples). The DNA templates were digested with DNase I prior to electrophoresis. (**B**) Relationship between fraction uncleaved RNA and ΔΔCt values measured with RT-qPCR. Shaded regions designate the areas the six validation RNAs from Figure 1 should fall within.

### 4 Analysis of ribozyme cleavage during sample preparation and RT-qPCR

Ribozyme cleavage results in two RNA products. The higher molecular weight cleavage product is labeled cleaved and corresponds to the region of the transcript downstream (DS) of the cleavage site. The lower molecular weight cleavage product corresponds to the region of the transcript upstream (US) of the cleavage site and is labeled as such when shown below. On most of the gels in this study, only the downstream cleavage product is shown, as the lower molecular weight product often ran off the gel.

#### 4.1 Ribozyme cleavage induced during sample preparation and reverse transcription

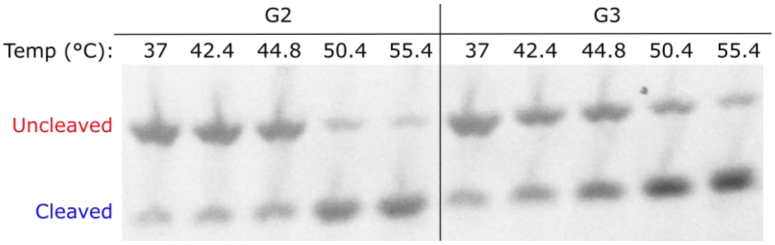

**Supplementary Figure S13:** Higher temperatures can induce cleavage in gates that do not cleave at 37 °C. (**A**,**B**) Denaturing gel electrophoresis results of different gates after 15 min incubation at the temperatures indicated above the plots. RNA was prepared by 30 min *in vitro* transcription at 37 °C followed by a 30 min DNase I digestion at 37 °C. RNA was then incubated for 15 min at the indicated temperatures. Formamide with EDTA was added to quench the cleavage reaction prior to electrophoresis (Methods). DNA templates were at 25 nmol/L and T7 RNAP was at 1 U/µL during *in vitro* transcription.

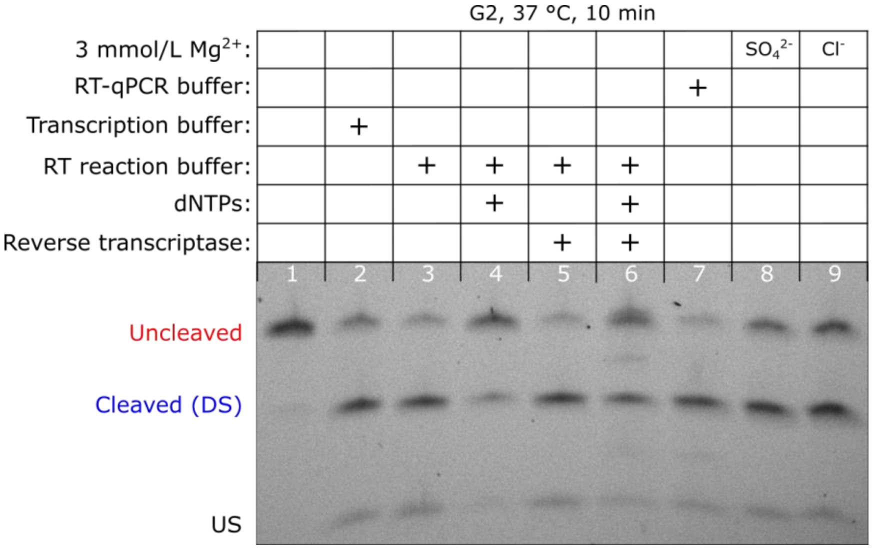

**Supplementary Figure S14:** Denaturing gel electrophoresis results of IVT G2 RNA after incubation in different conditions. IVT G2 RNA was added to a final concentration of 7.5 ng/ µL. Each incubation reaction contained a volume fraction of 45 % RNA storage solution. In the relevant samples, dNTPs were added to a final concentration of 0.75 mmol/L each (3 mmol/L total). Magnesium with two different counter ions was added to the samples shown Lanes 8 and 9. The RT-qPCR buffer came from a SuperScript III Platinum SYBR Green One-Step Kit (ThermoFisher, 11736059) with the following 1x composition [3 mmol/L $MgSO_4$, 0.4 mmol/L of each dNTP, and a propriety mix of buffer, SYBR Green I, and stabilizers]. The transcription buffer came with T7 RNAP (ThermoFisher, EP0113) with the following 1x composition [40 mmol/L tris-HCl (pH 7.9), 6 mmol/L $MgCl_2$, 10 mmol/L dithiothreitol, 10 mmol/L NaCl, and 2 mmol/L spermidine]. The reverse transcriptase enzyme and RT reaction buffer came from SuperScript III Platinum Two-Step qRT-PCR Kit with SYBR Green Kit (ThermoFisher, 11735-032) with the buffer having the following 1x composition [6 mmol/L $MgCl_2$ and 0.75 mmol/L of each dNTP in a propriety buffer].

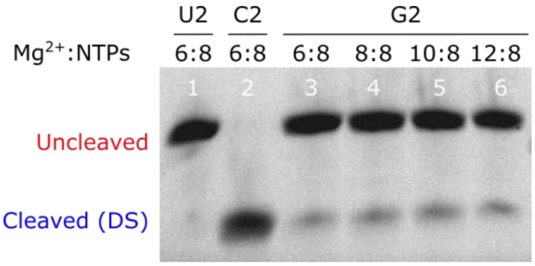

**Supplementary Figure S15:** Denaturing gel electrophoresis results showing that addition of magnesium after transcription but before RNA purification does not induce cleavage. RNAs were transcribed *in vitro* at 37 °C for 1 h, DNA templates were degraded with DNase I for 20 min, then additional $MgCl_2$ was added to lanes 4 to 6 and the samples were incubated for another 10 min before denaturing gel electrophoresis. The transcription reactions contained 6 mM $MgCl_2$ and 2 mM of each NTP (8 mM total). The ratios above the gel indicate the concentration of $MgCl_2$ to total NTPs during the final 10 min incubation.

#### 4.2 Analysis of the blocking oligo to prevent ribozyme cleavage during reverse transcription

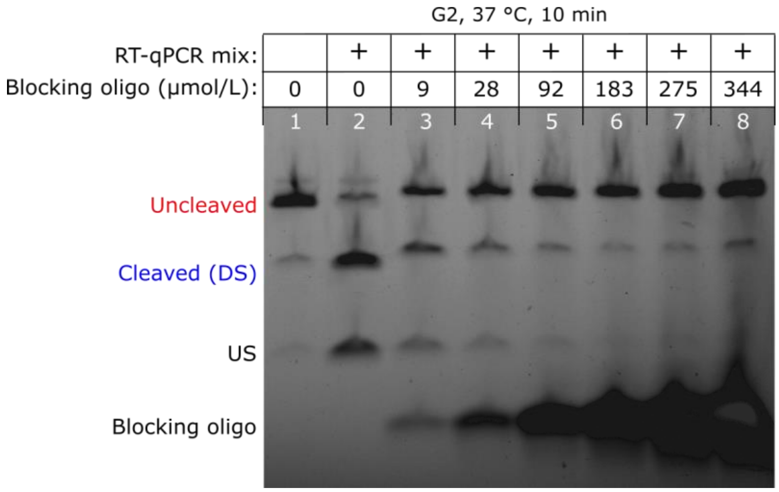

**Supplementary Figure S16:** Denaturing gel electrophoresis results of blocking oligo titrations with IVT G2 RNA. The concentrations of blocking oligo shown above the gel were mixed with ≈600 nmol/L of G2 RNA and then annealed. After annealing, the samples were mixed with an equal volume of RT-qPCR mix when indicated above the gel and subsequently incubated at 37 °C for 10 min. Formamide and EDTA were added to quench the cleavage reaction prior to electrophoresis (Methods). Blocking oligo concentrations of 92 µmol/L and above prevent cleavage during the incubation in RT-qPCR mix. In this experiment, 92 µmol/L is an ≈150-fold molar excess of blocking oligo to gate RNA. In most experiments, an ≈175-fold molar excess of blocking oligo was used. Such a large excess of blocking oligo may be required because there is no free magnesium present during the anneal or because of the strong secondary structure of the HDV ribozyme (3, 4).

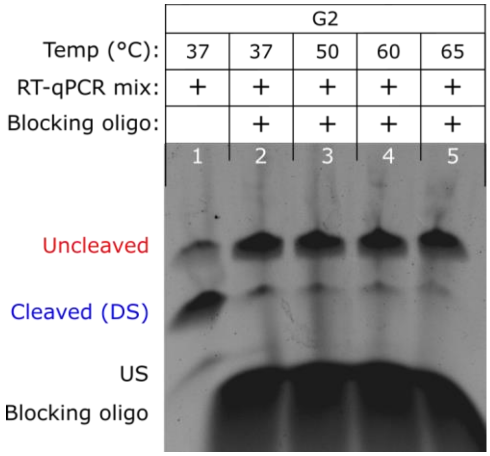

**Supplementary Figure S17:** Denaturing gel electrophoresis results of IVT G2 RNA with blocking oligo at increasing temperatures. 157 µmol/L of blocking oligo was mixed with ≈600 nmol/L of IVT G2 RNA and then annealed. After annealing, the samples were mixed with an equal volume with RT-qPCR mix and subsequently incubated at the temperatures specified above the gel for 10 min. Formamide and EDTA were added to quench the cleavage reaction prior to electrophoresis (Methods).

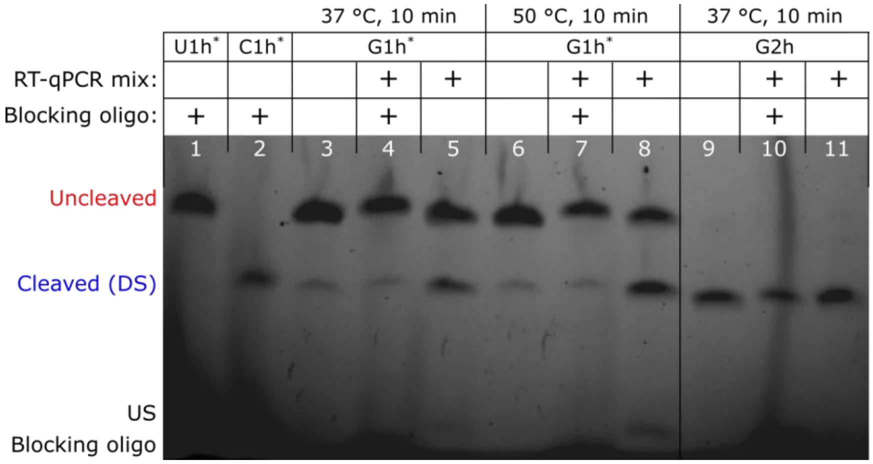

**Supplementary Figure S18:** Denaturing gel electrophoresis results of blocking oligo tests for gates with the CPEB-3 (Rh) ribozyme. When present, the blocking oligo concentration was 157 µmol/L. RNA was at ≈600 nmol/L for all samples. Lane 3 to lane 5 are also presented in Figure 3B of the main text. The blocking oligo prevents cleavage of G1h* at both 37 °C and 50 °C.

#### 5 References

1. Schaffter,S.W., Wintenberg,M.E., Murphy,T.M. and Strychalski,E.A. (2023) Design Approaches to Expand the Toolkit for Building Cotranscriptionally Encoded RNA Strand Displacement Circuits. *ACS Synth. Biol.*, **12**, 1546–1561.

2. Kim,M.W., Sun,G., Lee,J.H. and Kim,B. (2018) Development of Quenching-qPCR (Q-Q) assay for measuring absolute intracellular cleavage efficiency of ribozyme. *Analytical Biochemistry*, **550**, 27–33.

3. Vlková,M., Morampalli,B.R. and Silander,O.K. (2021) Efficiency of the synthetic self-splicing RiboJ ribozyme is robust to cis- and trans-changes in genetic background. *MicrobiologyOpen*, **10**, e1232.

4. Bustin,S.A. (2004) A-Z of Quantitative PCR International University Line.

5. Taylor,J.R. (2022) An Introduction to Error Analysis: The Study of Uncertainties in Physical Measurements University Science Books.
