## Supplementary Methods for "Prevention of ribozyme catalysis through cDNA synthesis enables accurate RT-qPCR measurements of context-dependent ribozyme activity"

### 1 PCR amplification of DNA

Materials

- Bio-Rad T-100 Thermal cycler (BioRad, 1861096)
- DeNovix D-11 Series Spectrophotometer
- 200 µL PCR tubes (USA Scientific, 1402-4700)
- Nuclease-Free Water (Ambion AM9938)
- 2x Phusion High-Fidelity Master Mix (ThermoFisher, F531L)
- QIAquick spin PCR Purification Kit (Qiagen, 28104)
- DNA gene fragments (ordered as eBlocks at 10 ng/µL in IDTE buffer from IDT)
- DNA oligonucleotides (ordered as standard desalted from IDT and prepared at 100 µmol/L stocks in water)

Protocol

1) Prepare the master mix below and then add to the 200 µL PCR tube containing the eBlock DNA
    a. For IVT reactions, use T7fwd / T7rev as the two primers. For preparation of inserts for plasmids, use ga4_fwd / pET_ds_rev as the two primers (Supporting File S3)

| Component | Final volume or concentration |
|---|---|
| 2x Phusion MM | 37.5 µL |
| Fwd primer | 0.5 µmol/L |
| Rev primer | 0.5 µmol/L |
| Ambion water | X µL to 74.85 µL total |

2) Add 0.15 µL of 10 ng/µL eBlock DNA into each PCR tube containing master mix, pipette mix
3) Add samples to a thermocycler and execute the following protocol with the lid temperature set to 105 °C

| Temp | Time | Repeat |
|---|---|---|
| 98 °C | 5 min | 1x |
| 98 °C | 30 s | 30x |
| 60 °C | 30 s | 30x |
| 72 °C | 30 s | 30x |
| 72 °C | 3 min | 1x |
| 4 °C | hold | |

4) Run a PCR clean up with a QIAquick Spin PCR Purification Kit
    a. Add 375 µL of Buffer PB to a spin column
    b. Add all 75 µL of the PCR sample to the spin column containing Buffer PB and pipette mix thoroughly (10 to 20 times)
    c. Spin column at 13,000 rpm for 1 min, discard flow through
    d. Add 750 µL of Buffer PE to the spin column
    e. Spin column at 13,000 rpm for 1 min, discard flow through

f. Spin column at 13,000 rpm for 1 min to remove any last Buffer PE, discard flow through
g. Place spin column insert into a fresh collection tube
h. To elute the PCR product, add 50 µL of Buffer EB to the center of the spin column
i. Let the column sit for 1 min to 2 min at room temperature
j. Spin column at 13,000 rpm for 1 min, discard spin column

5) Measure the concentration of the purified PCR products with A260 of undiluted samples on a DeNovix D-11 Series Spectrophotometer

### 2 RNA production, purification, and protection from IVT

Materials

- Bio-Rad T-100 Thermal cycler (BioRad, 1861096)
- Qubit-4, Fluorometer
- 200 µL PCR tubes (USA Scientific, 1402-4700)
- T7 RNA polymerase (RNAP) and associated 5x transcription buffer (ThermoFisher, EP0113)
- Ribonucleotide triphosphates (NTPs) (ThermoFisher, R0481)
  - These come as separate tubes at 100 mmol/L each
  - For IVT reactions, equal volumes of each NTP is mixed to produced a solution with 25 mmol/L of each of the four NTP types
- 0.5 mol/L ethylenediaminetetraacetic acid (EDTA), pH 8.0 (Invitrogen, AM9260G)
- THE RNA Storage Solution, (ThermoFisher, AM7001)
- RNase Away Decontamination Reagent, 250 mL (ThermoFisher, 10328011)
- Nuclease-Free Water (Ambion, AM9938)
- TE, pH 8.0, RNase-free (ThermoFisher, AM9849)
- DNase I HC (50 U/µL), RNase-free and associated 10x DNA Digestion Buffer (ThermoFisher, EN0523)
- RNA Binding Buffer (ZymoResearch, R1013)
- RNA Wash Buffer (ZymoResearch, R1003)
- DNA templates for transcription (prepared by PCR in Section 1)
- Blocking oligonucleotide (prepared at a stock concentration of 1000 µmol/L in water)
- Zymo-Spin IC columns (ZymoResearch, C1004-50)
- Zymo collection Tubes, (ZymoResearch, C1001-50)
- Qubit RNA HS Buffer Kit (ThermoFisher, Q32852)
- Formamide (Millipore Sigma, S4117)

Protocol

*RNA production from IVT:*

1) For IVT reactions, prepare the following reaction mix as below in 200 µL PCR tubes
   a. For most experiments, a master mix was prepared with all components other than the DNA templates and DNA templates were added to tubes after distributing the master mix
   b. For U2h and G2h, which had low transcription efficiency due to only a single G at the 5' end of the transcripts, the DNA template concentration was increased to 100 nmol/L and T7 RNAP concentration was increased to 10 U/µL

| **Component** | **Final volume or concentration** |
|---|---|
| Ambion water | X µL to 25 µL total |
| 5x txn buffer | 1x (5 µL for 25 µL reaction) |
| NTPs (25 mmol/L each) | 2 mmol/L each (2 µL for 25 µL reaction) |
| DNA template (PCR product) | 25 nmol/L |
| T7 RNAP (200 U/µL) | 1 U/µL (0.13 µL for 25 µL reaction) |

2) Add samples to a thermocycler and incubate at 37 °C with the lid temperature set to 105 °C for 1 h

*RNA purification from IVT:*

3) In RNase free PCR tubes, add 2 µL DNase I HC (50 U/µL) to 3 µL 10x DNA Digestion Buffer and mix by gentle inversion
4) Add all 5 µL of this DNase I mixture to 25 µL of IVT RNA samples
5) Incubate at 37°C for 20 m
6) Add 60 µL RNA Binding Buffer supplemented with EDTA (final concentration 36 mmol/L)
7) Add 90 µL of 100% ethanol and mix pipette mix immediately
8) Transfer sample to the Zymo-Spin IC column and centrifuge for 30 s at 7,000 x g
9) Transfer the column into a new Zymo collection tube
10) Add 700 µL RNA Wash Buffer to the column and centrifuge for 30 sat 7,000 x g
11) Discard the supernatant
12) Add 400 µL RNA Wash Buffer to the column and Centrifuge for 1.5 min at 12,000 x g
13) Transfer the column carefully into an RNase free tube
14) Add 20 µL of sterile TE buffer directly to the column matrix, incubate for 5 min at room temperature and centrifuge for 2 min at 12,000 x g
15) Transfer to the fresh tube with 20 µL of THE RNA Storage Solution for prolonged storage
16) Dilute total RNA 1:5 by volume with THE RNA Storage Solution before quantitation with Qubit
17) Verify RNA concentration using Qubit with Qubit RNA HS Buffer Kit
18) The protocol can be paused at this stage and the RNA stored at -80 ºC

*RNA protection with blocking oligonucleotide:*

19) Transfer 120 ng of purified IVT RNA into a fresh PCR tube (calculate necessary volume according to Qubit quantitation)
20) Add 2.2 µL of blocking oligonucleotide (stock concentration 1000 µmol/L) and add RNA Storage Solution to reach a total volume of 14 µL
21) Transfer the sample to a thermocycler and run the following annealing protocol:

| **Temp (ramp)** | **Time** | **Repeat** |
|---|---|---|
| 90 ºC | 5 m | 1x |
| 90 ºC (-0.1 ºC) | 6 s | 99x |
| 80 ºC (-0.1 ºC) | 6 s | 99x |
| 70 ºC (-0.1 ºC) | 6 s | 99x |
| 60 ºC (-0.1 ºC) | 6 s | 99x |
| 50 ºC (-0.1 ºC) | 6 s | 99x |
| 40 ºC (-0.1 ºC) | 6 s | 99x |
| 30 ºC (-0.1 ºC) | 6 s | 99x |
| 20 ºC | hold | |

22) Verify RNA concentration using Qubit with Qubit RNA HS Buffer Kit
    a. We found that the presence of the blocking oligonucleotide would typically result in an increased concentration compared to our measurements of the RNA prior to blocking oligonucleotide addition. This could be up to four times higher than the nominal concentration of RNA we added to the blocking oligonucleotide reaction

### 3 Gel electrophoresis

Materials

- Bio-Rad T-100 Thermal cycler (BioRad, 1861096)
- E-Gel Power Snap-Electrophoresis Devise (Invitrogen, G8100)
- Gel Documentation System: FastGene FAST-Digi PRO (NIPPON EUROPE Genetics, GP-07LED)
- DNase I (2 U/µL), RNase-free (New England Biolabs, M0303S)
- 50 mmol/L $CaCl_2$ (prepared from $CaCl_2$ salt in DI water)
- Formamide (Millipore Sigma, S4117)
- 0.5 mol/L ethylenediaminetetraacetic acid (EDTA), pH 8.0 (Invitrogen, AM9260G)

Protocol

*For denaturing gel electrophoresis of IVT RNA without blocking oligonucleotide:*

1) Mix 0.5 µL of DNase I (stock at 2 U/µL) and 0.5 µL of 50 mmol/L $CaCl_2$ with 10 µL of the IVT samples described in Section 2 after the 1 h incubation 37 °C, pipette mix
2) Allow samples to incubate for 20 min at 37 °C in the thermocycler
3) Add 10 µL of ≈100 % formamide supplemented with 36 mmol/L of EDTA, pipette mix
4) Increase the temperature of the thermocycler to 85 °C and incubate the samples for 5 min at this temperature
5) Immediately load 20 µL of sample to a well in a 4% Agarose E-gel EX with SYBR Gold II
6) Run gel for 30 min on an E-Gel Power Snap-Electrophoresis Devise
    a. Select the 4% agarose gel setting in the device when setting up protocol
7) Allow gel to cool to room temperature and then image on a gel documentation system
    a. We found that gels taken for imaging right after electrophoresis were hot to the touch and did not image well until allowed to cool down

*For denaturing gel electrophoresis of IVT RNA with blocking oligonucleotide:*

1) Starting with RNA that has been annealed with blocking oligonucleotide (taken through Step 21 of Section 2), make the following mixture:

| **Component** | **Final volume** |
|---|---|
| SuperScriptR III RT/PlatinumR *Taq* Mix | 0.2 µL |
| 2X SYBRR Green Reaction Mix | 5 µL |
| Purified and blocked RNA | 4.8 µL |

2) Incubate samples for 10 min in thermocycler at desired temperature (37 ºC or 50 ºC)
3) Add 10 µL of ≈100 % formamide supplemented with 36 mmol/L of EDTA, pipette mix
4) Increase the temperature of the thermocycler to 85 °C and incubate the samples for 5 min at this temperature
5) Immediately load 20 µL of sample to a well in a 4% Agarose E-gel EX with SYBR Gold II

6) Run gel for 30 min on an E-Gel Power Snap-Electrophoresis Devise
    a. Select the 4% agarose gel setting in the device when setting up protocol
7) Allow gel to cool to room temperature and then image on a gel documentation system

### 4 Plasmid assembly and cloning

Materials

- Bio-Rad T-100 Thermal cycler (BioRad, 1861096)
- Eppendorf Eporator (Eppendorf, 4309000027)
- pETDuet-1 plasmid DNA (Millipore Sigma, 71146-3)
- Falcon 14 mL Round Bottom High Clarity PP Test Tube (Corning, 352059)
- 2x Phusion High-Fidelity Master Mix (ThermoFisher, F531L)
- SOC outgrowth medium (New England Biolabs, B9020S)
- QIAquick spin PCR Purification Kit (Qiagen, 28104)
- DPNI (ThermoFisher, FD1703)
- Qiagen Spin Miniprep Kit (Qiagen, 27104)
- 2x Gibson Assembly Master Mix (New England Biolabs, E2611L)
- Electrocompetent cells derived from DH5α Cells (ThermoFisher, 18265017).
- Electrocompetent cells derived from One Shot BL21 Star (DE3) Chemically Competent *E. coli* (Invitrogen, C601003)
- LB, Miller (ThermoFisher, BP1426)
- 2% Agarose E-gels, SYBR Safe (Invitrogen, A42135)
- LB agar plates with ampicillin (100 µg/mL)
- MicroPulser Electroporation Cuvettes, 0.2 cm gap (BioRad, 1652086)

Protocol

1) Prepare the insert DNA with PCR as described in Section 1
    a. Use ga4_fwd / pET_ds_rev as these will result in a PCR product that has the 30 base homology domains for Gibson Assembly
    b. We ordered eBlocks with the 30 base homology domains flanking the T7 promoter and terminator domains, but extended T7fwd / T7rev primers containing the homology domains could also be used to prepare inserts for cloning (T7fwd_ga4 / T7rev_pET_ds in Supporting File S3)
2) Prepare the pET backbone with PCR:

| **Component** | **Final volume or concentration** |
|---|---|
| 2x Phusion MM | 37.5 µL |
| ga4_fwd primer | 0.5 µmol/L |
| pET_ds_rev primer | 0.5 µmol/L |
| Ambion water | X µL to 74.85 µL total |

3) Add 0.15 µL of 10 ng/µL pETDuet-1 plasmid DNA into each PCR tube containing master mix, pipette mix

4) Add samples to a thermocycler and execute the following protocol with the lid temperature set to 105 °C

| Temp | Time | Repeat |
|---|---|---|
| 98 °C | 5 min | 1x |
| 98 °C | 30 s | 30x |
| 62 °C | 30 s | 30x |
| 72 °C | 2 min | 30x |
| 72 °C | 5 min | 1x |
| 4 °C | hold | |

5) Run a PCR clean up with a QIAquick Spin PCR Purification Kit
    a. See Step 4 in Section 1
    b. Elute in 30 µL Buffer EB
6) To the 35 µL purified pET backbone DNA, add 1.5 µL DPNI and 3.5 µL DPNI CutSmart Reaction Buffer
7) Incubate at 37 °C for 1 h
8) Run a PCR clean up with a QIAquick Spin PCR Purification Kit
    a. See Step 4 in Section 1
    b. Elute in 30 µL Buffer EB
9) Measure the concentration of the purified PCR products with A260 of undiluted samples on a DeNovix D-11 Series Spectrophotometer
10) Using purified DNA insert and DNA backbone, prepare the following Gibson Assembly mix:

| Component | Final volume or concentration |
|---|---|
| 2x NEB GA mix | 3.75 µL |
| DNA insert | 15 ng/µL (final concentration) |
| DNA backbone | 15 ng/µL (final concentration) |
| Ambion water | X µL to 7.5 µL total |

11) Incubate Gibson Assembly mix at 50 ºC for 1 h
12) Mix 1.5 µL of Gibson Assembly mixture with 50 µL of electrocompetent DH5α cells in MicroPulser Electroporation Cuvettes
13) Load cuvette into Eppendorf Eporator and run electroporation at 2500 V
14) Mix all 50 µL cells with 900 µL of SOC media in 1.7 mL microfuge tubes and grow with 1000 rpm shaking for 1 h time at 37 ºC
15) Spin down the outgrowth cultures at 12800 rpm for 1 min
16) Remove 700 µL of supernatant and discard
17) Resuspend the cell pellet with the remaining media in the 1.7 mL microfuge tube and plate all of this resuspension (≈200 µL) on LB agar plates supplemented with ampicillin
18) Incubate plates overnight at 37 ºC

19) The next day, pick 2 to 5 colonies, mix in 50 µL of water and conduct a colony PCR to verify insert size with gel electrophoresis
    a. Use the following PCR protocol:

| **Component** | **Final volume or concentration** |
|---|---|
| 2x Phusion MM | 27.5 µL |
| pET_seq_fwd primer | 0.5 µmol/L |
| pET_seq_rev primer | 0.5 µmol/L |
| Colony in water | 2 µL |
| Ambion water | X µL to 55 µL total |

    b. Mix 2 µL of PCR product with 18 µL of water and load all 20 µL on a 2 % Agarose E-gel to check insert size
20) For colonies that have the correct insert size, purify the PCR products with a QIAquick Spin PCR Purification Kit
    a. See Step 4 in Section 1
21) Send purified PCR products for Sanger sequencing verification using the primers from the colony PCRs as the sequencing primers
22) For colonies with sequence verified inserts, start overnight cultures in 5 mL of LB+Amp media in 14 mL Falcon tubes
    a. Grow at 37 ºC with 300 rpm shaking for at least 16 hr
23) From the overnight cultures, extract plasmids with a Qiagen Spin Miniprep Kit according to manufacture protocol
24) Transform purified plasmids into electrocompetent BL21 Star (DE3) *E. coli*
    a. Mix 0.5 µL to 2 µL of purified plasmid with 50 µL of electrocompetent cells
        i. Typically adding ≈50 ng total plasmid DNA
        ii. Do not exceed 2 µL of total plasmid DNA volume
    b. Repeat steps 13 through 18
25) Prepare glycerol stocks from transformants and store at -80 ºC for future use
    a. Prepare 3 mL overnight liquid cultures (LB+Amp)
    b. Mix 500 µL of 40 % by volume glycerol with 500 µL of overnight culture

### 5 RNA production, extraction, and protection from *E. coli*

<u>Materials</u>

- Bio-Rad T-100 Thermal cycler (BioRad, 1861096)
- Qubit-4 Fluorometer (ThermoFisher)
- Genesys 30 visible spectrophotometer (ThermoFisher, 840-277000).
- Falcon 14 mL Round Bottom High Clarity PP Test Tube (Corning, 352059)
- Falcon 50 mL High Clarity Conical Centrifuge Tubes (Corning, 352098)
- Falcon 15 mL Conical Centrifuge Tubes (Corning, 352196)
- 0.5 mol/L ethylenediaminetetraacetic acid (EDTA), pH 8.0 (Invitrogen, AM9260G)
- THE RNA Storage Solution (ThermoFisher, AM7001)
- RNAprotect Bacteria Reagent (Qiagen, 1018380)
- RNase Away Decontamination Reagent, 250 mL (ThermoFisher, 10328011)
- Nuclease-Free Water (Ambion, AM9938)
- TE, pH 8.0, RNase-free (ThermoFisher, AM9849)
- Ethanol, 200 PROOF, (Decon Labs, DSP-AZ-1)
- LB media, Miller (ThermoFisher, BP1426)
- Isopropyl ß-D-1-thiogalactopyranoside (IPTG) (Millipore Sigma, I5502)
- DNase I HC (50 U/µL), RNase-free and associated 10x DNA Digestion Buffer (ThermoFisher, EN0523)
- TRI Reagent (ZymoResearch, R2050)
- Chloroform, Molecular Biology Reagent (ThermoFisher, J67241-AP)
- RNA Binding Buffer (ZymoResearch, R1013)
- RNA Wash Buffer (ZymoResearch, R1003)
- Zymo-Spin™ IIICG Column (ZymoResearch, C1006-50-G)
- Collection Tubes (ZymoResearch, C1001-50)
- Qubit RNA HS Buffer Kit (ThermoFisher, Q32852)

<u>Protocol</u>

*RNA production in* E. coli:

1) Start overnight culture in 14 mL Falcon tubes from glycerol stock stab in 3 mL LB+Amp (100 µg/mL) media and grow at 37 ºC and 300 rpm shaking for 16 hours
2) Dilute overnight cultures 1:100 into 10 mL of LB+Amp (100 µg/mL) with 100 µmol/L IPTG and grow in 50 mL Falcon tubes at 37 ºC and 300 rpm shaking until the culture reaches an optical density at 600 nm of 0.5 to 0.8
    a. Optical density measurements were conducted on a Genesys 30 visible spectrophotometer

*RNA extraction from* E. coli:

3) Transfer 1.5 mL of culture to 15 mL Falcon centrifuge tube
4) Add 3 mL of RNA protect Bacteria Reagent incubate 5 min at room temperature
5) Centrifuge for 10 min at 5000 x g at 4 °C

6) Decant the supernatant, remove residual supernatant
7) Resuspend the cells pellet in 100 µL 50 mmol/L EDTA solution
8) Add 1 mL of TRI Reagent
9) Homogenize for 5 min at room temperature
10) Add 200 µL of Chloroform and vortex for 15 s
11) Incubate the homogenate for 5 min at room temperature
12) Centrifuge at 12,000 x g for 10 min at 4°C for phase separation
13) Transfer 600 µL of top aqueous phase to a new 1.5 mL microcentrifuge tube
14) Add 600 µL of 100 % ethanol and mix immediately
15) Transfer 700 µL of the sample to the Zymo-Spin™ IIICG Column in a Zymo Collection Tube
16) Centrifuge for 30 s at 7,000 x g at room temperature, remove the supernatant
17) Transfer the remaining 500 µL of the sample to the same Zymo-Spin™ IIICG Column in a Zymo Collection Tube
18) Centrifuge for 30 s at 7,000 x g
19) Transfer the column into a new collection tube
20) Add 700 µL of RNA Wash Buffer to the column and centrifuge for 30 s at 7,000 x g at room temperature, remove the supernatant
21) Add 400 µL of RNA Wash Buffer to the column and centrifuge for 1.5 min at 12,000 x g
22) Transfer the column carefully into an RNase free tube
23) Add 50 µL DNase/RNase-Free Water directly to the column matrix, incubate for 5 min at room temperature and centrifuge for 2 min at 12,000 x g
24) In an RNase-free PCR tube, add 5 µL DNase I HC (50 U/µL) to 6 µL 10x DNA Digestion Buffer and mix by gentle inversion
25) Add all 11 µL of DNase I mixture to 50 µL of extracted total RNA
26) Incubate at 37°C for 30 min
27) Add 120 µL RNA Binding Buffer supplemented with EDTA (final concentration 36 mmol/L**)**
28) Add 180 µL of 100% ethanol and mix immediately
29) Transfer sample to the Zymo-Spin IC column and centrifuge for 30 s at 7,000 x g
30) Transfer the column into a new Zymo collection tube
31) Add 700 µL RNA Wash Buffer to the column and centrifuge for 30 sat 7,000 x g
32) Discard the supernatant
33) Add 400 µL RNA Wash Buffer to the column and Centrifuge for 1.5 min at 12,000 x g
34) Transfer the column carefully into an RNase free tube
35) Add 20 µL of sterile TE buffer directly to the column matrix, incubate for 5 min at room temperature and centrifuge for 2 min at 12,000 x g
36) Transfer to the fresh tube with 20 µL of THE RNA Storage Solution for prolonged storage
37) Dilute total RNA 1:5 by volume with THE RNA Storage Solution before quantitation with Qubit
38) Verify RNA concentration using Qubit with Qubit RNA HS Buffer Kit
39) The protocol can be paused at this stage and the RNA stored at -80 $^{o}$C

*Total RNA protection with blocking oligonucleotide:*

40) Transfer 240 ng of purified total RNA from *E. coli* into a fresh PCR tube (calculate necessary volume according to Qubit quantitation)
41) Add 2.2 µL of blocking oligonucleotide (stock concentration 1000 µmol/L) and add RNA Storage Solution to reach a total volume of 14 µL
42) Transfer the sample to a thermocycler and run the following annealing protocol:

| **Temp (ramp)** | **Time** | **Repeat** |
|---|---|---|
| 90 °C | 5 min | 1x |
| 90 °C (-0.1 °C) | 6 s | 99x |
| 80 °C (-0.1 °C) | 6 s | 99x |
| 70 °C (-0.1 °C) | 6 s | 99x |
| 60 °C (-0.1 °C) | 6 s | 99x |
| 50 °C (-0.1 °C) | 6 s | 99x |
| 40 °C (-0.1 °C) | 6 s | 99x |
| 30 °C (-0.1 °C) | 6 s | 99x |
| 20 °C | hold | |

43) Verify RNA concentration using Qubit with Qubit RNA HS Buffer Kit
    a. We found that the presence of the blocking oligonucleotide would typically result in an increased concentration from our measurements of the RNA prior to blocking oligonucleotide addition

### 6 RT-qPCR experiments

Materials

- ViiA 7 Real-Time PCR System (Applied Biosystems)
- SuperScript III Platinum SYBR Green One-Step qRT-PCR Kit (ThermoFisher, 11736059)
- Platinum *Taq* DNA Polymerase (ThermoFisher, 10966018)
- Applied Biosystems MicroAmp Optical 96-Well Reaction Plate (ThermoFisher, N8010560)
- Applied Biosystems MicroAmp Optical Adhesive Film (ThermoFisher, 4360954)
- Eppendorf PCR-Cooler 0.2 mL for 96-well PCR plates (Eppendorf, 022510541)
- 8 channel VOYAGER adjustable tip spicing Pipette, 12.5 µL (INTEGRA Biosciences, 4721)
- 96-well PCR-Cooler rack (Eppendorf, 022510541)
- 12.5 µL tips (INTEGRA Biosciences, 4405)
- VIAFLO, single channel electronic Pipette, 300 µL (INTEGRA Biosciences, 4013)
- 300 µL tips (INTEGRA Biosciences, 4435)

Protocol

1) For all samples, normalize purified RNA that is annealed with blocking oligonucleotide to a final concentration of 1 ng/µL with THE RNA Storage Solution
    a. Verify RNA concentrations after dilutions using a Qubit – 1 ng/µL is approximately the lowest concentration that can be reliably measured
2) Using THE RNA Storage Solution, dilute the normalized RNA to a final concentration of 50 pg/µL if from *E. coli* or a 1 pg/µL if from IVT
    a. These dilutions should be done for both gate (G) RNA and uncleaved control (U) RNA, as well as cleaved control RNA (C) if included
3) Prepare aliquots in 1.5 mL microcentrifuge tubes, store in -80°C
    a. Thaw before using, vortex and centrifuge well.
4) Set up RT-qPCR reactions on ice using SuperScript III Platinum SYBR Green One-Step qRT-PCR Kit. Below are the volumes for a single reaction, but these volumes were typically scaled up to make a master mix for multiple samples without the RNA added

| Components (Single reaction) | Sample | NTC | NRTC |
|---|---|---|---|
| SuperScriptR III RT/PlatinumR *Taq* Mix | 0.5 µL | 0.5 µL | 0.5 µL - Platinum *Taq* DNA Polymerase |
| 2X SYBRR Green Reaction Mix | 12.5 µL | 12.5 µL | 12.5 µL |
| Forward primer (10 µmol/L stock) | 0.5 µL | 0.5 µL | 0.5 µL |
| Reverse primer (10 µmol/L stock) | 0.5 µL | 0.5 µL | 0.5 µL |
| ROX Reference Dye | 0.05 µL | 0.05 µL | 0.05 µL |
| Purified and oligo blocked RNA (1 pg/µL or 50 pg/µL) | 2 µL | No template | 2 µL |
| RNase free water to 25 µL | 9 µL | 7 µL | 9 µL |

5) Add 23 µL of master mix to the wells of Applied Biosystems MicroAmp Optical 96-Well Reaction Plate using 300 µL single channel electronic for repeated dispensing
6) Add 2 µL of template from PCR strip tubes to the plate well using 12.5 µL 8 channel adjustable tip electronic pipette for repeated transfer and pipette mixing
7) Seal the plate with Applied Biosystems MicroAmp Optical Adhesive Film
8) Centrifuge the 96-well plate in PCR-Cooler rack at 1000g for 1 min at 4 °C
    a. Keep plate in the PCR-Cooler rack until ready to load the plate into the instrument
9) **Supplementary Information Section 2.1 has a diagram of the plate layout we typically used for these experiments**
    a. This section also describes how we prepared the dilution series for primer efficiency analysis using the normalized RNA samples
10) Set up ViiA™ 7 Real-Time PCR System experiment program:
    a. Select the 96-Well Block type (0.2 mL) using to run the experiment
    b. Select the Comparative ΔΔCt experiment type
    c. Select the SYBR® Green reagent using to detect the target sequence
    d. Select the Standard ramp speed for the experiment
    e. Melting curve analysis is run by default for the SYBR Green reagent
    f. Use the following cycling protocol:

| Temp | Time | Repeat |
|---|---|---|
| 37 °C or 50 °C | 10 min | 1x |
| 95 °C | 5 min | 1x |
| 95 °C | 15 s | 40x |
| 60 °C | 1 min | 40x |
| 60 °C to 95 °C | | 1x |
