## Supplementary material for "Prevention of ribozyme catalysis through cDNA synthesis enables accurate RT-qPCR measurements of context-dependent ribozyme activity": Uncropped gel images, raw data, and analysis code: RT_qPCR_Rz_analysis_ave3.py

```
import numpy as np
import matplotlib.pyplot as plt
from matplotlib.ticker import MaxNLocator
import openpyxl as opxl
import pandas as pd
import warnings
from sklearn.metrics import r2_score
from sklearn.linear_model import LinearRegression


def RT_qPCR_Rz_analysis_ave3(file_path,n_samples,primers,no_PCR_cycles,G_pos,\
                        C,C_pos,U_pos,non_std_crv,shift_ave,tech_rep_ave,ref_well,ref_well_u,ctrls,\
                        rn_type,std_crv_x,n_primer_sets,fity_o,fity_u,ignore_y_int,is_std_crv):
 
    # generating the samples to pull in from above data
    plate_positions = []
    con_ref_wells = ref_well + ref_well_u
    frst = int(ref_well[0][1:]) # pulling out the column to start at
    for r in range(len(con_ref_wells)):
        for c in range(n_samples+ctrls):
            plate_positions.append(con_ref_wells[r][0]+str(c+frst))
    
    #print(plate_positions)
    
    well_hash = {}
    wrow = ['A','B','C','D','E','F','G','H']
    wcol = [i+1 for i in range(12)]
    for n in range(len(wrow)):
        for m in range(len(wcol)):
            well_hash[wrow[n]+str(wcol[m])] = m+12*n+1
            
    well_numbers = [well_hash[plate_positions[n]] for n in range(len(plate_positions))]
    
    '''
    Amplification data
    '''
    # reading in data
    rn_data = pd.read_excel(file_path,sheet_name='Amplification Data',skiprows=35)
    results_data = pd.read_excel(file_path,sheet_name='Results',skiprows=35)
    
    results_df = pd.DataFrame(data=results_data,columns=['Well','Well Position','CT'])
    results_df = results_df[results_df['Well Position'].isin(plate_positions)]
    
    Rn_df = pd.DataFrame(data=rn_data,columns=['Well','Cycle',rn_type])
    Rn1 = Rn_df.loc[Rn_df['Well'] == 1]
    Rn_cycles = pd.DataFrame(data=Rn1,columns=['Cycle'])
    
    '''
    Melt curves data
    '''
    Rn_array = []
    Rn_full_curves = []
    
    # reading in data
    mcp_data = pd.read_excel(file_path,sheet_name='Multicomponent Data',skiprows=35)
   
    for m in range(len(well_numbers)):
        Rn = Rn_df.loc[Rn_df['Well'] == well_numbers[m]]
        Rn_array.append(pd.DataFrame(data=Rn,columns=[rn_type]))
        
    # Rn_full_curves = []
    # for m in range(len(plate_positions)):
    #     Rn_full_curves.append(mcp_data[mcp_data['Well Position'] == plate_positions[m]]['SYBR']/mcp_data[mcp_data['Well Position'] == plate_positions[m]]['ROX'])
        
    for m in range(len(plate_positions)):
        Rn_full_curves.append(np.diff(-mcp_data[mcp_data['Well Position'] == plate_positions[m]]['SYBR']))
    
    mc_xdata = (np.array([n for n in range(257)])-2*no_PCR_cycles)/5.11+60
    
    '''
    Ct data
    '''
    ct_vec = results_df['CT'].values
    
    # creating a ct matrix
    ct_mat = np.zeros([n_primer_sets,n_samples])
    i = 0
    cnt = 0
    for k in range(primers):
        for n in range(len(ref_well)):
            for m in range(n_samples):
                ct_mat[cnt,m] = ct_vec[i+m]
            i += n_samples + ctrls
            cnt += 1
    
    # average of ct values for each individual sample
    ct_ave_PCRo = np.mean(ct_mat[0:len(ref_well),:],0)
    ct_ave_PCRu = np.mean(ct_mat[len(ref_well_u):,:],0)
    # std of ct values for each individual sample
    ct_std_PCRo = np.std(ct_mat[0:len(ref_well),:],0)
    ct_std_PCRu = np.std(ct_mat[len(ref_well_u):,:],0)

   
    if tech_rep_ave == 1: 
        
        '''
        Curves shifted relative to PCRo
        '''
        # placeholders for variables not used in this analysis
        ct_shift = 0
        delta_ct_unc = 0
        delta_ct_gate = 0
        delta_ct_gate_crt = 0
        ddCt_Gu_i =  np.zeros(len(ref_well))
        fit_frac_ddCt_Gu_i = []
        
        ## A possible normalization factor for the comparing the two PCRs
        # ct_shift_nf = np.zeros(len(ref_well))
        # for k in range(len):
        #     ct_shift_nf[k] = results_df[results_df['Well Position'] == ref_well_u[k]]['CT'].values/results_df[results_df['Well Position'] == ref_well[k]]['CT'].values
        ct_shift_nf = np.ones(len(ref_well)) # no normalization correction
        
        # Computing the average difference in PCRo Ct between U and G
        
        ct_shift_ave = np.zeros([n_samples]) # ct shift matrix
        ct_shift_std = np.zeros([n_samples]) # ct shift matrix
        for n in range(n_samples):
            ct_shift_ave[n] = (ct_ave_PCRo[n] - ct_ave_PCRo[U_pos-1])
            ct_shift_std[n] = np.sqrt(ct_std_PCRo[n]**2 + ct_std_PCRo[U_pos-1]**2)
            
        # delta ct for the standard curve
        delta_ct_unc_ave_o = np.zeros([n_samples-non_std_crv])
        delta_ct_unc_ave_u = np.zeros([n_samples-non_std_crv])
        delta_ct_unc_std_o = np.zeros([n_samples-non_std_crv])
        delta_ct_unc_std_u = np.zeros([n_samples-non_std_crv])
        i = 0
        for m in range(n_samples-non_std_crv):
            delta_ct_unc_ave_o[m] = (ct_ave_PCRo[m+1]) - (ct_ave_PCRo[U_pos-1])
            delta_ct_unc_ave_u[m] = (ct_ave_PCRu[m+1]) - (ct_ave_PCRu[U_pos-1])
            delta_ct_unc_std_o[m] = np.sqrt((ct_std_PCRo[m+1])**2 + (ct_std_PCRo[U_pos-1])**2)
            delta_ct_unc_std_u[m] = np.sqrt((ct_std_PCRu[m+1])**2 + (ct_std_PCRu[U_pos-1])**2)
        
        # delta ct for gate
        delta_ct_gate_ave_u = (ct_ave_PCRu[G_pos-1]) - (ct_ave_PCRu[U_pos-1])
        delta_ct_gate_ave_o = (ct_ave_PCRo[G_pos-1]) - (ct_ave_PCRo[U_pos-1])
        delta_ct_gate_std_u = np.sqrt((ct_std_PCRu[G_pos-1])**2 + (ct_std_PCRu[U_pos-1])**2)
        delta_ct_gate_std_o = np.sqrt((ct_std_PCRo[G_pos-1])**2 + (ct_std_PCRo[U_pos-1])**2)
        
        
        # # shifted delta ct for gate
        # delta_ct_gate_crt_ave_o = (ct_ave_PCRo[G_pos-1]-ct_shift_ave[G_pos-1]) - (ct_ave_PCRo[U_pos-1]-ct_shift_ave[U_pos-1])
        # delta_ct_gate_crt_ave_u = (ct_ave_PCRu[G_pos-1]-ct_shift_ave[G_pos-1]) - (ct_ave_PCRu[U_pos-1]-ct_shift_ave[U_pos-1])
        # delta_ct_gate_crt_std_o = np.sqrt((ct_std_PCRo[G_pos-1]-ct_std_PCRo[U_pos-1])**2 + ct_shift_std[G_pos-1]**2)
        # delta_ct_gate_crt_std_u = np.sqrt((ct_std_PCRu[G_pos-1]-ct_std_PCRu[U_pos-1])**2 + ct_shift_std[G_pos-1]**2)
        
        delta_ct_gate_crt_ave_o = delta_ct_gate_ave_o - ct_shift_ave[G_pos-1]
        delta_ct_gate_crt_ave_u = delta_ct_gate_ave_u - ct_shift_ave[G_pos-1]
        
        delta_ct_gate_crt_std_o = np.sqrt(delta_ct_gate_std_o**2 + ct_shift_std[G_pos-1]**2)
        delta_ct_gate_crt_std_u = np.sqrt(delta_ct_gate_std_u**2 + ct_shift_std[G_pos-1]**2)
        
        '''
        Fitting lines to cleavage standard curve 
        '''
        
        xdata = np.array(std_crv_x)
        
        if is_std_crv == 1: # only do fitting if there is a standard curve
            
            if std_crv_x[0] == 1: 
                # Including 0 point in fitting
                xdata_fit = xdata
                ydata_o = np.insert(delta_ct_unc_ave_o,0,0)
                ydata_u = np.insert(delta_ct_unc_ave_u,0,0)
            else:
                std_crv_x_fit = [1]+std_crv_x
                xdata_fit = np.array(std_crv_x_fit)
                ydata_o = delta_ct_unc_ave_o
                ydata_u = delta_ct_unc_ave_u
            
            # fitting line to standard curve for PCRo
            regr = LinearRegression(fit_intercept=fity_o).fit(np.log2(xdata.reshape(1,-1).T),ydata_o.reshape(1,-1).T)
            PCRo_fit_line = regr.predict(np.log2(xdata_fit.reshape(1,-1).T))
            
            r2_o = regr.score(np.log2(xdata.reshape(1,-1).T),ydata_o.reshape(1,-1).T)
            
            fit_line_par_o = np.append(regr.coef_[0],regr.intercept_)
            fit_line_par_o = np.append(fit_line_par_o,r2_o)
            
            # fitting line to standard curve for PCRu
            regr = LinearRegression(fit_intercept=fity_u).fit(np.log2(xdata.reshape(1,-1).T),ydata_u.reshape(1,-1).T)
            PCRu_fit_line = regr.predict(np.log2(xdata_fit.reshape(1,-1).T))
            
            r2_u = regr.score(np.log2(xdata.reshape(1,-1).T),ydata_u.reshape(1,-1).T)
            
            fit_line_par_u = np.append(regr.coef_[0],regr.intercept_)
            fit_line_par_u = np.append(fit_line_par_u,r2_u)
            
            ideal_frac_clv_gate_crt = []
            
            #for n in range(len(ref_well)):
            ideal_frac_clv_gate_crt.append(2**-delta_ct_gate_crt_ave_u)
            mean_ideal_clv_gate_crt = np.mean(ideal_frac_clv_gate_crt)
            
            # technically divided by -1 for slope
            std_ideal_clv_gate_crt = (mean_ideal_clv_gate_crt)*np.log(2)*delta_ct_gate_crt_std_u
            
            fit_frac_clv_gate_crt = []
            
            if ignore_y_int == 1:
                a = fit_line_par_u[0]
                b = 0
            else:
                a = fit_line_par_u[0]
                b = fit_line_par_u[1]
                
            fit_frac_clv_gate_crt.append(2**((delta_ct_gate_crt_ave_u-b)/a))
            mean_fit_clv_gate_crt = np.mean(fit_frac_clv_gate_crt)
            
            std_fit_clv_gate_crt = -(mean_fit_clv_gate_crt)*1/a*np.log(2)*delta_ct_gate_crt_std_u
        
        else:
            
            PCRo_fit_line = 0
            fit_line_par_o = 0
            PCRu_fit_line = 0
            fit_line_par_u = 0
            
            # Still compute the ideal fraction cleaved
            ideal_frac_clv_gate_crt = []
            ideal_frac_clv_gate_crt.append(2**-delta_ct_gate_crt_ave_u)
            mean_ideal_clv_gate_crt = np.mean(ideal_frac_clv_gate_crt)
            
            # technically divided by -1 for slope
            std_ideal_clv_gate_crt = (mean_ideal_clv_gate_crt)*np.log(2)*delta_ct_gate_crt_std_u
            
            fit_frac_clv_gate_crt = 0
            mean_fit_clv_gate_crt = 0
            std_fit_clv_gate_crt = 0
    
    
    else:  
        
        '''
        Curves shifted relative to PCRo
        '''
        
        ## A possible normalization factor for the comparing the two PCRs
        # ct_shift_nf = np.zeros(len(ref_well))
        # for k in range(len):
        #     ct_shift_nf[k] = results_df[results_df['Well Position'] == ref_well_u[k]]['CT'].values/results_df[results_df['Well Position'] == ref_well[k]]['CT'].values
        ct_shift_nf = np.ones(len(ref_well)) # no normalization correction
        
        l = np.zeros(len(ref_well),dtype=int) # locations of ref_wells
        for k in range(len(ref_well)): 
            l[k] = plate_positions.index(ref_well[k])-int(ref_well[k][1:])+1
            
        lu = np.zeros(len(ref_well),dtype=int) # locations of ref_wells
        for k in range(len(ref_well_u)): 
            lu[k] = plate_positions.index(ref_well_u[k])-int(ref_well_u[k][1:])+1
            
        l_refs = np.concatenate([l,lu])
        
        ct_shift = np.zeros([len(ref_well),n_samples]) # ct shift matrix
        for k in range(len(ref_well)): 
            for n in range(n_samples):
                ct_shift[k,n] = (results_df[results_df['Well Position'] == plate_positions[l[k]+n]]['CT'].values - results_df[results_df['Well Position'] == ref_well[k]]['CT'].values)*ct_shift_nf[k]
        
        # averaging the shift from the PCRo replicates
        # - This is equivalent to averaging U and G replicates and then subtracting 
        ct_shift_ave = np.mean(ct_shift,0)
        ct_shift_std = np.std(ct_shift,0)
        
        # delta ct for the standard curve
        delta_ct_unc = np.zeros([n_primer_sets,n_samples-non_std_crv])
        i = 0
        cnt = 0
        for k in range(primers):
            for n in range(len(ref_well)):
                for m in range(n_samples-non_std_crv):
                    delta_ct_unc[cnt,m] = (ct_vec[i+m+1]) - (ct_vec[i])
                i += n_samples + ctrls
                cnt += 1
                
        delta_ct_unc_ave_o = np.mean(delta_ct_unc[0:len(ref_well),:],0)
        delta_ct_unc_ave_u = np.mean(delta_ct_unc[len(ref_well):,:],0)
        delta_ct_unc_std_o = np.std(delta_ct_unc[0:len(ref_well),:],0)
        delta_ct_unc_std_u = np.std(delta_ct_unc[len(ref_well):,:],0)
        
        # delta ct for gate
        delta_ct_gate = np.zeros(n_primer_sets)
        i = 0
        cnt = 0
        for k in range(primers):
            for n in range(len(ref_well)):
                delta_ct_gate[cnt] = (ct_vec[i+G_pos-1]) - (ct_vec[i])
                i += n_samples + ctrls
                cnt += 1
        
        delta_ct_gate_ave_o = np.mean(delta_ct_gate[0:len(ref_well)],0)
        delta_ct_gate_ave_u = np.mean(delta_ct_gate[len(ref_well):],0)
        delta_ct_gate_std_o = np.std(delta_ct_gate[0:len(ref_well)],0)
        delta_ct_gate_std_u = np.std(delta_ct_gate[len(ref_well):],0)
        
        # individual dCt_Gu values shifted by ave dCt_Go
        ddCt_Gu_i =  np.zeros(len(ref_well))
        for n in range(len(ref_well)):
            ddCt_Gu_i[n] = delta_ct_gate[len(ref_well)+n] - ct_shift_ave[G_pos-1]
        
        # # shifted delta ct for gate
        delta_ct_gate_crt = np.zeros(n_primer_sets)
        # i = 0
        # cnt = 0
        # for k in range(primers):
        #     for n in range(len(ref_well)): 
        #         if shift_ave == 1:
        #             delta_ct_gate_crt[cnt] = (ct_vec[i+G_pos-1]-ct_shift_ave[G_pos-1]) - (ct_vec[i]-ct_shift_ave[0])
        #         else:
        #             delta_ct_gate_crt[cnt] = (ct_vec[i+G_pos-1]-ct_shift[n,G_pos-1]) - (ct_vec[i]-ct_shift[n,0])
        #         i += n_samples + ctrls
        #         cnt += 1
                
        # delta_ct_gate_crt_ave_o = np.mean(delta_ct_gate_crt[0:len(ref_well)],0)
        # delta_ct_gate_crt_ave_u = np.mean(delta_ct_gate_crt[len(ref_well):],0)
        # delta_ct_gate_crt_std_o = np.std(delta_ct_gate_crt[0:len(ref_well)],0)
        # delta_ct_gate_crt_std_u = np.std(delta_ct_gate_crt[len(ref_well):],0)
        
        delta_ct_gate_crt_ave_o = delta_ct_gate_ave_o - ct_shift_ave[G_pos-1]
        delta_ct_gate_crt_ave_u = delta_ct_gate_ave_u - ct_shift_ave[G_pos-1]
        
        delta_ct_gate_crt_std_o = np.sqrt(delta_ct_gate_std_o**2 + ct_shift_std[G_pos-1]**2)
        delta_ct_gate_crt_std_u = np.sqrt(delta_ct_gate_std_u**2 + ct_shift_std[G_pos-1]**2)
        
        '''
        Fitting lines to cleavage standard curve 
        '''
        
        xdata = np.array(std_crv_x)
        std_crv_x_fit = [1]+std_crv_x
        xdata_fit = np.array(std_crv_x_fit)
        
        # fitting line to standard curve for PCRo
        ydata_o = delta_ct_unc_ave_o
        p_val_o = np.polyfit(np.log2(xdata),ydata_o,1)
        PCRo_fit_line = p_val_o[1] + p_val_o[0]*np.log2(xdata_fit)
        r2_o = r2_score(ydata_o,PCRo_fit_line[1:])
        
        fit_line_par_o = np.append(p_val_o,r2_o)
        
        # fitting line to standard curve for PCRu
        ydata_u = delta_ct_unc_ave_u
        p_val_u = np.polyfit(np.log2(xdata),ydata_u,1)
        PCRu_fit_line = p_val_u[1] + p_val_u[0]*np.log2(xdata_fit)
        r2_u = r2_score(ydata_u,PCRu_fit_line[1:])
        
        fit_line_par_u = np.append(p_val_u,r2_u)
        
        fit_frac_clv_gate = []
        
        #for n in range(len(ref_well)):
        ideal_frac_clv_gate_crt = []
        
        #for n in range(len(ref_well)):
        ideal_frac_clv_gate_crt.append(2**-delta_ct_gate_crt_ave_u)
        mean_ideal_clv_gate_crt = np.mean(ideal_frac_clv_gate_crt)
        
        std_ideal_clv_gate_crt = -(mean_ideal_clv_gate_crt)*np.log(2)*delta_ct_gate_crt_std_u
        
        
        fit_frac_clv_gate_crt = []
        fit_frac_ddCt_Gu_i = []
        
        #for n in range(len(ref_well)):
        fit_frac_clv_gate_crt.append(2**((delta_ct_gate_crt_ave_u-p_val_u[1])/p_val_u[0]))
        
        for n in range(len(ref_well)):
            fit_frac_ddCt_Gu_i.append(2**((ddCt_Gu_i[n]-p_val_u[1])/p_val_u[0]))
    
        mean_fit_clv_gate_crt = np.mean(fit_frac_clv_gate_crt)
        
        std_fit_clv_gate_crt = np.std(fit_frac_clv_gate)
        std_fit_clv_gate_crt = -(mean_fit_clv_gate_crt)*1/p_val_u[0]*np.log(2)*delta_ct_gate_crt_std_u
        
        # ideal_ct = np.log2(np.ones(len(std_crv_x))/np.array(std_crv_x))
        
        # x_list = len(ref_well)*std_crv_x
        # xdata = np.array(x_list)
        
        # # fitting line to standard curve for PCRo
        # ydata_o = delta_ct_unc[0:len(ref_well),:].flatten()
        # p_val_o = np.polyfit(np.log2(xdata),ydata_o,1)
        # PCRo_fit_line = p_val_o[1] + p_val_o[0]*np.log2(xdata[0:len(std_crv_x)])
        
        # # fitting line to standard curve for PCRu
        # ydata_u = delta_ct_unc[len(ref_well):,:].flatten()
        # p_val_u = np.polyfit(np.log2(xdata),ydata_u,1)
        # PCRu_fit_line = p_val_u[1] + p_val_u[0]*np.log2(xdata[0:len(std_crv_x)])
        
        # ideal_frac_clv_gate = []
        # fit_frac_clv_gate = []
        
        # for n in range(len(ref_well)):
        #     ideal_frac_clv_gate.append(2**(-delta_ct_gate[n+len(ref_well)]))
        #     fit_frac_clv_gate.append(2**((delta_ct_gate[n+len(ref_well)]-p_val_u[1])/p_val_u[0]))
        
        # mean_ideal_clv_gate = np.mean(ideal_frac_clv_gate)
        # mean_fit_clv_gate = np.mean(fit_frac_clv_gate)
        
        # std_ideal_clv_gate = np.std(ideal_frac_clv_gate)
        # std_fit_clv_gate = np.std(fit_frac_clv_gate)
        
        # ideal_frac_clv_gate_crt = []
        # fit_frac_clv_gate_crt = []
        
        # for n in range(len(ref_well)):
        #     ideal_frac_clv_gate_crt.append(2**(-delta_ct_gate_crt[n+len(ref_well)]))
        #     fit_frac_clv_gate_crt.append(2**((delta_ct_gate_crt[n+len(ref_well)]-p_val_u[1])/p_val_u[0]))
        
        # mean_ideal_clv_gate_crt = np.mean(ideal_frac_clv_gate_crt)
        # mean_fit_clv_gate_crt = np.mean(fit_frac_clv_gate_crt)
        
        # std_ideal_clv_gate_crt = np.std(ideal_frac_clv_gate_crt)
        # std_fit_clv_gate_crt = np.std(fit_frac_clv_gate_crt)
         
    
    
    return  xdata,Rn_full_curves,Rn_array,Rn_cycles,plate_positions,mc_xdata,\
            ct_shift,ct_shift_ave,ct_shift_std,ddCt_Gu_i,fit_frac_ddCt_Gu_i,\
            delta_ct_unc,delta_ct_gate,delta_ct_gate_crt,delta_ct_gate_ave_u,delta_ct_gate_ave_o,delta_ct_gate_std_u,delta_ct_gate_std_o,\
            PCRu_fit_line,PCRo_fit_line,fit_line_par_u,fit_line_par_o,\
            ideal_frac_clv_gate_crt,fit_frac_clv_gate_crt,\
            delta_ct_unc_ave_o,delta_ct_unc_ave_u,delta_ct_unc_std_o,delta_ct_unc_std_u,\
            delta_ct_gate_crt_ave_o,delta_ct_gate_crt_ave_u,delta_ct_gate_crt_std_o,delta_ct_gate_crt_std_u,\
            mean_ideal_clv_gate_crt,std_ideal_clv_gate_crt,\
            mean_fit_clv_gate_crt,std_fit_clv_gate_crt
```
