## Supplementary material for "Prevention of ribozyme catalysis through cDNA synthesis enables accurate RT-qPCR measurements of context-dependent ribozyme activity": Uncropped gel images, raw data, and analysis code: Supp_Figure_S7.py

```
import numpy as np
import matplotlib.pyplot as plt
from matplotlib.ticker import MaxNLocator
import openpyxl as opxl
import pandas as pd
import warnings
import copy
warnings.filterwarnings("ignore", \
message="Adding an axes using the same arguments as a previous axes currently \
reuses the earlier instance.  In a future version, a new instance will always be \
created and returned.  Meanwhile, this warning can be suppressed, and the future \
behavior ensured, by passing a unique label to each axes instance.")

import sys
sys.path.insert(1,'C:\\Users\\sws5\\OneDrive - National Institute of Standards and Technology (NIST)\\Presentations and write ups\\Papers\\ctRSD RT qPCR\\Analysis code')
from RT_qPCR_Rz_analysis_ave3 import RT_qPCR_Rz_analysis_ave3

'''
##############################################################################
USER DEFINED FILE LOCATION AND PLATE POSITIONS FOR IMPORTING
##############################################################################
'''
results_path = 'C:\\Users\\sws5\\OneDrive - National Institute of Standards and Technology (NIST)\\Presentations and write ups\\Papers\\ctRSD RT qPCR\\Data for upload\\'
exp_path = 'C:\\Users\\sws5\\OneDrive - National Institute of Standards and Technology (NIST)\\Presentations and write ups\\Papers\\ctRSD RT qPCR\\Data for upload\\list_experiments_paths_final.xlsx'

exp_files = pd.read_excel(exp_path,sheet_name='Supp Fig 7')

show_ideal = 1
show_ideal_crv = 0
show_amp = [0,0,\
            0,0] 

Rn_array_list = []
Rn_cycles_list = []
ct_shift_ave_list = []
Rn_full_curves_list = []
mc_xdata_list = []

mean_fit_clv_gate_list = []
std_fit_clv_gate_list = []
mean_ddCt_clv_gate_list = []
std_ddCt_clv_gate_list = []
ddCt_Gu_i_list = []

mean_dCt_Uo_list = []
mean_dCt_Uu_list = []
std_dCt_Uo_list = []
std_dCt_Uu_list = []

dCt_Uu_fit_line = []
dCt_Uo_fit_line = []

fit_line_params_u = []
fit_line_params_o = []

mean_ideal_clv_gate_list = []
std_ideal_clv_gate_list = []

for f in range(len(exp_files)):
    
    folder_path = exp_files['folder_path'][f]+'\\'
    file_name = exp_files['file_name'][f]
    file_path = results_path+folder_path+file_name
    
    n_samples = exp_files['n_samples'][f] 
    primers = exp_files['primers'][f] 
    G_pos = exp_files['G_pos'][f]  # column of plate G is in
    C_pos = exp_files['C_pos'][f]  # column of plate C is in 
    U_pos = 1 # column of plate U is in
    C = exp_files['C'][f]  # cleaved gate present? 1 or 0
    is_std_crv = exp_files['is_std_crv'][f]  # is there a standard curve in the exp?
    
    no_PCR_cycles = 40
    non_std_crv = 2+C # samples other than standard curve (U,C,G for example)
    shift_ave = 1 # should PCRo be averaged for shifting? 1 for technical replicates
    tech_rep_ave = 1 # should the standard curve replicates be shifted 
    ctrls = 0 # number of ctrl wells at the end of the plate
    
    # Fraction uncleaved gate in standard curve
    std_crv_x = [0.75,0.5,0.25,0.1,0.05,0.01]
    # Fit y-int?
    fity_o = 1
    fity_u = 1
    # ignore fit y-int in fraction cleaved calculation
    ignore_y_int = 0
    
    # the reference sample (100% uncleaved with PCRo primers)
    ref_well = exp_files['ref_well'][f].split()
    # the reference sample (100% uncleaved with PCRu primers)
    ref_well_u = exp_files['ref_well_u'][f].split()
    
    rn_type = 'Delta Rn' # which data type to import
    
    # primer types and number of different conditions or replicates
    n_primer_sets = primers*len(ref_well)
    
    
    [xdata,Rn_full_curves,Rn_array,Rn_cycles,plate_positions,mc_xdata,\
            ct_shift,ct_shift_ave,ct_shift_std,ddCt_Gu_i,fit_frac_ddCt_Gu_i,\
            delta_ct_unc,delta_ct_gate,delta_ct_gate_crt,delta_ct_gate_ave_u,delta_ct_gate_ave_o,delta_ct_gate_std_u,delta_ct_gate_std_o,\
            PCRu_fit_line,PCRo_fit_line,fit_line_par_u,fit_line_par_o,\
            ideal_frac_clv_gate_crt,fit_frac_clv_gate_crt,\
            delta_ct_unc_ave_o,delta_ct_unc_ave_u,delta_ct_unc_std_o,delta_ct_unc_std_u,\
            delta_ct_gate_crt_ave_o,delta_ct_gate_crt_ave_u,delta_ct_gate_crt_std_o,delta_ct_gate_crt_std_u,\
            mean_ideal_clv_gate_crt,std_ideal_clv_gate_crt,\
            mean_fit_clv_gate_crt,std_fit_clv_gate_crt] = \
      RT_qPCR_Rz_analysis_ave3(file_path,n_samples,primers,no_PCR_cycles,G_pos,\
                              C,C_pos,U_pos,non_std_crv,shift_ave,tech_rep_ave,ref_well,ref_well_u,ctrls,\
                              rn_type,std_crv_x,n_primer_sets,fity_o,fity_u,ignore_y_int,is_std_crv)
     
    # saving data for plotting each time through
    Rn_array_list.append(Rn_array)
    Rn_cycles_list.append(Rn_cycles)
    ct_shift_ave_list.append(ct_shift_ave)
    Rn_full_curves_list.append(Rn_full_curves)
    mc_xdata_list.append(mc_xdata_list)
    
    mean_fit_clv_gate_list.append(mean_fit_clv_gate_crt)
    std_fit_clv_gate_list.append(std_fit_clv_gate_crt)
    
    mean_ddCt_clv_gate_list.append(delta_ct_gate_crt_ave_u)
    std_ddCt_clv_gate_list.append(delta_ct_gate_crt_std_u)
    
    mean_dCt_Uu_list.append(delta_ct_unc_ave_u)
    mean_dCt_Uo_list.append(delta_ct_unc_ave_o)
    std_dCt_Uu_list.append(delta_ct_unc_std_u)
    std_dCt_Uo_list.append(delta_ct_unc_std_o)
    
    dCt_Uu_fit_line.append(PCRu_fit_line)
    dCt_Uo_fit_line.append(PCRo_fit_line)
    
    fit_line_params_u.append(fit_line_par_u)
    fit_line_params_o.append(fit_line_par_o)
    
    ddCt_Gu_i_list.append(fit_frac_ddCt_Gu_i)
    
    mean_ideal_clv_gate_list.append(mean_ideal_clv_gate_crt)
    std_ideal_clv_gate_list.append(std_ideal_clv_gate_crt)

exp_files['mean_fit_cleave_gate'] = mean_fit_clv_gate_list
exp_files['std_fit_cleave_gate'] = std_fit_clv_gate_list
exp_files['mean_ddCt_clv_gate'] = mean_ddCt_clv_gate_list
exp_files['std_ddCt_clv_gate'] = std_ddCt_clv_gate_list
exp_files['mean_dCt_Uu'] = ddCt_Gu_i_list

#exp_files['mean_dCt_Uu'] = mean_dCt_Uu_list

exp_files['fit_line_par_u'] = fit_line_params_u
exp_files['fit_line_par_o'] = fit_line_params_o

exp_files['fit ddCt_Gu_i'] = ddCt_Gu_i_list

exp_files['mean_ideal_cleave_gate'] = mean_ideal_clv_gate_list
exp_files['std_ideal_cleave_gate'] = std_ideal_clv_gate_list


if show_ideal == 1:
    fs = 13
    
    bar_dfI = exp_files.pivot(index='Gate',columns='bio_rep',values='mean_fit_cleave_gate')
    yerr_dfI = exp_files.pivot(index='Gate',columns='bio_rep',values='std_fit_cleave_gate')
    
    bar_dfI.plot(kind='bar',yerr=yerr_dfI,capsize=4,rot=0,color={1: [0.4,0.4,0.4], 2: [0.6,0.6,0.6], 3: [0.8,0.8,0.8]})
    #scatter_df.plot(kind='scatter')
    plt.ylabel('Fraction uncleaved',fontsize=fs)
    plt.xlabel('Gate',fontsize=fs)
    plt.legend(['wo C','w C'],frameon=False,fontsize=12)
    plt.xticks(fontsize=fs)
    plt.ylim([0,1.1])
    plt.yticks([1,0.75,0.5,0.25,0.1,0],[1,0.75,0.5,0.25,0.1,0],fontsize=fs)
    ax1 = plt.gca()
    ax1.xaxis.set_tick_params(which='both',size=3,width=1,direction='in',top='on')
    ax1.yaxis.set_tick_params(which='both',size=3,width=1,direction='in',right='on')
    
    plt.plot([-1,7],[0.5,0.5],'r',linestyle='--')
    plt.plot([-1,7],[0.1,0.1],color=[0,0.25,1],linestyle='--')
    
    ax2 = ax1.twinx()
    plt.yticks([1,0.75,0.5,0.25,0.1],[0,0.5,1,2,3.32],fontsize=fs)
    plt.ylim([0,1.1])
    #ax2.invert_yaxis()
    ax2.yaxis.set_tick_params(which='both',size=3,width=1,direction='in',right='on')
    plt.ylabel('$\Delta$$\Delta$Ct',fontsize=fs,rotation=270)
        
'''
########################################
Plotting standard curves
########################################
'''
    
plt.figure(4)
ymax_stc = 7.5

p0 = np.zeros(len(exp_files),dtype=int)
p1 = np.zeros(len(exp_files),dtype=int)
smple = int(len(exp_files)/2)

for m in range(smple):
    p0[m] = m+1
    p0[m+smple] = m+7
    
    p1[m] = m+13
    p1[m+smple] = m+13

if std_crv_x[0] == 1:
    std_crv_x_fit = copy.deepcopy(std_crv_x)
    std_crv_x = std_crv_x[1:]
    plot0 = 1
else:
    std_crv_x_fit = [1]+std_crv_x
    plot0 = 0

ms = 4 # marker size
cs = 2 # cap size
fs = 11

'''
Corrected cleavage lines (gate only)
'''

for n in range(len(exp_files)):
    
    plt.subplot(3,6,p0[n])
    if show_ideal_crv == 1:
        plt.plot(-np.log2(std_crv_x_fit),-np.log2(std_crv_x_fit),color = [0.75,0.75,0.75],linestyle = ':',linewidth=1)
        plt.plot(mean_ddCt_clv_gate_list[n],mean_ddCt_clv_gate_list[n],'o',color=[0,0.5,1],alpha=0.25,markersize=ms)
    
    plt.plot(-np.log2(std_crv_x_fit),dCt_Uu_fit_line[n],color = [0.5,0.5,0.5],linestyle = '--',linewidth=1)
    plt.plot(-np.log2(std_crv_x),mean_dCt_Uu_list[n],'o',color='k',markersize=ms)
    plt.errorbar(-np.log2(std_crv_x),mean_dCt_Uu_list[n],yerr=std_dCt_Uu_list[n],capsize=cs,color=[1,0,1],fmt='none')
    
    plt.plot(-np.log2(mean_fit_clv_gate_list[n]),mean_ddCt_clv_gate_list[n],'o',color=[0,0.5,1],markersize=ms)
    plt.errorbar(-np.log2(mean_fit_clv_gate_list[n]),mean_ddCt_clv_gate_list[n],yerr=std_ddCt_clv_gate_list[n],capsize=cs,color=[1,0,1],fmt='none')
    plt.ylim(-1,ymax_stc)
    
    if plot0 == 1:
        plt.plot([0],[0],'o',color='k',markersize=ms)
    
    plt.xticks(fontsize=fs)
    plt.yticks(fontsize=fs)
    #plt.xlim(0,ymax)
    ax1 = plt.gca()
    ax1.xaxis.set_tick_params(which='both',size=3,width=1,direction='in',top='on')
    #ax1.xaxis.set_major_locator(MaxNLocator(integer=True))
    ax1.yaxis.set_tick_params(which='both',size=3,width=1,direction='in',right='on')
    plt.xlabel('-log2(Fraction uncleaved)',fontsize=fs)
    plt.ylabel('$\Delta$Ct_Uu',fontsize=fs)   
    r2 = exp_files['fit_line_par_u'][n][-1]
    slp = exp_files['fit_line_par_u'][n][0]
    y0 = exp_files['fit_line_par_u'][n][1]
    plt.annotate('R^2 = {:.3f}'.format(r2),(0,5.5))
    plt.annotate('y = {:.3f}'.format(-slp)+'x+({:.3f})'.format(y0),(0,6.5))
    
    # Overlaid plots
    plt.subplot(3,6,p1[n])
    plt.plot(-np.log2(std_crv_x_fit),dCt_Uu_fit_line[n],color = [0.5,0.5,0.5],linestyle = '--',linewidth=1)
    plt.plot(-np.log2(std_crv_x),mean_dCt_Uu_list[n],'o',color='k',markersize=ms)
    plt.plot(-np.log2(mean_fit_clv_gate_list[n]),mean_ddCt_clv_gate_list[n],'o',color=[0,0.5,1],markersize=ms)
    plt.ylim(-1,ymax_stc)
    
    plt.xticks(fontsize=fs)
    plt.yticks(fontsize=fs)
    #plt.xlim(0,ymax)
    ax1 = plt.gca()
    ax1.xaxis.set_tick_params(which='both',size=3,width=1,direction='in',top='on')
    #ax1.xaxis.set_major_locator(MaxNLocator(integer=True))
    ax1.yaxis.set_tick_params(which='both',size=3,width=1,direction='in',right='on')
    plt.xlabel('-log2(Fraction uncleaved)',fontsize=fs)
    plt.ylabel('$\Delta$Ct_Uu',fontsize=fs)
    
'''
PCRo standard curves
'''
plt.figure(5)

for n in range(len(exp_files)):
    
    plt.subplot(3,6,p0[n])
    plt.plot(-np.log2(std_crv_x_fit),dCt_Uo_fit_line[n],color = [0.5,0.5,0.5],linestyle = '--',linewidth=1)
    plt.plot(-np.log2(std_crv_x),mean_dCt_Uo_list[n],'o',color='k',markersize=ms)
    plt.errorbar(-np.log2(std_crv_x),mean_dCt_Uo_list[n],yerr=std_dCt_Uo_list[n],capsize=cs,color=[1,0,1],fmt='none')
    
    plt.plot(-np.log2(mean_fit_clv_gate_list[n]),mean_ddCt_clv_gate_list[n],'o',color=[0,0.5,1],markersize=ms)
    plt.errorbar(-np.log2(mean_fit_clv_gate_list[n]),mean_ddCt_clv_gate_list[n],yerr=std_ddCt_clv_gate_list[n],capsize=cs,color=[1,0,1],fmt='none')
    plt.ylim(-1,ymax_stc)
    
    if plot0 == 1:
        plt.plot([0],[0],'o',color='k',markersize=ms)
    
    plt.xticks(fontsize=fs)
    plt.yticks(fontsize=fs)
    #plt.xlim(0,ymax)
    ax1 = plt.gca()
    ax1.xaxis.set_tick_params(which='both',size=3,width=1,direction='in',top='on')
    #ax1.xaxis.set_major_locator(MaxNLocator(integer=True))
    ax1.yaxis.set_tick_params(which='both',size=3,width=1,direction='in',right='on')
    plt.xlabel('-log2(Fraction uncleaved)',fontsize=fs)
    plt.ylabel('$\Delta$Ct_Uo',fontsize=fs)   
    r2 = exp_files['fit_line_par_o'][n][-1]
    slp = exp_files['fit_line_par_o'][n][0]
    y0 = exp_files['fit_line_par_o'][n][1]
    plt.annotate('R^2 = {:.3f}'.format(r2),(0,5.5))
    plt.annotate('y = {:.3f}'.format(-slp)+'x+({:.3f})'.format(y0),(0,6.5))
    
    # Overlaid plots
    plt.subplot(3,6,p1[n])
    plt.plot(-np.log2(std_crv_x_fit),dCt_Uo_fit_line[n],color = [0.5,0.5,0.5],linestyle = '--',linewidth=1)
    plt.plot(-np.log2(std_crv_x),mean_dCt_Uo_list[n],'o',color='k',markersize=ms)
    plt.plot(-np.log2(mean_fit_clv_gate_list[n]),mean_ddCt_clv_gate_list[n],'o',color=[0,0.5,1],markersize=ms)
    plt.ylim(-1,ymax_stc)
    
    plt.xticks(fontsize=fs)
    plt.yticks(fontsize=fs)
    #plt.xlim(0,ymax)
    ax1 = plt.gca()
    ax1.xaxis.set_tick_params(which='both',size=3,width=1,direction='in',top='on')
    #ax1.xaxis.set_major_locator(MaxNLocator(integer=True))
    ax1.yaxis.set_tick_params(which='both',size=3,width=1,direction='in',right='on')
    plt.xlabel('-log2(Fraction uncleaved)',fontsize=fs)
    plt.ylabel('$\Delta$Ct_Uo',fontsize=fs)


# %%
# defining the whole plate to pull in
plate_positions = []
well_hash = {}
wrow = ['A','B','C','D','E','F','G','H']
wcol = [i+1 for i in range(12)]
for n in range(len(wrow)):
    for m in range(len(wcol)):
        well_hash[wrow[n]+str(wcol[m])] = m+12*n+1

well_index_list = []
results_df_list = []

C_pos_list = []
G_pos_list = []
C_list = []
n_samples_list = []

for f in range(len(exp_files)):
    
    folder_path = exp_files['folder_path'][f]+'\\'
    file_name = exp_files['file_name'][f]
    file_path = results_path+folder_path+file_name
    
    thld = exp_files['thld'][f]  # threshold value for experiment
    n_samples = exp_files['n_samples'][f]  
    G_pos = exp_files['G_pos'][f]  # column of plate G is in
    C_pos = exp_files['C_pos'][f]  # column of plate C is in 
    U_pos = 1 # column of plate U is in
    C = exp_files['C'][f]  # cleaved gate present? 1 or 0

    '''
    Well and Ct data
    '''
    # reading in data
    results_data = pd.read_excel(file_path,sheet_name='Results',skiprows=35)
    results_df = pd.DataFrame(data=results_data,columns=['Well','Well Position','CT','Ct Threshold'])[:-5]
    plate_positions = results_df['Well Position']
    well_numbers = [well_hash[plate_positions[n]] for n in range(len(plate_positions))]
    
    '''
    Amplification data
    '''
    Rn_array = []
    rn_data = pd.read_excel(file_path,sheet_name='Amplification Data',skiprows=35)
    Rn_df = pd.DataFrame(data=rn_data,columns=['Well','Cycle',rn_type])
    Rn1 = Rn_df.loc[Rn_df['Well'] == 1]
    Rn_cycles = pd.DataFrame(data=Rn1,columns=['Cycle'])
    
    for m in range(len(well_numbers)):
        Rn = Rn_df.loc[Rn_df['Well'] == well_numbers[m]]
        Rn_array.append(pd.DataFrame(data=Rn,columns=[rn_type]))
        
    results_df['Rn_data'] = Rn_array
    
    '''
    Melt curves data
    '''
    
    melt_curves = []
    
    # reading in data
    mcp_data = pd.read_excel(file_path,sheet_name='Multicomponent Data',skiprows=35)
        
    for m in range(len(plate_positions)):
        melt_curves.append(np.diff(-mcp_data[mcp_data['Well Position'] == plate_positions[m]]['SYBR']))
    
    # x axis for melt curves (hard coded)
    mc_xdata = (np.array([n for n in range(257)])-2*no_PCR_cycles)/5.11+60
    
    
    results_df['melt_data'] = melt_curves
    
    # removing spaces in the column headers
    results_df.columns = [c.replace(' ', '_') for c in results_df.columns]
    
    well_index = {}
    for i in range(len(results_df.Well_Position)):
        well_index[results_df.Well_Position[i]]=i
        

    # saving data for plotting each time through
    results_df_list.append(results_df)
    well_index_list.append(well_index)
    C_pos_list.append(C_pos)
    G_pos_list.append(G_pos)
    C_list.append(C)
    n_samples_list.append(n_samples)
    

'''
##############################################################################
USER DEFINED PLOTTING 
##############################################################################
Some notes
There are a few different ways to pull stuff out of the dataframe

It can be by index if you keep track of which index is in which well:
results_df.Rn_data[0]

It can be by well name through the following command:
results_df.Rn_data[results_df.Well_Position=='A1'].values

Lastly, it can be well name pulled out of the well_index dictionary above:
results_df.Rn_data[well_index['A1']]


'''

# colors for lines
cpU = [[0,0.7,0],
       [0,0.7,0],
       [0,0.7,0]]

cpG = [[0,0,1],
       [0,0,1],
       [0,0,1]]

cpC = [[0.6,0.6,0.6],
       [0.6,0.6,0.6],
       [0.6,0.6,0.6]]


'''
Plotting raw amplification curves
'''
fs = 12 # fontsize for plots
plotting_scale = 'log' # can also be 'linear'
ymin = 0.01 # min value on y axis
ymax = 35 # max value on y axis
lw = 1 # linewidth
pg = [1,2,13,14] 
tech_reps = 2


plt.figure()
plt.subplot(3,6,1)
plt.title('PCRo')
plt.plot(Rn_cycles,results_df_list[0].Rn_data[well_index['A1']],color=[0,0.6,0],linewidth=lw) # uncleaved
plt.plot(Rn_cycles,results_df_list[0].Rn_data[well_index['B1']],color=[0,0.6,0],linewidth=lw) 
plt.plot(Rn_cycles,results_df_list[1].Rn_data[well_index['C1']],color=[0.6,0.6,0.6],linewidth=lw) # uncleaved
plt.plot(Rn_cycles,results_df_list[1].Rn_data[well_index['D1']],color=[0.6,0.6,0.6],linewidth=lw)
plt.plot(Rn_cycles,results_df_list[0].Rn_data[well_index['A8']],color=[0,0,1],linewidth=lw) # uncleaved
plt.plot(Rn_cycles,results_df_list[0].Rn_data[well_index['B8']],color=[0,0,1],linewidth=lw)

plt.plot(Rn_cycles,results_df_list[0].Rn_data[well_index['A7']],color=[0,0,0],linewidth=lw) # uncleaved
plt.plot(Rn_cycles,results_df_list[0].Rn_data[well_index['B7']],color=[0,0,0],linewidth=lw)

plt.plot(Rn_cycles,results_df_list[0].Rn_data[well_index['A9']],color=[0,0,0],linestyle=':',linewidth=lw) # negative control
plt.plot([0,40],[results_df_list[0].Ct_Threshold[0],results_df_list[0].Ct_Threshold[0]],color=[1,0,0],linestyle='-',linewidth=0.5) # threshold line
plt.xticks(fontsize=fs)
plt.yticks(fontsize=fs)
plt.ylim(0.01,ymax)
plt.xlim(0,40)
ax1 = plt.gca()
ax1.xaxis.set_tick_params(which='both',size=3,width=1,direction='in',top='on')
#ax1.xaxis.set_major_locator(MaxNLocator(integer=True))
ax1.yaxis.set_tick_params(which='both',size=3,width=1,direction='in',right='on')
ax1.set_yscale(plotting_scale)
plt.xlabel('Cycle #',fontsize=fs)
plt.ylabel('log(Rn)',fontsize=fs)


plt.subplot(3,6,13)
plt.title('PCRu')
plt.plot(Rn_cycles,results_df_list[0].Rn_data[well_index['E1']],color=[0,0.6,0],linewidth=lw) # uncleaved
plt.plot(Rn_cycles,results_df_list[0].Rn_data[well_index['F1']],color=[0,0.6,0],linewidth=lw) 
plt.plot(Rn_cycles,results_df_list[0].Rn_data[well_index['E8']],color=[0,0,1],linewidth=lw) # uncleaved
plt.plot(Rn_cycles,results_df_list[0].Rn_data[well_index['F8']],color=[0,0,1],linewidth=lw)
plt.plot(Rn_cycles,results_df_list[0].Rn_data[well_index['E7']],color=[0,0,0],linewidth=lw) # uncleaved
plt.plot(Rn_cycles,results_df_list[0].Rn_data[well_index['F7']],color=[0,0,0],linewidth=lw)

plt.plot(Rn_cycles,results_df_list[0].Rn_data[well_index['E9']],color=[0,0,0],linestyle=':',linewidth=lw) # negative control
plt.plot([0,40],[results_df_list[0].Ct_Threshold[0],results_df_list[0].Ct_Threshold[0]],color=[1,0,0],linestyle='-',linewidth=0.5) # threshold line 
plt.xticks(fontsize=fs)
plt.yticks(fontsize=fs)
plt.ylim(0.01,ymax)
plt.xlim(0,40)
ax1 = plt.gca()
ax1.xaxis.set_tick_params(which='both',size=3,width=1,direction='in',top='on')
#ax1.xaxis.set_major_locator(MaxNLocator(integer=True))
ax1.yaxis.set_tick_params(which='both',size=3,width=1,direction='in',right='on')
ax1.set_yscale(plotting_scale)
plt.xlabel('Cycle #',fontsize=fs)
plt.ylabel('log(Rn)',fontsize=fs)
```
