## Supplementary material for "Prevention of ribozyme catalysis through cDNA synthesis enables accurate RT-qPCR measurements of context-dependent ribozyme activity": Uncropped gel images, raw data, and analysis code: Supp_Figures_S4_S5_S6.py

```
import numpy as np
import matplotlib.pyplot as plt
from matplotlib.ticker import MaxNLocator
import openpyxl as opxl
import pandas as pd
import warnings
import copy
import scipy.interpolate as scipy_int
from scipy.signal import savgol_filter
warnings.filterwarnings("ignore", \
message="Adding an axes using the same arguments as a previous axes currently \
reuses the earlier instance.  In a future version, a new instance will always be \
created and returned.  Meanwhile, this warning can be suppressed, and the future \
behavior ensured, by passing a unique label to each axes instance.")

from RT_qPCR_Rz_analysis_ave3 import RT_qPCR_Rz_analysis_ave3

'''
##############################################################################
USER DEFINED FILE LOCATION AND PLATE POSITIONS FOR IMPORTING
##############################################################################
'''
results_path = 'C:\\Users\\sws5\\OneDrive - National Institute of Standards and Technology (NIST)\\Presentations and write ups\\Papers\\ctRSD RT qPCR\\Data for upload\\'
exp_path = 'C:\\Users\\sws5\\OneDrive - National Institute of Standards and Technology (NIST)\\Presentations and write ups\\Papers\\ctRSD RT qPCR\\Data for upload\\list_experiments_paths_final.xlsx'

exp_files = pd.read_excel(exp_path,sheet_name='Supp Figs 4 5 6')
show_ideal = 1
show_amp = [0,0,0] 

no_PCR_cycles = 40
rn_type = 'Delta Rn' # which data type to import (can also be raw Rn)
tech_reps = 3 # number of technical replicates

    
# defining the whole plate to pull in
plate_positions = []
well_hash = {}
wrow = ['A','B','C','D','E','F','G','H']
wcol = [i+1 for i in range(12)]
for n in range(len(wrow)):
    for m in range(len(wcol)):
        well_hash[wrow[n]+str(wcol[m])] = m+12*n+1

well_index_list = []
results_df_list = []

C_pos_list = []
G_pos_list = []
C_list = []
n_samples_list = []


for f in range(len(exp_files)):
    
    folder_path = exp_files['folder_path'][f]+'\\'
    file_name = exp_files['file_name'][f]
    file_path = results_path+folder_path+file_name
    
    thld = exp_files['thld'][f]  # threshold value for experiment
    n_samples = exp_files['n_samples'][f]  
    G_pos = exp_files['G_pos'][f]  # column of plate G is in
    C_pos = exp_files['C_pos'][f]  # column of plate C is in 
    U_pos = 1 # column of plate U is in
    C = exp_files['C'][f]  # cleaved gate present? 1 or 0

    '''
    Well and Ct data
    '''
    # reading in data
    results_data = pd.read_excel(file_path,sheet_name='Results',skiprows=35)
    results_df = pd.DataFrame(data=results_data,columns=['Well','Well Position','CT','Ct Threshold'])[:-5]
    plate_positions = results_df['Well Position']
    well_numbers = [well_hash[plate_positions[n]] for n in range(len(plate_positions))]
    
    '''
    Amplification data
    '''
    Rn_array = []
    rn_data = pd.read_excel(file_path,sheet_name='Amplification Data',skiprows=35)
    Rn_df = pd.DataFrame(data=rn_data,columns=['Well','Cycle',rn_type])
    Rn1 = Rn_df.loc[Rn_df['Well'] == 1]
    Rn_cycles = pd.DataFrame(data=Rn1,columns=['Cycle'])
    
    for m in range(len(well_numbers)):
        Rn = Rn_df.loc[Rn_df['Well'] == well_numbers[m]]
        Rn_array.append(pd.DataFrame(data=Rn,columns=[rn_type]))
        
    results_df['Rn_data'] = Rn_array
    
    '''
    Melt curves data
    '''
    
    melt_curves = []
    melt_curves1 = []
    melt_curves_CS = []
    melt_curves_d = []
    
    # reading in data
    mcp_data = pd.read_excel(file_path,sheet_name='Multicomponent Data',skiprows=35)
   
    # x axis for melt curves (hard coded)
    mc_xdata1 = (np.array([n for n in range(258)])-2*no_PCR_cycles)/5.11+60
    mc_xdata = mc_xdata1[80:]
    mc_xdata_cs = np.arange(60,95,0.05)
        
    for m in range(len(plate_positions)):
        melt_curves1.append((mcp_data[mcp_data['Well Position'] == plate_positions[m]]['SYBR']))
        #melt_curves1.append((mcp_data[mcp_data['Well Position'] == plate_positions[m]]['SYBR']/mcp_data[mcp_data['Well Position'] == plate_positions[m]]['ROX']))
        melt_curves.append(savgol_filter(melt_curves1[m][81:], 11, 3))
        melt_curves_d.append(np.diff(-melt_curves[m]))
    
    
    results_df['melt_data'] = melt_curves_d
    
    # removing spaces in the column headers
    results_df.columns = [c.replace(' ', '_') for c in results_df.columns]
    
    well_index = {}
    for i in range(len(results_df.Well_Position)):
        well_index[results_df.Well_Position[i]]=i
        

    # saving data for plotting each time through
    results_df_list.append(results_df)
    well_index_list.append(well_index)
    C_pos_list.append(C_pos)
    G_pos_list.append(G_pos)
    C_list.append(C)
    n_samples_list.append(n_samples)
    
    plt.figure()
    plt.subplot(2,2,1)
    plt.plot(mc_xdata,melt_curves1[0][81:],color=[0,0.8,0],marker='o',linestyle='')
    plt.plot(mc_xdata,melt_curves[0],color=[0,0.8,0])
    plt.xlim(60,95)
    
    plt.subplot(2,2,2)
    plt.plot(mc_xdata[:-1],np.diff(-melt_curves1[0][81:]),color=[0,0.8,0])
    plt.plot(mc_xdata[:-1],np.diff(-melt_curves[0]),color=[0,0,0])
    plt.xlim(60,95)
    

# %%  
'''
##############################################################################
USER DEFINED PLOTTING 
##############################################################################
Some notes
There are a few different ways to pull stuff out of the dataframe

It can be by index if you keep track of which index is in which well:
results_df.Rn_data[0]

It can be by well name through the following command:
results_df.Rn_data[results_df.Well_Position=='A1'].values

Lastly, it can be well name pulled out of the well_index dictionary above:
results_df.Rn_data[well_index['A1']]


'''

# colors for lines
cpU = [[0,0.7,0],
       [0,0.7,0],
       [0,0.7,0]]

cpG = [[0,0,1],
       [0,0,1],
       [0,0,1]]

cpC = [[0.6,0.6,0.6],
       [0.6,0.6,0.6],
       [0.6,0.6,0.6]]

index = 0
well_index = well_index_list[index]

'''
Plotting raw amplification curves
'''
fs = 12 # fontsize for plots
plotting_scale = 'log' # can also be 'linear'
ymin = 0.01 # min value on y axis
ymax = 25 # max value on y axis
lw = 1 # linewidth

rep_row = ['A','B','C'] # rows of replicates
plt.subplot(2,3,1)
plt.title('G2 PCRo')
for n in range(tech_reps):
    plt.plot(Rn_cycles,results_df_list[index].Rn_data[well_index[rep_row[n]+str(U_pos)]],color=cpU[n],linewidth=lw) # uncleaved
    plt.plot(Rn_cycles,results_df_list[index].Rn_data[well_index[rep_row[n]+str(G_pos_list[index])]],color=cpG[n],linewidth=lw) # gate
    if C_list[index] == 1:
        plt.plot(Rn_cycles,results_df_list[index].Rn_data[well_index[rep_row[n]+str(C_pos_list[index])]],color=cpC[n],linewidth=lw) # cleaved

plt.plot(Rn_cycles,results_df_list[index].Rn_data[well_index['B10']],color=[0,0,0],linestyle=':',linewidth=lw) # negative control
plt.plot(Rn_cycles,results_df_list[index].Rn_data[well_index['A10']],color=[0,0,1],linestyle='--',linewidth=lw) # positive control
plt.plot([0,40],[results_df_list[index].Ct_Threshold[0],results_df_list[index].Ct_Threshold[0]],color=[1,0,0],linestyle='-',linewidth=0.5) # threshold line
plt.xticks(fontsize=fs)
plt.yticks(fontsize=fs)
plt.ylim(0.01,ymax)
plt.xlim(0,40)
ax1 = plt.gca()
ax1.xaxis.set_tick_params(which='both',size=3,width=1,direction='in',top='on')
#ax1.xaxis.set_major_locator(MaxNLocator(integer=True))
ax1.yaxis.set_tick_params(which='both',size=3,width=1,direction='in',right='on')
ax1.set_yscale(plotting_scale)
plt.xlabel('Cycle #',fontsize=fs)
plt.ylabel('log(Rn)',fontsize=fs)

rep_row = ['D','E','F'] # rows of replicates
plt.subplot(2,3,4)
plt.title('G2 PCRu')
for n in range(tech_reps):
    plt.plot(Rn_cycles,results_df_list[index].Rn_data[well_index[rep_row[n]+str(U_pos)]],color=cpU[n],linewidth=lw) # uncleaved
    plt.plot(Rn_cycles,results_df_list[index].Rn_data[well_index[rep_row[n]+str(G_pos_list[index])]],color=cpG[n],linewidth=lw) # gate
    plt.plot(Rn_cycles,results_df_list[index].Rn_data[well_index[rep_row[n]+str(G_pos_list[index]-2)]],color=[0,0,0],linewidth=lw) # gate
    if C_list[index] == 1:
        plt.plot(Rn_cycles,results_df_list[index].Rn_data[well_index[rep_row[n]+str(C_pos_list[index])]],color=cpC[n],linewidth=lw) # cleaved

plt.plot(Rn_cycles,results_df_list[index].Rn_data[well_index['E10']],color=[0,0,0],linestyle=':',linewidth=lw) # negative control
plt.plot(Rn_cycles,results_df_list[index].Rn_data[well_index['D10']],color=[0,0,1],linestyle='--',linewidth=lw) # positive control
plt.plot([0,40],[results_df_list[index].Ct_Threshold[0],results_df_list[index].Ct_Threshold[0]],color=[1,0,0],linestyle='-',linewidth=0.5) # threshold line
plt.xticks(fontsize=fs)
plt.yticks(fontsize=fs)
plt.ylim(0.01,ymax)
plt.xlim(0,40)
ax1 = plt.gca()
ax1.xaxis.set_tick_params(which='both',size=3,width=1,direction='in',top='on')
#ax1.xaxis.set_major_locator(MaxNLocator(integer=True))
ax1.yaxis.set_tick_params(which='both',size=3,width=1,direction='in',right='on')
ax1.set_yscale(plotting_scale)
plt.xlabel('Cycle #',fontsize=fs)
plt.ylabel('log(Rn)',fontsize=fs)


# CELLS
index = 2
well_index = well_index_list[index]

'''
Plotting raw amplification curves
'''
fs = 12 # fontsize for plots
plotting_scale = 'log' # can also be 'linear'
ymin = 0.01 # min value on y axis
ymax = 25 # max value on y axis
lw = 1 # linewidth

rep_row = ['A','B','C'] # rows of replicates
plt.subplot(2,3,2)
plt.title('G2 PCRo')
for n in range(tech_reps):
    plt.plot(Rn_cycles,results_df_list[index].Rn_data[well_index[rep_row[n]+str(U_pos)]],color=cpU[n],linewidth=lw) # uncleaved
    plt.plot(Rn_cycles,results_df_list[index].Rn_data[well_index[rep_row[n]+str(G_pos_list[index])]],color=cpG[n],linewidth=lw) # gate
    if C_list[index] == 1:
        plt.plot(Rn_cycles,results_df_list[index].Rn_data[well_index[rep_row[n]+str(C_pos_list[index])]],color=cpC[n],linewidth=lw) # cleaved

plt.plot(Rn_cycles,results_df_list[index].Rn_data[well_index['A10']],color=[0,0,0],linestyle=':',linewidth=lw) # negative control
plt.plot(Rn_cycles,results_df_list[index].Rn_data[well_index['A11']],color=[0,0.6,0],linestyle='--',linewidth=lw) # positive control
plt.plot(Rn_cycles,results_df_list[index].Rn_data[well_index['B11']],color=[0.6,0.6,0.6],linestyle='--',linewidth=lw) # positive control
plt.plot(Rn_cycles,results_df_list[index].Rn_data[well_index['C11']],color=[0,0,1],linestyle='--',linewidth=lw) # positive control
plt.plot([0,40],[results_df_list[index].Ct_Threshold[0],results_df_list[index].Ct_Threshold[0]],color=[1,0,0],linestyle='-',linewidth=0.5) # threshold line
plt.xticks(fontsize=fs)
plt.yticks(fontsize=fs)
plt.ylim(0.01,ymax)
plt.xlim(0,40)
ax1 = plt.gca()
ax1.xaxis.set_tick_params(which='both',size=3,width=1,direction='in',top='on')
#ax1.xaxis.set_major_locator(MaxNLocator(integer=True))
ax1.yaxis.set_tick_params(which='both',size=3,width=1,direction='in',right='on')
ax1.set_yscale(plotting_scale)
plt.xlabel('Cycle #',fontsize=fs)
plt.ylabel('log(Rn)',fontsize=fs)

rep_row = ['D','E','F'] # rows of replicates
plt.subplot(2,3,5)
plt.title('G2 PCRu')
for n in range(tech_reps):
    plt.plot(Rn_cycles,results_df_list[index].Rn_data[well_index[rep_row[n]+str(U_pos)]],color=cpU[n],linewidth=lw) # uncleaved
    plt.plot(Rn_cycles,results_df_list[index].Rn_data[well_index[rep_row[n]+str(G_pos_list[index])]],color=cpG[n],linewidth=lw) # gate
    plt.plot(Rn_cycles,results_df_list[index].Rn_data[well_index[rep_row[n]+str(G_pos_list[index]-2)]],color=[0,0,0],linewidth=lw) # gate
    if C_list[index] == 1:
        plt.plot(Rn_cycles,results_df_list[index].Rn_data[well_index[rep_row[n]+str(C_pos_list[index])]],color=cpC[n],linewidth=lw) # cleaved

plt.plot(Rn_cycles,results_df_list[index].Rn_data[well_index['D10']],color=[0,0,0],linestyle=':',linewidth=lw) # negative control
plt.plot(Rn_cycles,results_df_list[index].Rn_data[well_index['D11']],color=[0,0.6,0],linestyle='--',linewidth=lw) # positive control
plt.plot(Rn_cycles,results_df_list[index].Rn_data[well_index['E11']],color=[0.6,0.6,0.6],linestyle='--',linewidth=lw) # positive control
plt.plot(Rn_cycles,results_df_list[index].Rn_data[well_index['F11']],color=[0,0,1],linestyle='--',linewidth=lw) # positive control
plt.plot([0,40],[results_df_list[index].Ct_Threshold[0],results_df_list[index].Ct_Threshold[0]],color=[1,0,0],linestyle='-',linewidth=0.5) # threshold line
plt.xticks(fontsize=fs)
plt.yticks(fontsize=fs)
plt.ylim(0.01,ymax)
plt.xlim(0,40)
ax1 = plt.gca()
ax1.xaxis.set_tick_params(which='both',size=3,width=1,direction='in',top='on')
#ax1.xaxis.set_major_locator(MaxNLocator(integer=True))
ax1.yaxis.set_tick_params(which='both',size=3,width=1,direction='in',right='on')
ax1.set_yscale(plotting_scale)
plt.xlabel('Cycle #',fontsize=fs)
plt.ylabel('log(Rn)',fontsize=fs)


# G2h
index = 3
well_index = well_index_list[index]

'''
Plotting raw amplification curves
'''
fs = 12 # fontsize for plots
plotting_scale = 'log' # can also be 'linear'
ymin = 0.01 # min value on y axis
ymax = 25 # max value on y axis
lw = 1 # linewidth

rep_row = ['A','B','C'] # rows of replicates
plt.subplot(2,3,3)
plt.title('G2h PCRo')
for n in range(tech_reps):
    plt.plot(Rn_cycles,results_df_list[index].Rn_data[well_index[rep_row[n]+str(U_pos)]],color=cpU[n],linewidth=lw) # uncleaved
    plt.plot(Rn_cycles,results_df_list[index].Rn_data[well_index[rep_row[n]+str(G_pos_list[index])]],color=cpG[n],linewidth=lw) # gate
    if C_list[index] == 1:
        plt.plot(Rn_cycles,results_df_list[index].Rn_data[well_index[rep_row[n]+str(C_pos_list[index])]],color=cpC[n],linewidth=lw) # cleaved

plt.plot(Rn_cycles,results_df_list[index].Rn_data[well_index['A11']],color=[0,0,0],linestyle=':',linewidth=lw) # negative control
plt.plot(Rn_cycles,results_df_list[index].Rn_data[well_index['A10']],color=[0,0,1],linestyle='--',linewidth=lw) # positive control
plt.plot([0,40],[results_df_list[index].Ct_Threshold[0],results_df_list[index].Ct_Threshold[0]],color=[1,0,0],linestyle='-',linewidth=0.5) # threshold line
plt.xticks(fontsize=fs)
plt.yticks(fontsize=fs)
plt.ylim(0.01,ymax)
plt.xlim(0,40)
ax1 = plt.gca()
ax1.xaxis.set_tick_params(which='both',size=3,width=1,direction='in',top='on')
#ax1.xaxis.set_major_locator(MaxNLocator(integer=True))
ax1.yaxis.set_tick_params(which='both',size=3,width=1,direction='in',right='on')
ax1.set_yscale(plotting_scale)
plt.xlabel('Cycle #',fontsize=fs)
plt.ylabel('log(Rn)',fontsize=fs)

rep_row = ['D','E','F'] # rows of replicates
plt.subplot(2,3,6)
plt.title('G2h PCRu')
for n in range(tech_reps):
    plt.plot(Rn_cycles,results_df_list[index].Rn_data[well_index[rep_row[n]+str(U_pos)]],color=cpU[n],linewidth=lw) # uncleaved
    plt.plot(Rn_cycles,results_df_list[index].Rn_data[well_index[rep_row[n]+str(G_pos_list[index])]],color=cpG[n],linewidth=lw) # gate
    plt.plot(Rn_cycles,results_df_list[index].Rn_data[well_index[rep_row[n]+str(G_pos_list[index]-2)]],color=[0,0,0],linewidth=lw) # gate
    if C_list[index] == 1:
        plt.plot(Rn_cycles,results_df_list[index].Rn_data[well_index[rep_row[n]+str(C_pos_list[index])]],color=cpC[n],linewidth=lw) # cleaved

plt.plot(Rn_cycles,results_df_list[index].Rn_data[well_index['D11']],color=[0,0,0],linestyle=':',linewidth=lw) # negative control
plt.plot(Rn_cycles,results_df_list[index].Rn_data[well_index['D10']],color=[0,0,1],linestyle='--',linewidth=lw) # positive control
plt.plot([0,40],[results_df_list[index].Ct_Threshold[0],results_df_list[index].Ct_Threshold[0]],color=[1,0,0],linestyle='-',linewidth=0.5) # threshold line
plt.xticks(fontsize=fs)
plt.yticks(fontsize=fs)
plt.ylim(0.01,ymax)
plt.xlim(0,40)
ax1 = plt.gca()
ax1.xaxis.set_tick_params(which='both',size=3,width=1,direction='in',top='on')
#ax1.xaxis.set_major_locator(MaxNLocator(integer=True))
ax1.yaxis.set_tick_params(which='both',size=3,width=1,direction='in',right='on')
ax1.set_yscale(plotting_scale)
plt.xlabel('Cycle #',fontsize=fs)
plt.ylabel('log(Rn)',fontsize=fs)

# %%
# MELTING CURVES
plt.figure()


ymin = -1000
#ymax = 1.5
ymaxo = 60000
ymaxu = 90000
xmin = 60
xmax = 95

index = 0
well_index = well_index_list[index]

rep_row = ['A','B','C'] # rows of replicates
plt.subplot(2,3,1)
plt.title('G2 PCRo')
for n in range(tech_reps):
    plt.plot(mc_xdata[1:],results_df_list[index].melt_data[well_index[rep_row[n]+str(U_pos)]],color=cpU[n],linewidth=lw) # uncleaved
    plt.plot(mc_xdata[1:],results_df_list[index].melt_data[well_index[rep_row[n]+str(G_pos_list[index])]],color=cpG[n],linewidth=lw) # gate
    if C_list[index] == 1:
        plt.plot(mc_xdata[1:],results_df_list[index].melt_data[well_index[rep_row[n]+str(C_pos_list[index])]],color=cpC[n],linewidth=lw) # cleaved

plt.plot(mc_xdata[1:],results_df_list[index].melt_data[well_index['B10']],color=[0,0,0],linestyle=':',linewidth=lw) # negative control
plt.plot(mc_xdata[1:],results_df_list[index].melt_data[well_index['A10']],color=[0,0,1],linestyle='--',linewidth=lw) # positive control
plt.xticks(fontsize=fs)
plt.yticks(fontsize=fs)
plt.ylim(ymin,ymaxo)
plt.xlim(xmin,xmax)
ax1 = plt.gca()
ax1.xaxis.set_tick_params(which='both',size=3,width=1,direction='in',top='on')
#ax1.xaxis.set_major_locator(MaxNLocator(integer=True))
ax1.yaxis.set_tick_params(which='both',size=3,width=1,direction='in',right='on')
plt.xlabel('Temp (C)',fontsize=fs)
plt.ylabel('d(-SYBR)/dC',fontsize=fs)

rep_row = ['D','E','F'] # rows of replicates
plt.subplot(2,3,4)
plt.title('G2 PCRu')
for n in range(tech_reps):
    plt.plot(mc_xdata[1:],results_df_list[index].melt_data[well_index[rep_row[n]+str(U_pos)]],color=cpU[n],linewidth=lw) # uncleaved
    plt.plot(mc_xdata[1:],results_df_list[index].melt_data[well_index[rep_row[n]+str(G_pos_list[index])]],color=cpG[n],linewidth=lw) # gate
    if C_list[index] == 1:
        plt.plot(mc_xdata[1:],results_df_list[index].melt_data[well_index[rep_row[n]+str(C_pos_list[index])]],color=cpC[n],linewidth=lw) # cleaved

plt.plot(mc_xdata[1:],results_df_list[index].melt_data[well_index['E10']],color=[0,0,0],linestyle=':',linewidth=lw) # negative control
plt.plot(mc_xdata[1:],results_df_list[index].melt_data[well_index['D10']],color=[0,0,1],linestyle='--',linewidth=lw) # positive control
plt.xticks(fontsize=fs)
plt.yticks(fontsize=fs)
plt.ylim(ymin,ymaxu)
plt.xlim(xmin,xmax)
ax1 = plt.gca()
ax1.xaxis.set_tick_params(which='both',size=3,width=1,direction='in',top='on')
#ax1.xaxis.set_major_locator(MaxNLocator(integer=True))
ax1.yaxis.set_tick_params(which='both',size=3,width=1,direction='in',right='on')
plt.xlabel('Temp (C)',fontsize=fs)
plt.ylabel('d(-SYBR)/dC',fontsize=fs)


index = 2
well_index = well_index_list[index]

rep_row = ['A','B','C'] # rows of replicates
plt.subplot(2,3,2)
plt.title('G2 PCRo cells')
for n in range(tech_reps):
    plt.plot(mc_xdata[1:],results_df_list[index].melt_data[well_index[rep_row[n]+str(U_pos)]],color=cpU[n],linewidth=lw) # uncleaved
    plt.plot(mc_xdata[1:],results_df_list[index].melt_data[well_index[rep_row[n]+str(G_pos_list[index])]],color=cpG[n],linewidth=lw) # gate
    if C_list[index] == 1:
        plt.plot(mc_xdata[1:],results_df_list[index].melt_data[well_index[rep_row[n]+str(C_pos_list[index])]],color=cpC[n],linewidth=lw) # cleaved

plt.plot(mc_xdata[1:],results_df_list[index].melt_data[well_index['A10']],color=[0,0,0],linestyle=':',linewidth=lw) # negative control
plt.plot(mc_xdata[1:],results_df_list[index].melt_data[well_index['A11']],color=[0,0.6,0],linestyle='--',linewidth=lw) # positive control
plt.plot(mc_xdata[1:],results_df_list[index].melt_data[well_index['B11']],color=[0.6,0.6,0.6],linestyle='--',linewidth=lw) # positive control
plt.plot(mc_xdata[1:],results_df_list[index].melt_data[well_index['C11']],color=[0,0,1],linestyle='--',linewidth=lw) # positive control
plt.xticks(fontsize=fs)
plt.yticks(fontsize=fs)
plt.ylim(ymin,ymaxo)
plt.xlim(xmin,xmax)
ax1 = plt.gca()
ax1.xaxis.set_tick_params(which='both',size=3,width=1,direction='in',top='on')
#ax1.xaxis.set_major_locator(MaxNLocator(integer=True))
ax1.yaxis.set_tick_params(which='both',size=3,width=1,direction='in',right='on')
plt.xlabel('Temp (C)',fontsize=fs)

rep_row = ['D','E','F'] # rows of replicates
plt.subplot(2,3,5)
plt.title('G2 PCRu cells')
for n in range(tech_reps):
    plt.plot(mc_xdata[1:],results_df_list[index].melt_data[well_index[rep_row[n]+str(U_pos)]],color=cpU[n],linewidth=lw) # uncleaved
    plt.plot(mc_xdata[1:],results_df_list[index].melt_data[well_index[rep_row[n]+str(G_pos_list[index])]],color=cpG[n],linewidth=lw) # gate
    if C_list[index] == 1:
        plt.plot(mc_xdata[1:],results_df_list[index].melt_data[well_index[rep_row[n]+str(C_pos_list[index])]],color=cpC[n],linewidth=lw) # cleaved

plt.plot(mc_xdata[1:],results_df_list[index].melt_data[well_index['D10']],color=[0,0,0],linestyle=':',linewidth=lw) # negative control
plt.plot(mc_xdata[1:],results_df_list[index].melt_data[well_index['D11']],color=[0,0.6,0],linestyle='--',linewidth=lw) # positive control
plt.plot(mc_xdata[1:],results_df_list[index].melt_data[well_index['E11']],color=[0.6,0.6,0.6],linestyle='--',linewidth=lw) # positive control
plt.plot(mc_xdata[1:],results_df_list[index].melt_data[well_index['F11']],color=[0,0,1],linestyle='--',linewidth=lw) # positive control
plt.xticks(fontsize=fs)
plt.yticks(fontsize=fs)
plt.ylim(ymin,ymaxu)
plt.xlim(xmin,xmax)
ax1 = plt.gca()
ax1.xaxis.set_tick_params(which='both',size=3,width=1,direction='in',top='on')
#ax1.xaxis.set_major_locator(MaxNLocator(integer=True))
ax1.yaxis.set_tick_params(which='both',size=3,width=1,direction='in',right='on')
plt.xlabel('Temp (C)',fontsize=fs)


index = 3
well_index = well_index_list[index]

rep_row = ['A','B','C'] # rows of replicates
plt.subplot(2,3,3)
plt.title('G2h PCRo')
for n in range(tech_reps):
    plt.plot(mc_xdata[1:],results_df_list[index].melt_data[well_index[rep_row[n]+str(U_pos)]],color=cpU[n],linewidth=lw) # uncleaved
    plt.plot(mc_xdata[1:],results_df_list[index].melt_data[well_index[rep_row[n]+str(G_pos_list[index])]],color=cpG[n],linewidth=lw) # gate
    if C_list[index] == 1:
        plt.plot(mc_xdata[1:],results_df_list[index].melt_data[well_index[rep_row[n]+str(C_pos_list[index])]],color=cpC[n],linewidth=lw) # cleaved

plt.plot(mc_xdata[1:],results_df_list[index].melt_data[well_index['A11']],color=[0,0,0],linestyle=':',linewidth=lw) # negative control
plt.plot(mc_xdata[1:],results_df_list[index].melt_data[well_index['A10']],color=[0,0,1],linestyle='--',linewidth=lw) # positive control
plt.xticks(fontsize=fs)
plt.yticks(fontsize=fs)
plt.ylim(ymin,ymaxo)
plt.xlim(xmin,xmax)
ax1 = plt.gca()
ax1.xaxis.set_tick_params(which='both',size=3,width=1,direction='in',top='on')
#ax1.xaxis.set_major_locator(MaxNLocator(integer=True))
ax1.yaxis.set_tick_params(which='both',size=3,width=1,direction='in',right='on')
plt.xlabel('Temp (C)',fontsize=fs)

rep_row = ['D','E','F'] # rows of replicates
plt.subplot(2,3,6)
plt.title('G2h PCRu')
for n in range(tech_reps):
    plt.plot(mc_xdata[1:],results_df_list[index].melt_data[well_index[rep_row[n]+str(U_pos)]],color=cpU[n],linewidth=lw) # uncleaved
    plt.plot(mc_xdata[1:],results_df_list[index].melt_data[well_index[rep_row[n]+str(G_pos_list[index])]],color=cpG[n],linewidth=lw) # gate
    if C_list[index] == 1:
        plt.plot(mc_xdata[1:],results_df_list[index].melt_data[well_index[rep_row[n]+str(C_pos_list[index])]],color=cpC[n],linewidth=lw) # cleaved

plt.plot(mc_xdata[1:],results_df_list[index].melt_data[well_index['D11']],color=[0,0,0],linestyle=':',linewidth=lw) # negative control
plt.plot(mc_xdata[1:],results_df_list[index].melt_data[well_index['D10']],color=[0,0,1],linestyle='--',linewidth=lw) # positive control
plt.xticks(fontsize=fs)
plt.yticks(fontsize=fs)
plt.ylim(ymin,ymaxu)
plt.xlim(xmin,xmax)
ax1 = plt.gca()
ax1.xaxis.set_tick_params(which='both',size=3,width=1,direction='in',top='on')
#ax1.xaxis.set_major_locator(MaxNLocator(integer=True))
ax1.yaxis.set_tick_params(which='both',size=3,width=1,direction='in',right='on')
plt.xlabel('Temp (C)',fontsize=fs)
```
