## Supplementary material for "Prevention of ribozyme catalysis through cDNA synthesis enables accurate RT-qPCR measurements of context-dependent ribozyme activity": Uncropped gel images, raw data, and analysis code: README.txt

--------------
UPLOADED DATA:
--------------
Black boxes in gel images show the region that was cropped for use in figures
Note that part the gel in Supplementary Figure 17 is also presented in Figure 3 of the main text

Excel sheets contain the data exported from the Applied Biosystems ViiA 7 Real-Time PCR System
"Results" tab includes:
 - Ct values used for data analysis
 - Ct threshold values 
 - Sample name with percentages rather than fractions for the RNA ratio (i.e, U-100 = U-1 and U-75 = U-0.75)
"Amplification Data" tab
 - Contains the Rn data used to produce amplification plots
"Multicomponent Data" tab
 - Includes the SYBR melt curve data. There are 260 cycles of data here: The first 80 cycles are from qPCR and the remaining 180 are the melt curve data

The "list_experiments_paths_final.xlsx" file is used by the Python analysis scripts to locate the relevant data files for plotting

-------------------------------
UPLOADED DATA ANALYSIS SCRIPTS:
-------------------------------
Python analysis scripts were run with Spyder 5.4.3 on Python 3.8.18
 - RT_qPCR_Rz_analysis_ave3.py contains the data import and analysis code
 - All other scripts produce the plots for the figures
