## Supplementary figures and images for "Prevention of ribozyme catalysis through cDNA synthesis enables accurate RT-qPCR measurements of context-dependent ribozyme activity"

### Figure1D.png

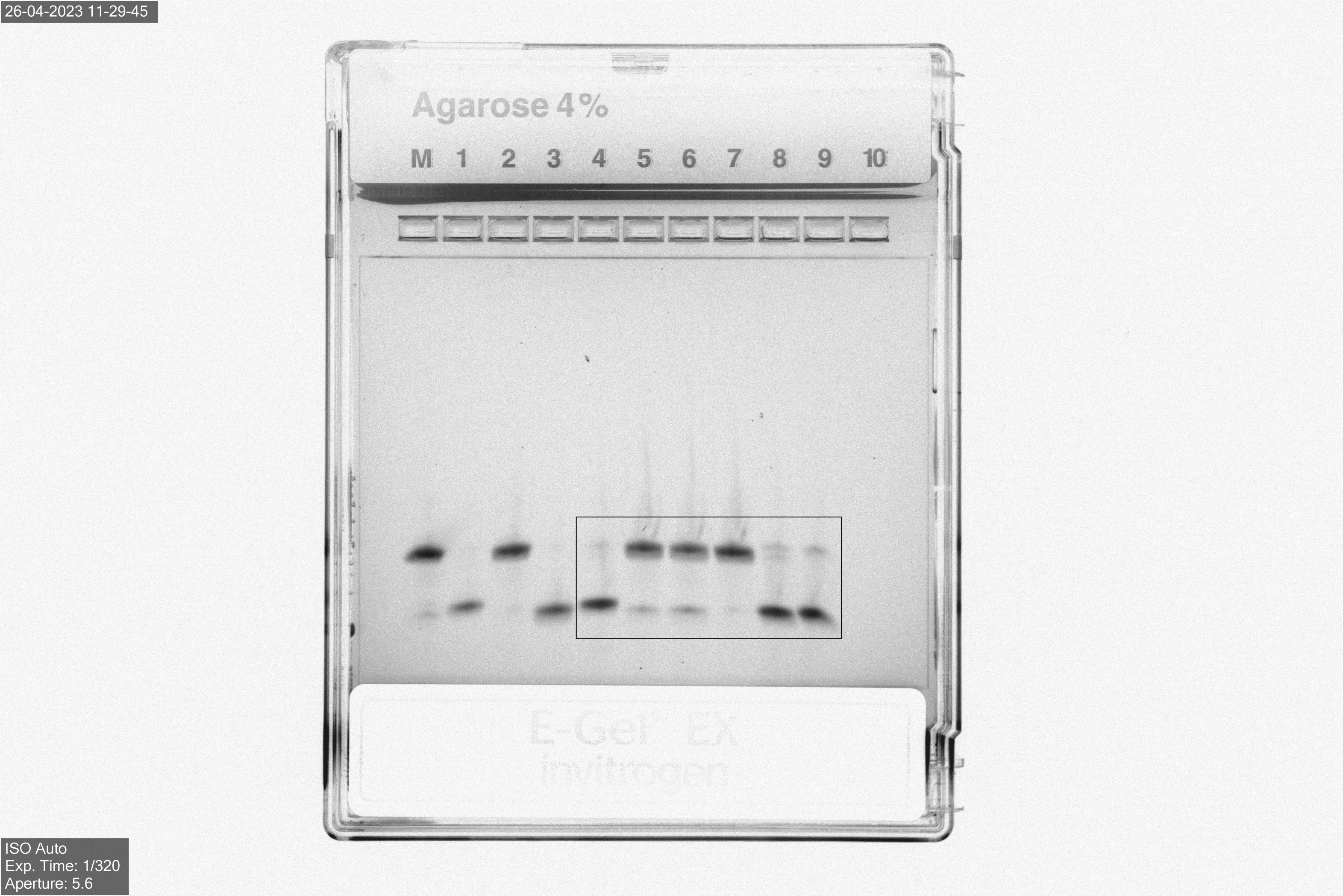

### Figure2_37_60_gel.png

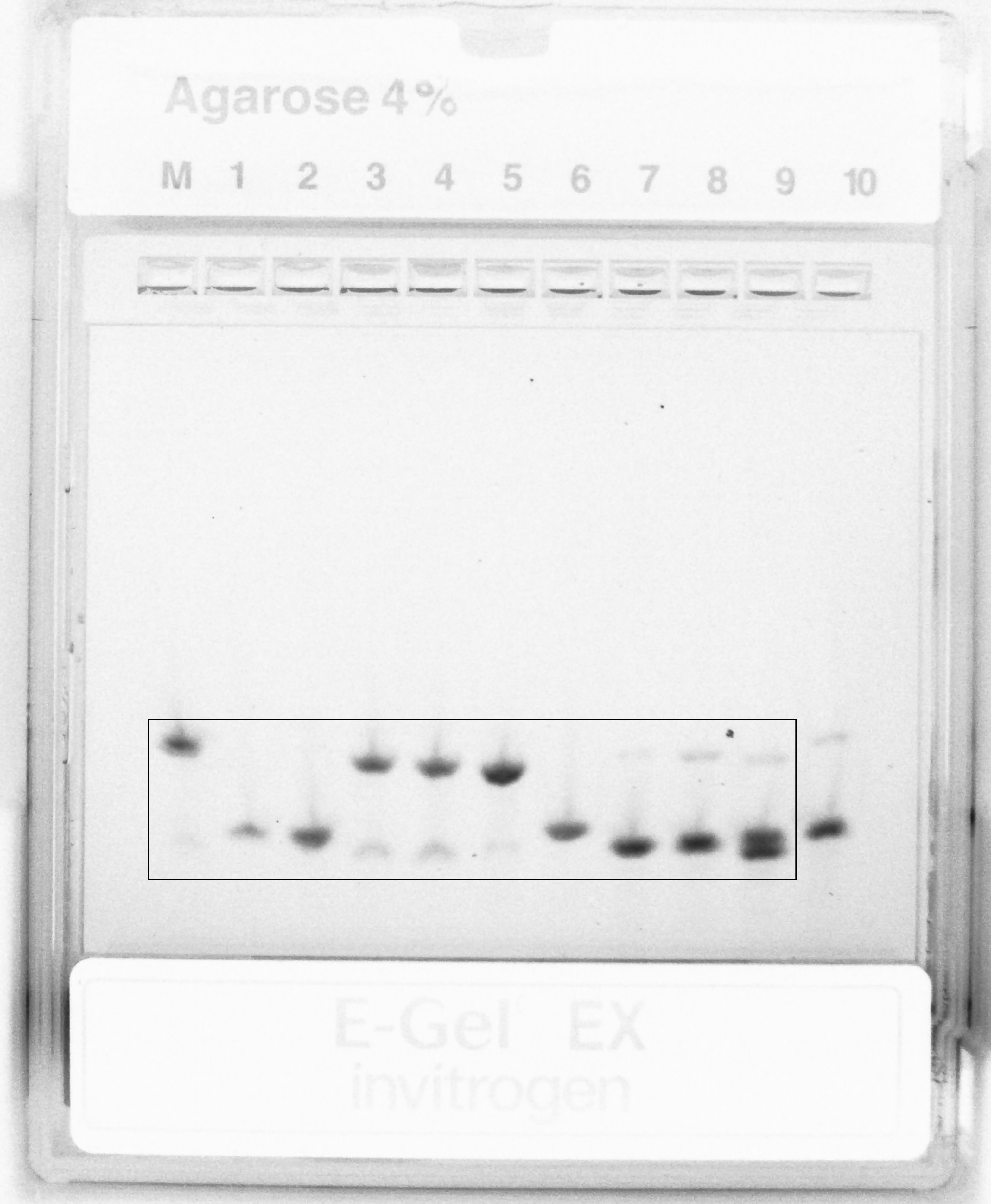

### Figure2_G1_G2_gel.png

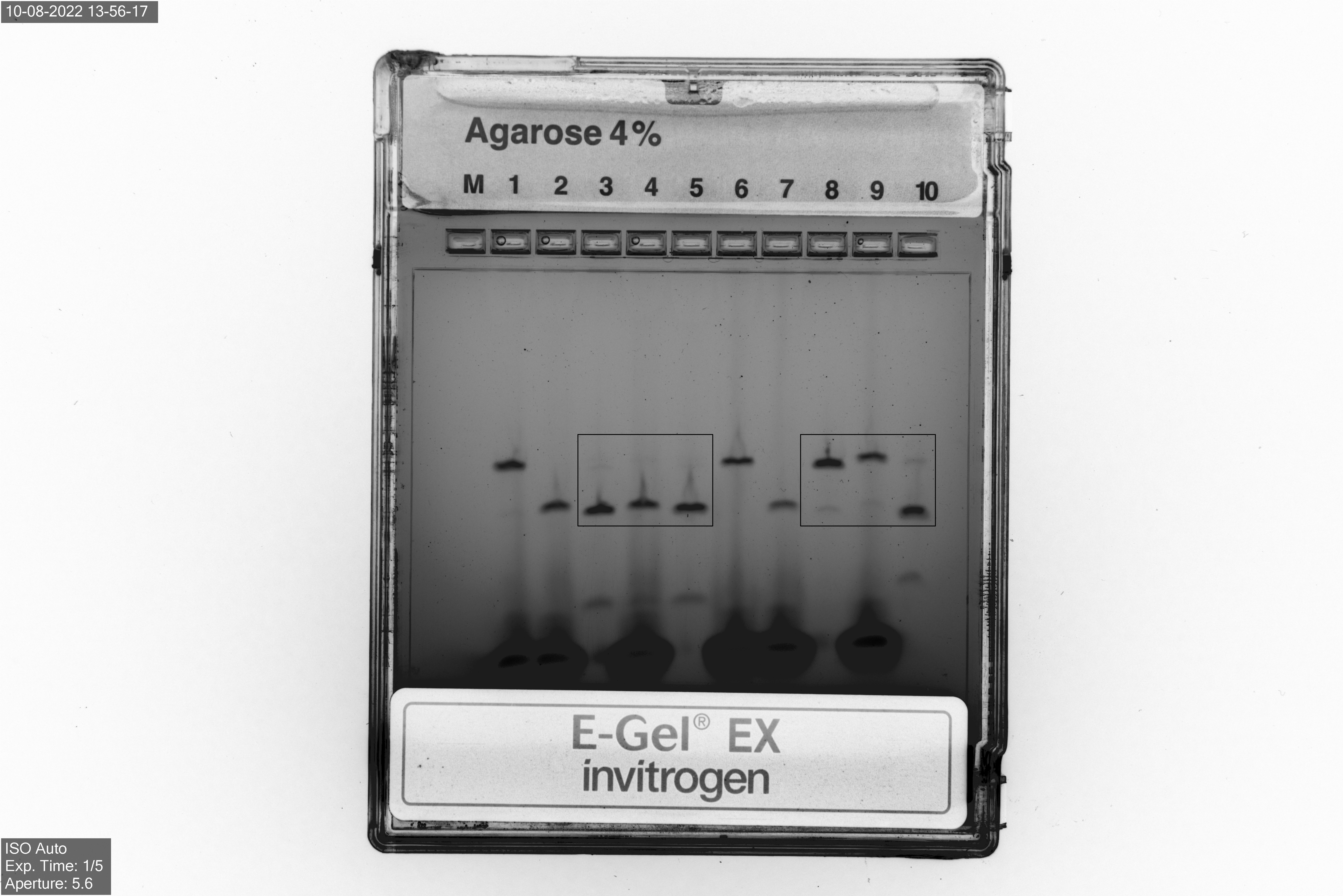

### Figure2_G3_G4_gel.png

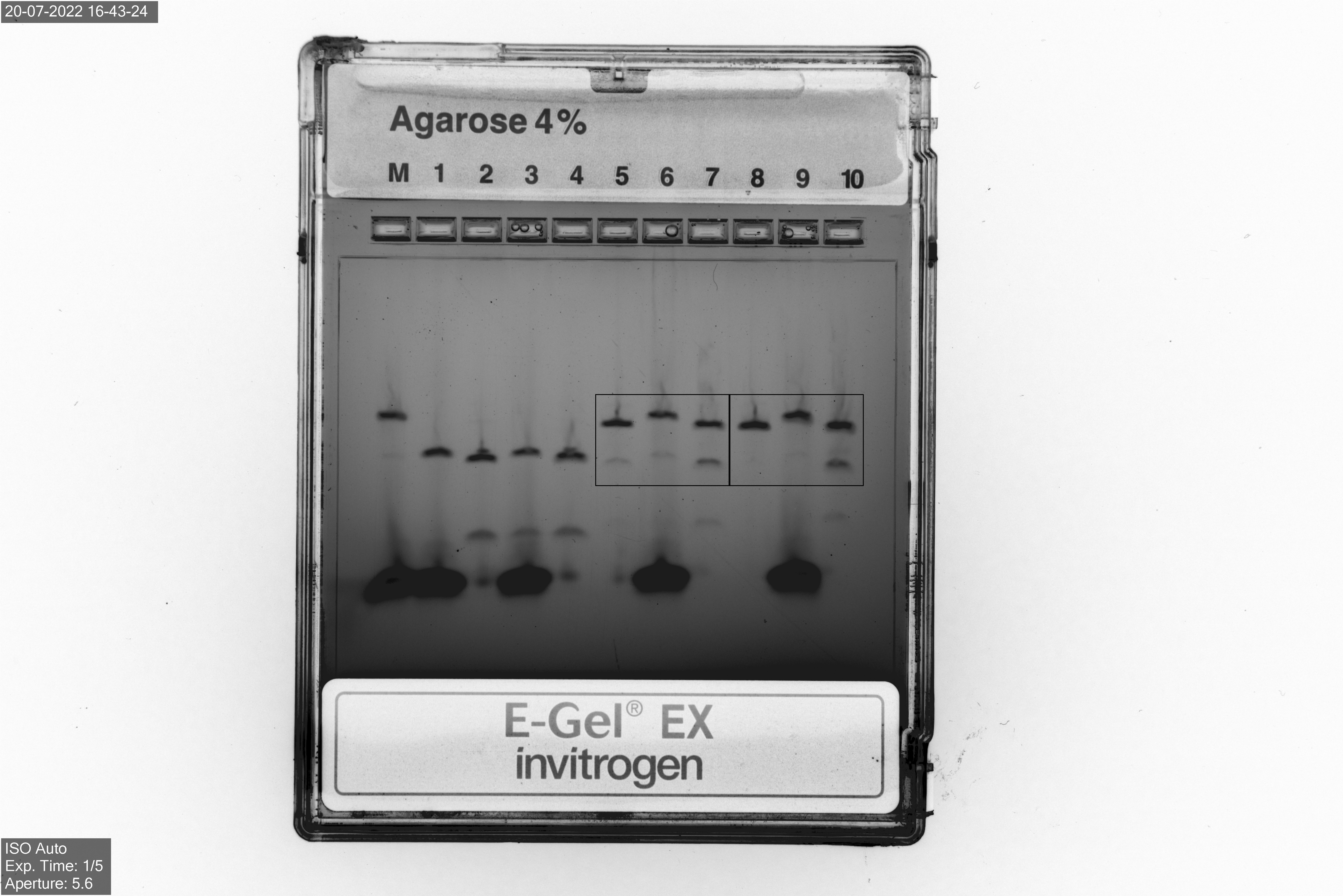

### Figure3_G1h_gel.png

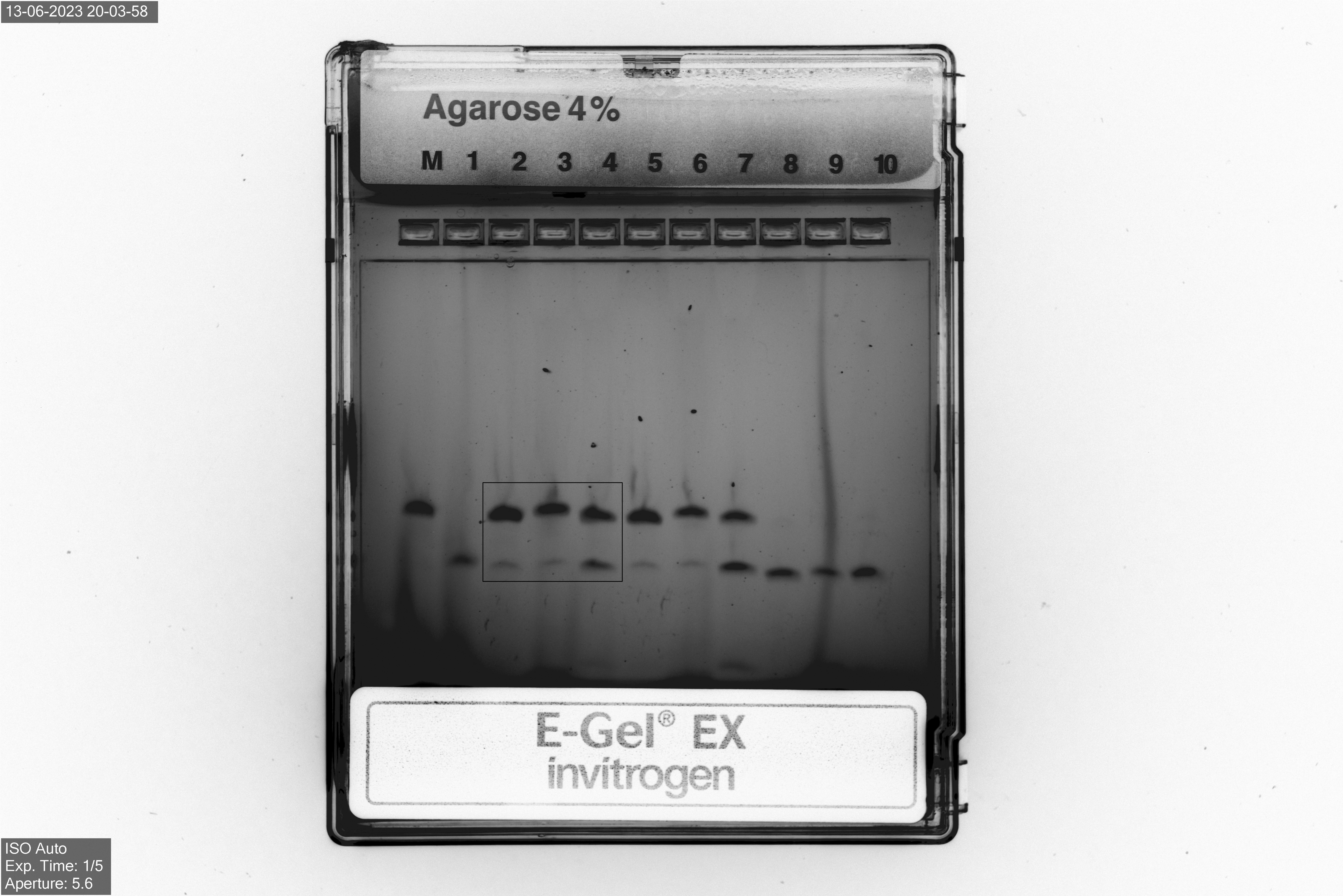

### Figure3_G2h_gel.png

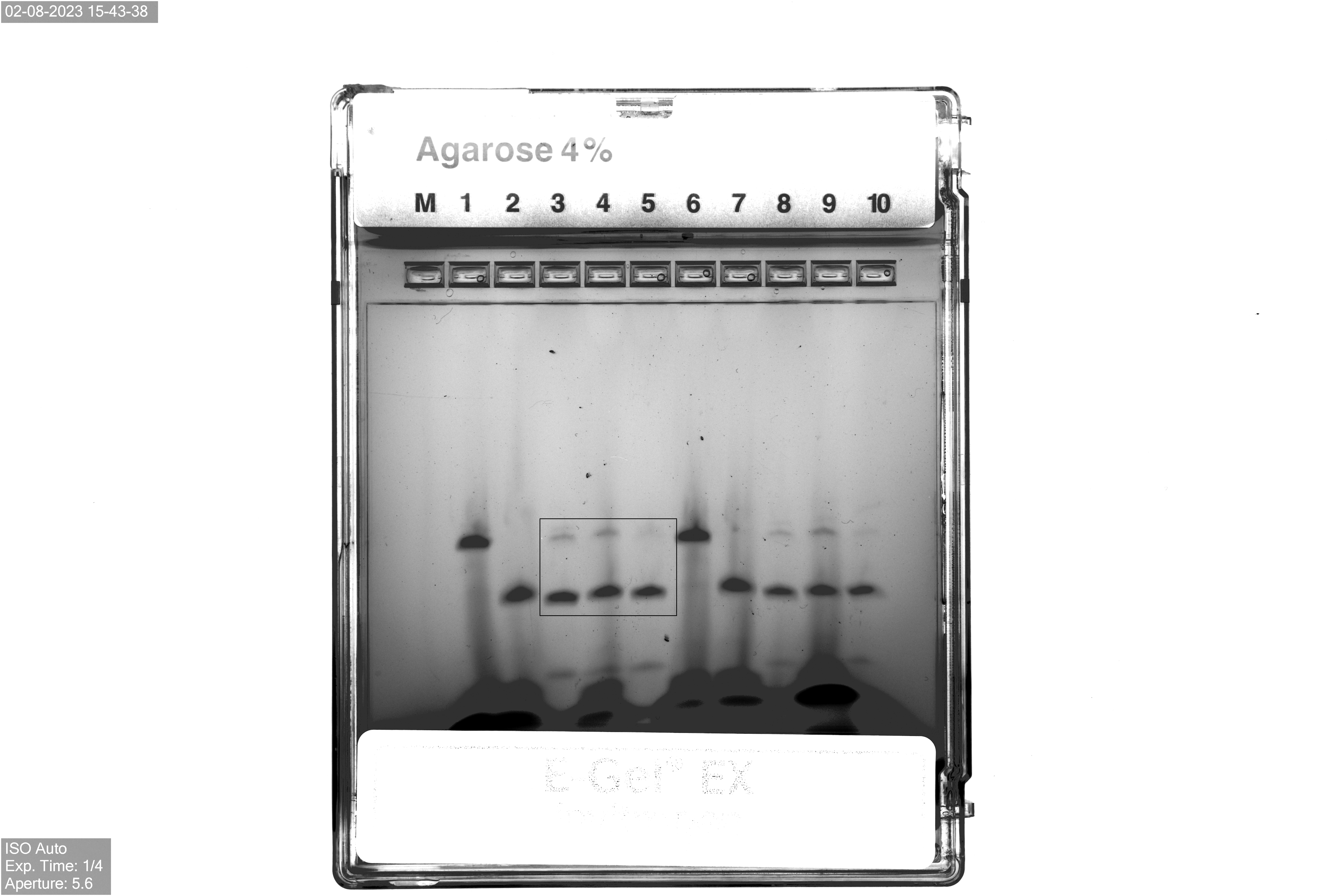

### Supp_Figure4_PCR.tiff

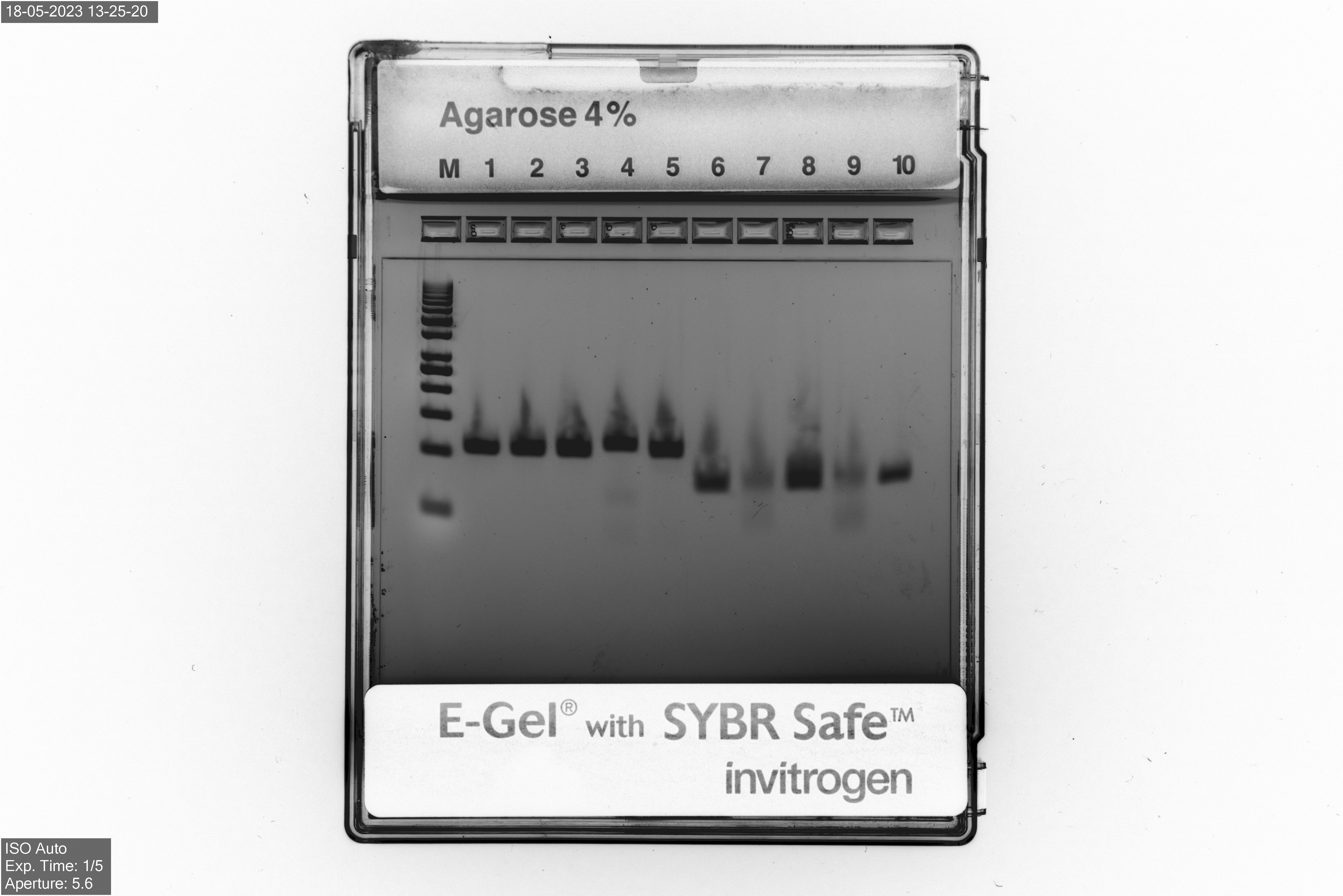

### Supp_Figure12_metrics.tiff

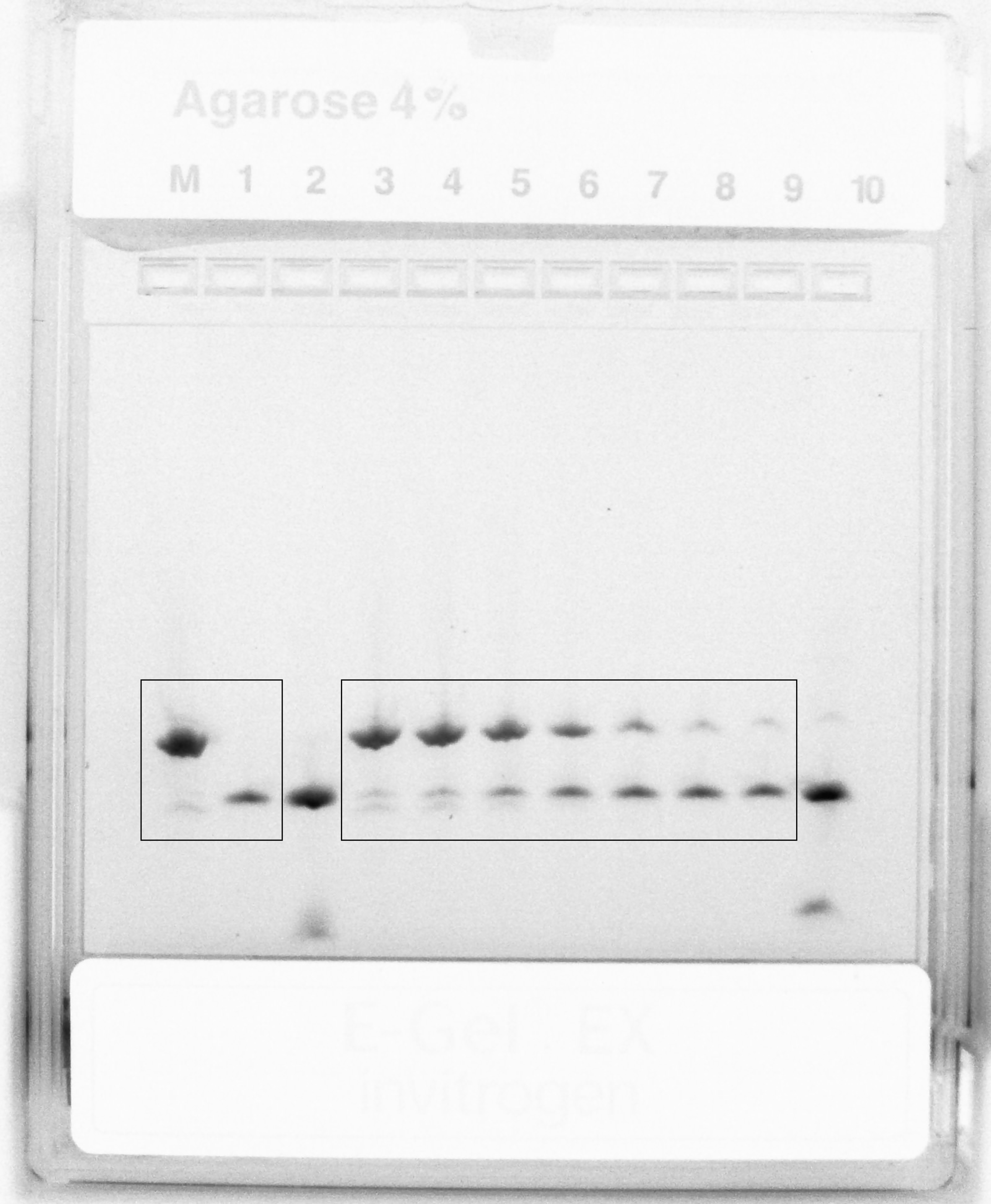

### Supp_Figure13_G2_G3_temps.tiff

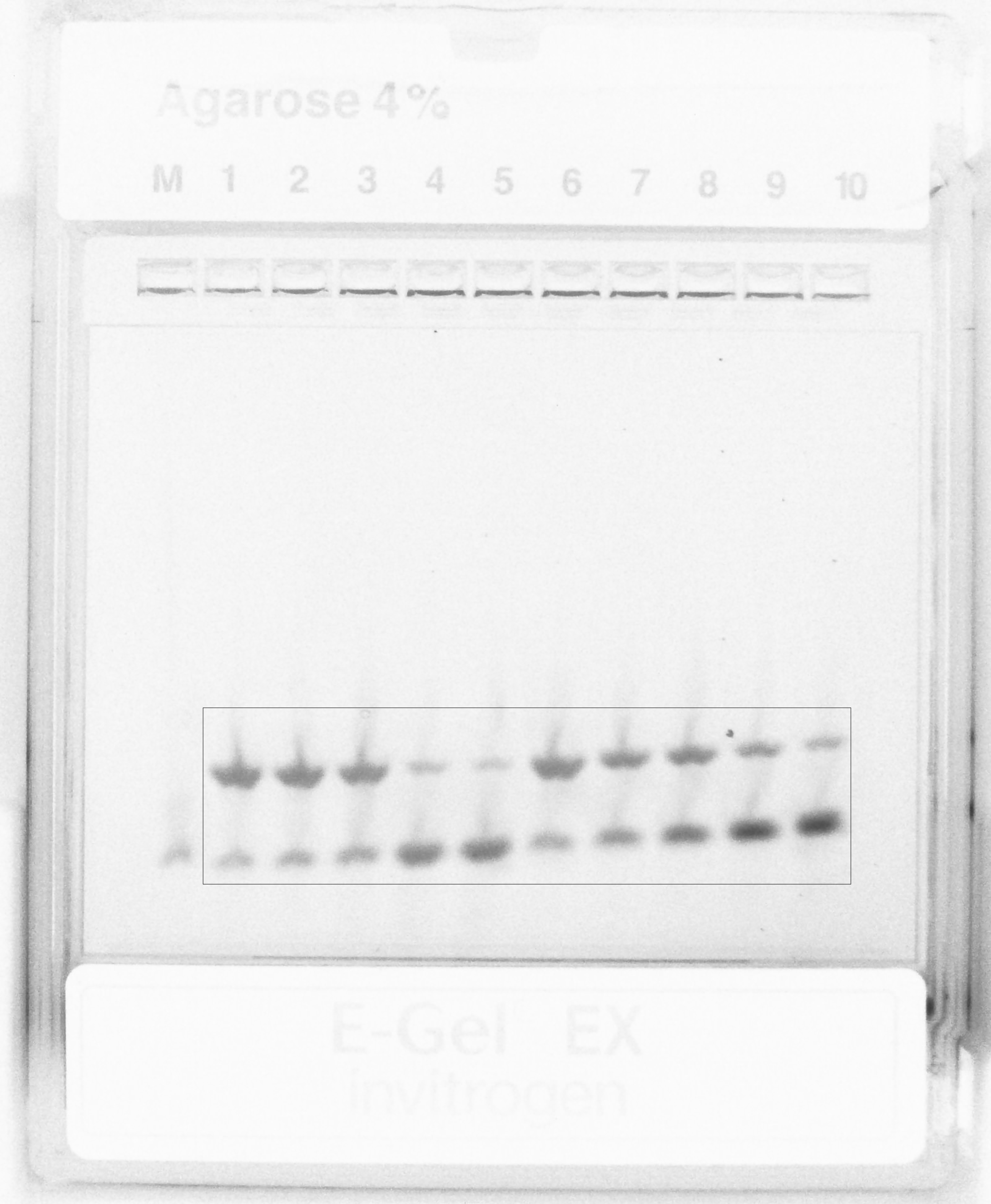

### Supp_Figure16_BO_titration.tiff

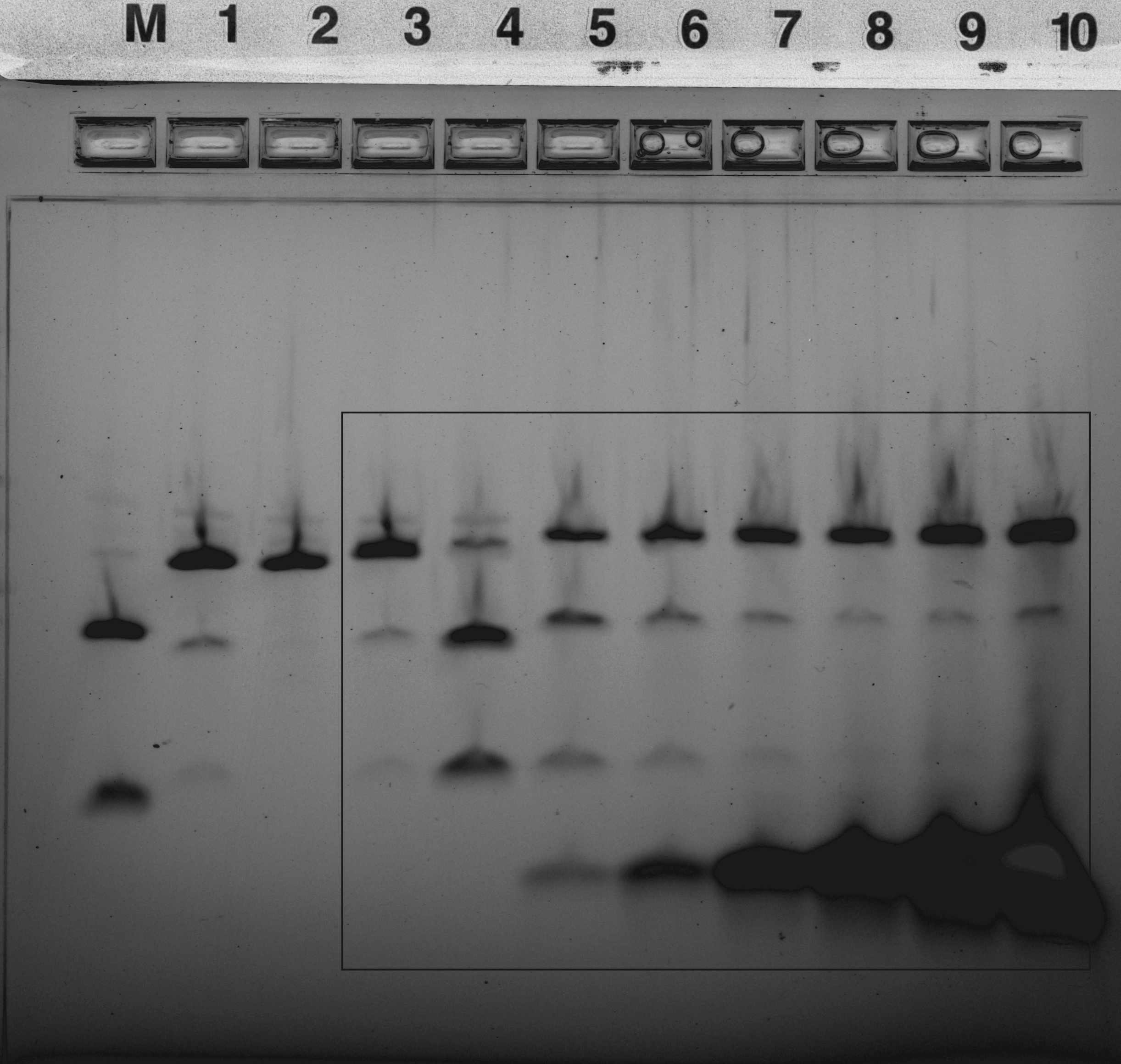

### Supp_Figure17_Temp37-50-60-65.tiff

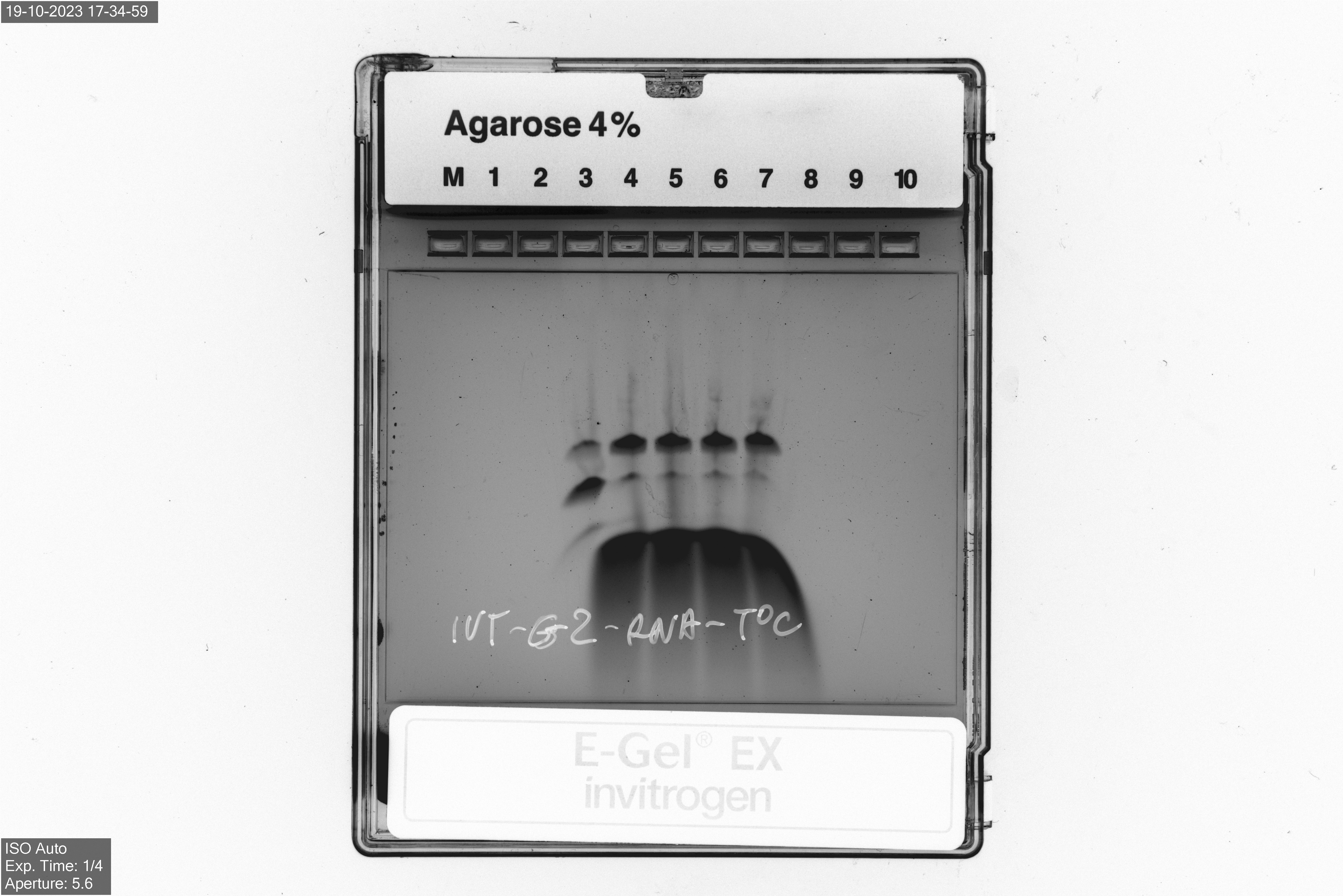
